# Invasion analysis for structured populations: analytical principles and numerical workflows

**DOI:** 10.64898/2026.11.04.763670

**Authors:** Joe Wan, Hsiang-Chih Lo, Guan-Yu Chen, Shiang-Chi Huang, Jia-Zhen Liu, Chin-Te Tsai, Po-Ju Ke

## Abstract

All real populations show some form of structure—differences among individuals, such as life stage or physiological condition, that shape a species’ ability to persist. Yet it remains challenging to incorporate population structure into models of species interactions. We provide a general framework for studying competition in structured populations by quantifying species’ invasion growth rates. The crux of our approach is separation of timescales: the initial dynamics of a structured population depend on how exactly it is introduced, but these transient differences do not affect long-term invasion success. Depending on the research question, the relevant long-term growth rate when rare may equivalently be derived using analytical separation of timescales, matrix population theory, or numerical simulations. Using case studies spanning developmental stage, environmental state, and infection status, we illustrate how our methodology derives simple, ecologically meaningful invasion conditions for a variety of systems. Finally, we extend the approach to fluctuating resident dynamics, showing how to use simulations or a general mathematical framework known as Floquet theory to correctly quantify invasion growth. By unifying the quantitative methods of population biology and community ecology, our framework will allow ecologists to study species persistence and coexistence across a greater variety of demographically complex systems.

## 1 Introduction

All living things change over time: whether through their own growth and development, or through interactions with their biotic and abiotic environment, individuals differ in ways that affect their success. Yet community ecology typically treats entire species as its fundamental unit. In other words, classic models of ecological communities represent each species’ population using a single variable (Nakazawa, 2015). This is in contrast to the field of population ecology, which has long recognized the importance of population structure—the distribution of individuals among different states such as sex, age, and size—for understanding population dynamics (e.g., De Roos & Persson, 2013). Such structure is not limited to individual development, as individuals may also differ in their physiological conditions (e.g., dormancy, infection status; Letten *et al*., 2024), spatial location (Nakazawa & Huang, 2016), or environmental state (Ke & Levine, 2021). Classic population-level theory highlights how differences in stage structure affect populations’ ability to persist (Caswell, 2001). Meanwhile, a growing perspective emphasizes that stage structure can also determine the outcome of species competition (Werner *et al*., 2025). Accordingly, stage structure plays an important yet understudied role in determining whether species can coexist—a long-standing question in community ecology.

Invasion analysis, the foundational tool for studying species’ persistence within a community, nonetheless remains challenging in structured populations. Ecologists quantify a species’ invasion growth rate (IGR) as its growth rate when it is at low density within the environment created by its competitors. Species with positive invasion growth rates are predicted to persist because they recover from low abundance and avoid competitive exclusion. Beyond predicting coexistence based on mutual invasibility (Barabás *et al*., 2018; Spaak & De Laender, 2020), invasion analysis is the first step for partitioning the mechanisms driving coexistence (Ellner *et al*., 2019; Shoemaker *et al*., 2020) and provides a fundamental way to quantify fitness in evolutionary biology (Yamamichi *et al*., 2022). However, the invasion growth rate is hard to define in structured populations: whether a rare population initially grows or shrinks depends strongly on which stages are dominant (Iles *et al*., 2016). Formal theory has established that the persistence of structured populations is determined by precise long-term growth metrics (e.g., their dominant Lyapunov exponents; Benaïm & Schreiber, 2009; Hofbauer & Schreiber, 2010; Roth & Schreiber, 2014). Nonetheless, these quantities are rarely derived in more system-specific ecological models. Instead, studies of coexistence in structured populations have more commonly relied on numerical simulations (Werner *et al*., 2025) or strong simplifying assumptions (DeMalach *et al*., 2021; Song *et al*., 2021), indicating that general analytical workflows for calculating invasion growth rates remain poorly established (but see Klausmeier, 2008). Taken together, these studies highlight the need for a generally applicable approach for invasion analysis in structured populations.

Structured populations may often present an additional challenge: complex dynamics such as population cycles or chaos may emerge due to delayed density dependence, reflecting stage-specific feedbacks in reproduction, maturation, and mortality (De Roos & Persson, 2013; De Roos *et al*., 2008; Gurney & Nisbet, 1985). These population fluctuations introduce two additional complications to invasion analysis. An invading species’ growth rate varies over time, while its initial success depends on the particular state of residents’ population dynamics it experiences at introduction. With the loss of a fixed resident equilibrium, analytical treatments quickly become mathematically cumbersome or intractable (Ellner *et al*., 2016; Klausmeier, 2008). In these cases, invasion analysis requires concrete numerical simulations to estimate asymptotic growth over the resident cycle while simultaneously accounting for the invader’s own internal population structure. Yet, to date, numerical approaches for structured invasion analysis remain scattered across specific models (Letten *et al*., 2024; Werner *et al*., 2025), with few overarching principles for estimating and validating invasion growth rates in time-varying structured populations.

Here, we develop a general framework for invasion analysis in structured populations, enabling researchers to study coexistence in diverse new systems and research questions. After briefly reviewing invasion analysis in classical models (Box 1), we use a simple age-structured competition model to illustrate how population structure complicates invasion analysis (section 2). Then we show how to address this challenge using two approaches. First, we demonstrate that the invader’s population structure has much faster dynamics than population growth, allowing us to calculate invasion growth rates using separation of timescales (section 3). Second, we show that invasion analysis in structured populations is mathematically closely related to matrix population models, and thus, that linear algebra–based tools for population ecology provide a succinct and general method to quantify invasion (section 4). We illustrate the generality of our framework through case studies in which population structure represents not only developmental stages, but also diverse factors such as disease status or spatially structured habitats (section 5). Finally, for systems that are not analytically tractable, we provide reproducible numerical recipes guided by the analytical insights for rigorously studying invasion dynamics in arbitrarily complicated systems (section 6). By making the connection between invasion analysis and structured-population theory explicit, we provide a coherent and accessible toolkit for studying species persistence and coexistence across a wider range of ecological systems.

### Box 1. Review: Invasion analysis in unstructured populationsBox 1. Review: Invasion analysis in unstructured populations

In this Box, we briefly summarize how classic invasion analysis can be used to study species coexistence in models without population structure. Consider a model of competition between *n* different species’ populations *N*_*i*_, where *i* = 1, …, *n*. In continuous time, we can model the dynamics of species *i* according to

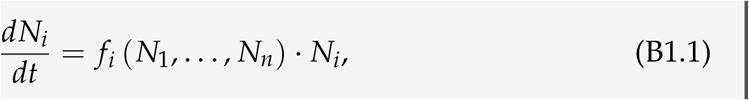

where *f*_*i*_ is the per-capita growth rate as a function of species abundances; for instance, *f*_*i*_ (*N*_1_, …, *N*_*n*_) = *r*_*i*_ *−* ∑_*j*_ *α*_*ij*_ *N*_*j*_ for the Lotka–Volterra model. We are interested in determining whether a species of interest will persist in the community after starting at some positive abundance. This species, which we will label 1, is known as the *invader*, while all other species are known as *residents*.

**Resident equilibrium** The first step is to consider the equilibrium reached by the residents when the invader is absent (i.e., *N*_1_ = 0). To do so, we find the abundances 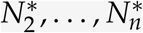 such that all residents experience zero growth, giving the system of equations

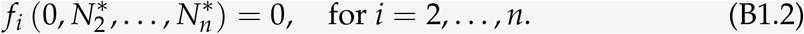

For instance, in a two-species Lotka–Volterra model, the resident equilibrium abundance of species 2 is obtained by solving 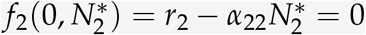, yielding 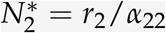.

**Invasion growth rate** Assuming the residents reach a stable and well-defined equilibrium, we can then evaluate the per-capita growth rate of species 1, the invader, when introduced at extremely low abundance (*N*_1_ *≈* 0) into the resident equilibrium. Crucially, under this assumption, known as the *invasion scenario*, the invader is too rare to affect the residents’ abundances or its own growth. Thus, at the beginning of invasion, it grows exponentially with a constant per-capita rate known as the *invasion growth rate*, given by

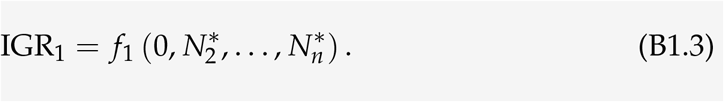

The invader persists when its invasion growth rate is positive, i.e., IGR_1_ > 0. Stated conceptually, by ensuring that the species recovers from low abundance, this condition ensures that the species will not be excluded from the community. In our two-species Lotka–Volterra example, the invasion growth rate of species 1 is IGR_1_ = *r*_1_ *− r*_2_*α*_12_/*α*_22_, and it persists if and only if (*r*_1_/*r*_2_) *·* (*α*_22_/*α*_12_) > 1.

**Coexistence conditions** By repeating the invasion analysis for all species, we can predict when the entire community can robustly coexist. Specifically, coexistence requires *mutual invasibility*: each species *i* must have positive invasion growth. In other words, all species persist because they can recover from low abundance.

## 2 Problem: Accounting for population structure in invasion analysis

Invasion analysis is an important technique routinely covered in theoretical ecology text-books and is relatively straightforward in simple theoretical models (Box 1). However, applying it in more realistic settings is complicated by the problem of *population structure*. Rather than considering a species’ total population *N*_*i*_, we might consider a series of discrete states 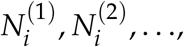, with superscripts representing our state(s) of interest (e.g., life stages, habitat, or disease status). In this section, we introduce how such models account for population structure, and illustrate why this complicates invasion analysis.

### Simple stage-structured competition model

As a concrete example of our general framework (Appendix S1), we begin with one of the simplest models where population structure matters. In particular, we consider the case where our invader of interest (species 1) has two life stages (juveniles 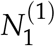 and adults 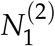), but its competitor (species 2) does not (represented simply by *N*_2_; Fig. 1a). Dynamics obey the following system of ordinary differential equations:

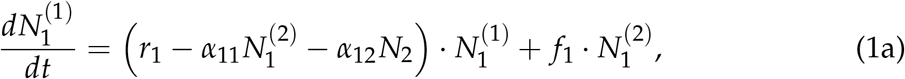

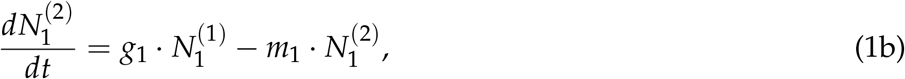

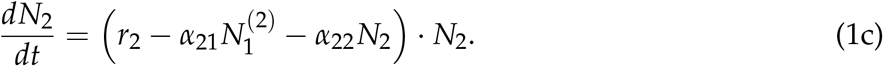

**Figure 1:**
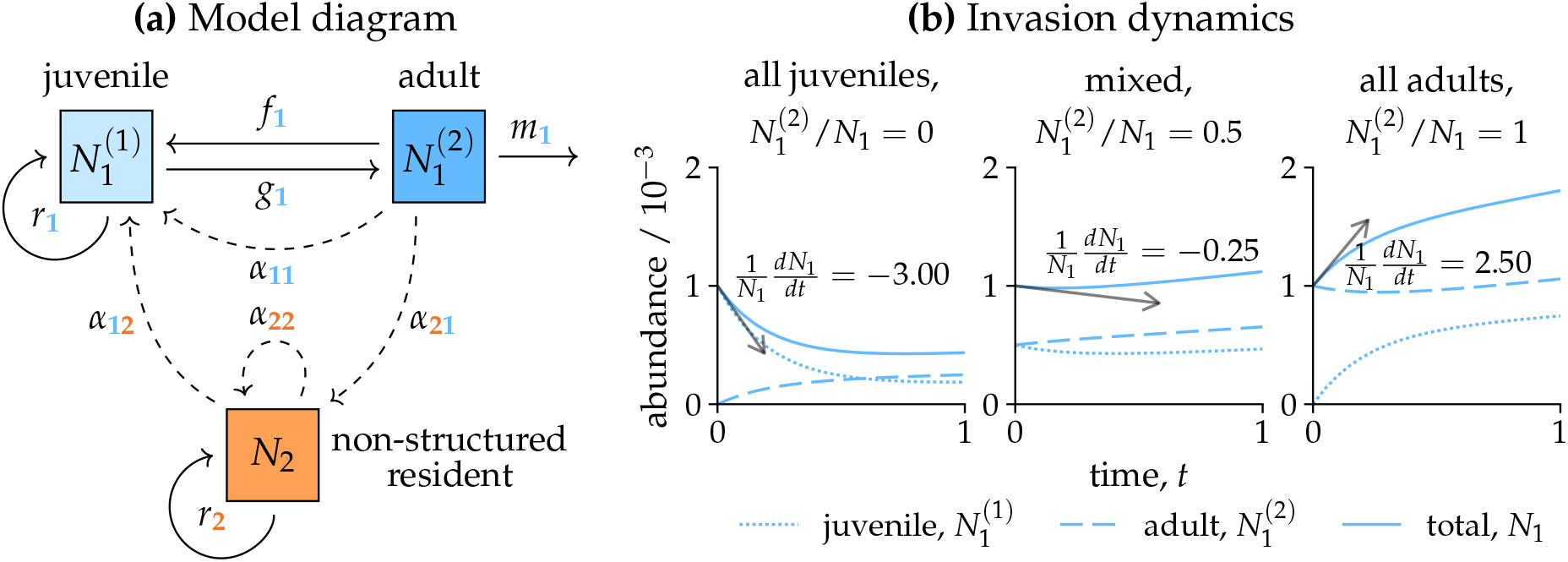
Simple model with stage structure. **(a)** Model diagram of a stage-structured species 1 competing against species 2 with no stage structure. Species 1 has two stages, juveniles 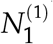 and adults 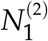. Solid arrows indicate population transitions: juveniles mature into adults at a rate *g*_1_ and replenish their own population at a rate *r*_1_, whereas adults produce new juveniles at a rate *f*_1_ and die at a rate *m*_1_. Dashed arrows indicate competition: juvenile replenishment rate *r*_1_ is linearly reduced by competitors of species *j* according to the competition coefficient *α*_*ij*_; we assume that all competitive effects due to species 1 are solely imposed by adults. **(b)** Instantaneous invasion dynamics under different initial population structures: 0% adults (left), 50% adults (middle), and 100% adults (right). In all cases, an initial total population of 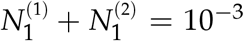 invades a resident population of species 2 at its equilibrium (not shown). We plot the abundance (*y*-axis) over time of species 1 (juveniles 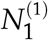 : dotted; adults 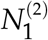 : dashed; total *N*_1_: solid). The initial rate of change of the total population (gray tangent arrow) differs between panels. Parameter values: *r*_1_ = *−*1, *f*_1_ = 3, *g*_1_ = 1, *m*_1_ = 0.5, *r*_2_ = 1, *α*_11_ = 5, *α*_12_ = 3, *α*_21_ = 0.5, *α*_22_ = 1.

Here, *N*_2_ follows the standard Lotka–Volterra dynamics considered in Box 1, but species 1 experiences additional complexity due to transitions between its two states (Fig. 1a, solid arrows): juveniles 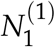 replenish their own population at a rate *r*_1_ and mature into adults 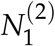 at a rate *g*_1_, while adults produce new juveniles at a rate *f*_1_ and die at a rate *m*_1_. Note that the juvenile-to-juvenile replenishment rate *r*_1_ reflects the balance of juvenile reproduction minus mortality and maturation, and thus may be positive or negative. The species also experience competition: competitor density negatively affects each species *i*’s rate *r*_*i*_, representing the juvenile-to-juvenile transition for species 1 and general population growth for species 2. We assume this effect is mediated by a linear competition coefficient *α*_*ij*_, where *j* is the competitor species, and that all competitive effects due to species 1 are solely mediated by adults 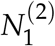 (Fig. 1a, dashed lines). For the general analysis, we assume all variables are positive except *r*_1_, which is negative due to mortality and maturation (specific parameter values are given in Fig. 1).

### Invasion dynamics depend on population structure

To study whether species 1 can persist when facing species 2, we consider its invasion scenario: species 2 is at its equilibrium abundance 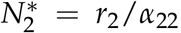, as calculated in Box 1, while species 1 is introduced at low total abundance (i.e., 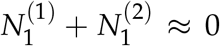). A naive approach might be to simply introduce one of the stages of species 1 at low abundance and calculate the instantaneous per-capita growth rate. For instance, introducing juveniles alone, we would find the per-capita growth rate of the population (*dN*_1_/*dt*)/*N*_1_ approaches 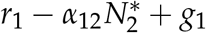 when the initial abundance is near zero. For adults, we would instead find that (*dN*_1_/*dt*)/*N*_1_ = *f*_1_ *− m*_1_. If we interpreted these outcomes as evidence of invasion success, our conclusion would change depending on the invader’s initial stage distribution (Fig. 1b): introducing only juveniles would suggest that species 1 cannot persist, whereas introducing only adults would suggest successful invasion. How can ecologists obtain consistent, and more importantly, biologically meaningful predictions about invasion success in structured populations?

## 3 Method 1: Taking advantage of fast population structure dynamics

Fortunately, there is a conceptually simple solution: during invasion, the transient dynamics of population structure can be separated from the long-term growth of invader abundance. Under the invasion scenario, the invader remains sufficiently rare that it has negligible effects on the resident community and its own density-dependent demographic rates. Thus, the invader experiences an effectively fixed resident environment, and the effects of its initial stage distribution disappear before the overall population grows to appreciable abundance. We illustrate this by simulating the invasion of species 1, with various initial stage distributions, into a resident population of species 2 in the example model (dark to light blue in Fig. 2). The proportion of adults converges within a few time units regardless of the initial stage distribution (left of the *x*-axis break; Fig. 2a). Accordingly, there is a well-defined invasion growth rate that predicts invasion success: as shown in Fig. 2b, species 1’s per-capita growth rate remains stable (top panel) while the population remains rare (bottom panel). Thus, invasion success should be evaluated using the per-capita growth rate after population structure has settled, rather than the instantaneous growth rate associated with any particular initial stage distribution. In this section, we use this insight to derive analytical expressions for invasion growth rates in structured populations using the technique of timescale separation (Otto & Day, 2011). We present two variants of this approach: a direct method applicable to any model, and a mathematical transformation technique which greatly simplifies calculations in many cases (see Appendix S2 for detailed derivations).

**Figure 2:**
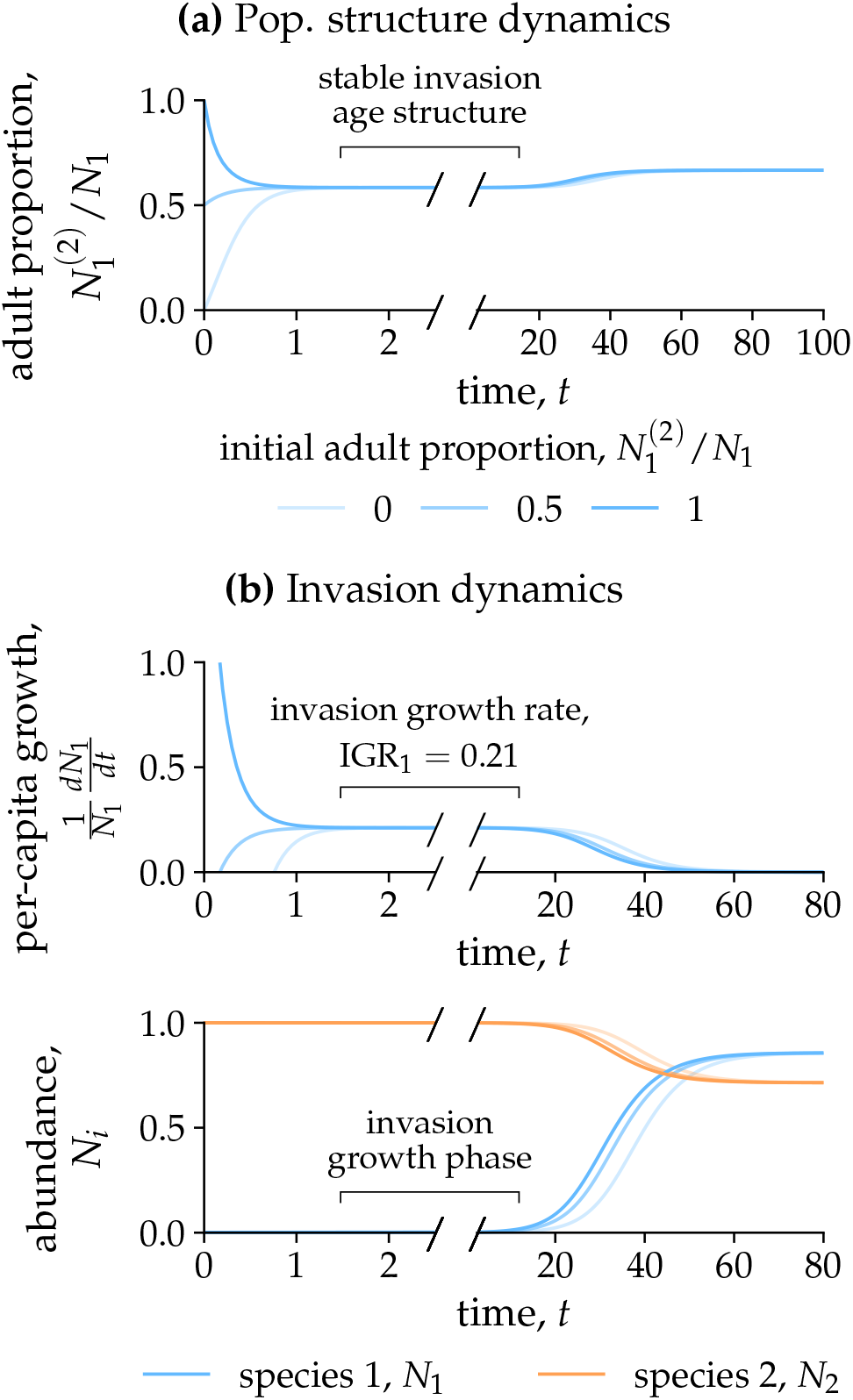
Fast and slow dynamics under invasion. We plot the temporal trajectory of **(a)** population structure (the proportion of adults within the entire population) and **(b)** per-capita growth rate for the invader population (top) and the total species abundance (bottom). Parameters are as in Figure 1 with an initial population 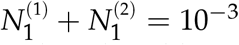. Different choices of initial population structure are shown as differently colored lines. Note that the resolution differs before (i.e., during the initial invasion) and after (i.e., subsequently approaching equilibrium) the break in the axis (after *t* = 2). The period when the invader reaches a stable population structure while remaining sufficiently rare gives the relevant invasion growth rate (IGR = 0.21; eq. 4).

### Method 1.1: Direct analytical timescale separation

Under timescale separation, we can consider population structure as a fast variable, and overall population growth as a slow variable. To do so, we transform the model into two new variables representing each process: let *Q* represent population structure and *N* represent overall population size. There are many ways to capture these two aspects, but a mathematically simple choice is simply to use the juvenile-to-adult ratio and the adult abundance: 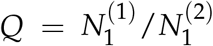 and 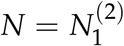 (see Appendix S2 for other choices). Using the chain rule, we can rewrite the dynamics of species 1 in terms of these new variables:

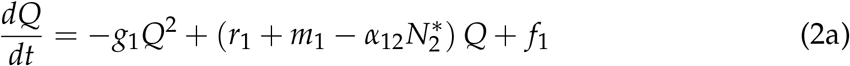

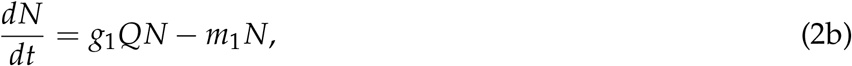

where we have dropped the 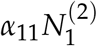 term because it is near zero during invasion. Next, to perform timescale separation, we assume that *Q* rapidly approaches a steady state known as the *quasi-equilibrium Q*\* for a given near-constant value of the slow variable *N*. Setting *dQ*/*dt* = 0 in eq. 2a, we can solve a quadratic equation to find that the sole positive root is

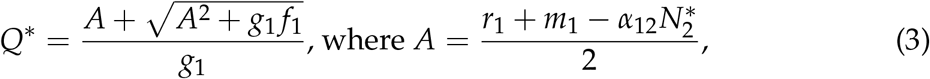

and substituting this value of *Q*\* into eq. 2b gives the invasion growth rate as

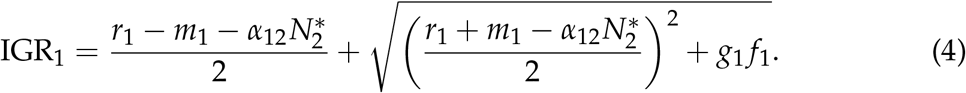

Inserting the particular parameter values used in Fig. 1, we find an invasion growth rate IGR_1_ = 0.21 matching the simulation result (Fig. 2b). This is no surprise, as direct simulation applies the same idea of a quasi-equilibrium stage distribution: we identify the time at which different starting states converge on a stable stage structure and evaluate the per-capita growth rate there (see general method in Appendix S3, Algorithm S3.1). Finally, we solve for the conditions under which IGR_1_ > 0 to find a simple condition for successful invasion by species 1:

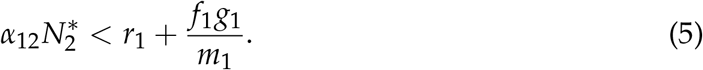

In other words, species 1 can invade whenever the negative effect of the resident species 2 is outweighed by the sum of species 1’s population growth from juveniles (*r*_1_) and from adults (*f*_1_), after correcting for the effects of maturation and mortality (*g*_1_/*m*_1_). This method can be applied to any ordinary differential equation model with population structure, and relies only on separation of timescales, a technique routinely covered in introductory curricula on theoretical ecology (e.g., Otto & Day, 2011). Thus, we emphasize that invasion analysis in structured populations need not be mathematically challenging.

### Method 1.2: Strategic linear combination

Though the final invasion condition in eq. 5 is simple, the intermediate calculations may require cumbersome algebra. However, knowing that stage structure rapidly approaches a quasi-equilibrium, we can derive conditions for invasion in a much simpler way. Consider defining a new variable as some linear combination of the juvenile and adult abundances, 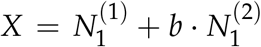, where *b* is a positive constant. By linearity, 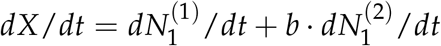. The key insight is that we can strategically choose *b* such that one stage variable is eliminated from *dX*/*dt*. As one option, we can choose *b* = *f*_1_/*m*_1_, which causes the *−m*_1_ term in the adults’ dynamics to cancel with the *f*_1_ term in the juveniles’ dynamics, leaving

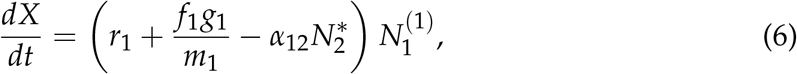

where we have again dropped the 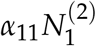 term due to the invasion scenario. Further-more, thanks to the constant population structure at quasi-equilibrium, the remaining stage 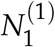 is a positive constant multiple of *X*: substituting 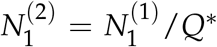 into the definition of *X* gives 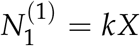, where 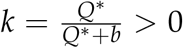. Dividing eq. 6 by *X* and using the above expression, we obtain the invasion growth rate

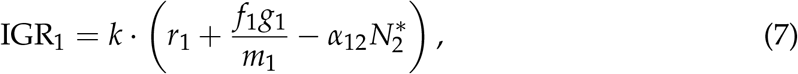

which is equivalent to eq. 4 after multiplying out the full expression for *k*. However, since *k* is always positive, it never affects the sign of the invasion growth rate, and we can immediately see that the term within the parentheses leads to the same invasion condition we found earlier in eq. 5. In many models, a suitable positive linear combination can eliminate one or more stage variables from *dX*/*dt*, allowing the sign of the invasion growth rate to be determined without explicitly solving for the quasi-equilibrium stage distribution.

#### Box 2. Review: Population growth in matrix population models

In this Box, we review classic results for a family of models from population ecology known as matrix population models. In continuous time, the abundance of a single structured population can be represented by a vector 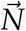, whose elements give the abundances in each state. When transition rates are constant, the population dynamics obey the vector differential equation

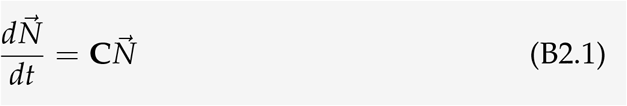

where **C** is a constant *transition matrix* representing changes between states. Note that a similar analysis applies to the discrete-time analogue of this system, i.e., the classic Leslie and Lefkovitch models (Appendix S1).

**Solving long-term dynamics using linear algebra** A key result from matrix population theory is that differences in the initial population structure only affect the transient dynamics. After this transient phase, the relative abundances of the states approach a stable structure, and the whole population grows or declines exponentially with a common rate *λ*, the *long-term population growth rate*:

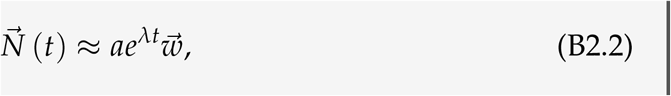

where *a* is an arbitrary constant and 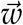 gives the relative proportions of different states, i.e., the *long-term state structure*. Basic tools from linear algebra allow us to solve for the long-term growth rate *λ* and the long-term state structure 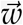. Specifically, *λ* and 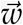 are mathematical properties of the transition matrix **C**, obtained as solutions to the equation

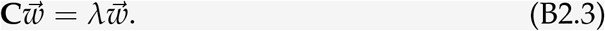

Mathematically, 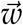 is an *eigenvector*, a vector whose elements’ relative magnitudes are unchanged by matrix multiplication, but rather scaled by a scalar factor *λ*, known as the *eigenvalue*. For an *m × m* transition matrix **C**, there are up to *m* distinct eigenvalues. The long-term growth rate of the population is given by the *leading eigenvalue*, i.e., the *λ*_*i*_ with the greatest real part, and the long-term population structure by the corresponding eigenvector 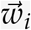. On the other hand, non-leading eigenvalues govern transient changes in population structure. Thus, eigenvalues provide a precise mathematical description of the timescale separation between fast changes in population structure and the long-term growth rate defining invasion success.

## 4 Method 2: Applying classic theory from matrix population models

The conceptual insight above, that population structure has rapid dynamics during invasion, is closely related to classic results from the theory of matrix population models (Box 2). This connection provides a succinct and mathematically complete way to quantify invasion growth rates in structured populations. To see this, we first rewrite the dynamics of species 1 in matrix form as

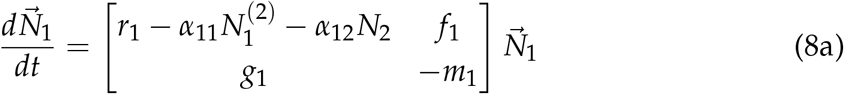

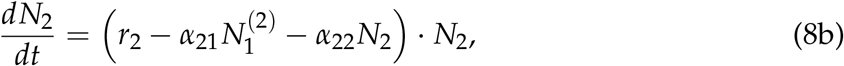

where the matrix in the first equation is the transition matrix specifying changes between and within the two states of species 1. As we have discussed above, under the invasion scenario, species 1 is rare and species 2 is at its equilibrium abundance 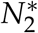 ; thus, as it invades, species 1 grows according to the constant transition matrix **C**,

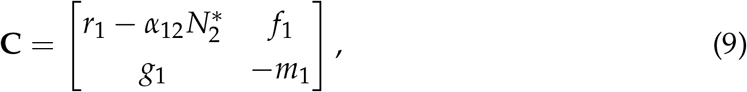

such that it obeys the linear vector differential equation 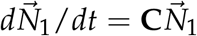. Mathematically, this is the invader’s Jacobian matrix evaluated at the resident equilibrium. Biologically, this matrix formulation shows that the rare invader behaves as a structured population growing in the fixed environment created by the resident. The eigenvalue perspective in Box 2 formalizes the timescale separation described above (Fig. 2): non-leading eigen-values capture transient changes in stage composition, whereas the leading eigenvalue provides the invasion growth rate after the effects of arbitrary initial stage distribution have disappeared. This interpretation holds as long as the invader remains sufficiently rare for its dynamics to be governed by the constant **C**. Accordingly, we can directly apply the results from Box 2 to quantify invasion growth rates in structured populations. We present two methods; as before, the first directly solves for the invasion growth rate (i.e., the leading eigenvalue), while the second derives conditions for invasion without explicitly solving for the eigenvalues.

### Method 2.1: Direct solution for leading eigenvalue

Following Box 2, after transient effects of the initial stage distribution have decayed, the long-term growth rate of the invading species 1 is given by the leading eigenvalue *λ*_1_ of the transition matrix **C**. The invader population grows exponentially at this rate, with a stable stage distribution given by the corresponding eigenvector. We can find the eigenvalues by solving the equation det(**C** *− λ***I**) = 0, giving the characteristic equation

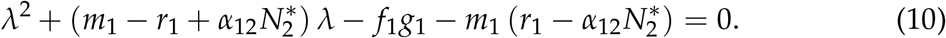

Solving this quadratic equation, we find that the dominant eigenvalue *λ*_1_, and hence the invasion growth rate IGR_1_, is

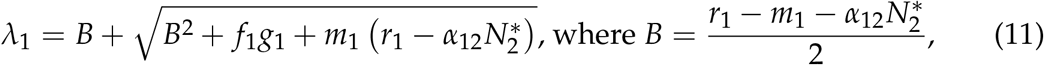

where we have taken the greater of the two roots. We can verify that this is equal to eq. 4 derived using timescale separation, confirming that this method recovers the correct invasion growth rate.

### Method 2.2: Applying stability conditions

For three or more stages, analytical expressions for eigenvalues can be biologically uninterpretable or may need to be solved numerically, limiting the usefulness of the above approach (Iritani & Day, 2026). Here, as in Method 1.2, we show that the sign of the leading eigenvalue can be determined without directly solving for its analytical expression, which greatly simplifies the algebra while providing the necessary information to predict invasion success. To do so, we apply the Routh–Hurwitz stability criteria (Appendix S1). For 2 *×* 2 matrices such as the one we consider here, the criteria reduce to two simple conditions: a system is stable—that is, all eigenvalues have negative real parts—if the matrix has a positive determinant det(**C**) > 0 and a negative trace (i.e., the sum of diagonal elements) tr(**C**) < 0. Thus, the resident equilibrium is unstable to invasion, and the invasion growth rate is positive, when at least one of the following conditions is true: (1) the determinant is negative, det(**C**) < 0, or (2) the trace is positive, tr(**C**) > 0, excluding boundary cases where each expression equals exactly zero. Applying this condition to **C** from eq. 9, we find that the determinant is det 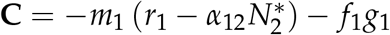, and the condition for it to be negative exactly matches the invasion condition derived using earlier approaches (eq. 5). On the other hand, the trace is 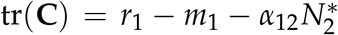, which is positive when 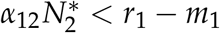. Because *m*_1_, *f*_1_, and *g*_1_ are positive, we note that the trace condition always implies the determinant condition, making the latter necessary and sufficient for positive invasion growth. Together, these complementary approaches show that structured invasion analysis can often yield concise, interpretable invasion criteria without requiring explicit solutions for the invader’s state distribution.

## 5 Case studies: Population structure in diverse systems

Before turning to systems requiring fully numerical analysis, we illustrate how the analytical approaches developed above can be applied to answer questions from diverse systems and subfields of community ecology. Across our three case studies, population structure represents developmental stage, environmental conditions, and infection status, demonstrating that structured invasion analysis is not limited to age- or stage-structured life cycles. For each case, we derive an analytical invasion criterion and use it to identify how key biological parameters determine invasion success, thereby showing how structured invasion analysis can translate complex state-dependent dynamics into biologically interpretable results.

### Case study 1: Stage-specific intraguild predation

Stage-dependent interactions provide a classic example of how population structure can modify species invasion. In some insect species, for instance, individuals of later developmental stages are aggressive cannibalistic herbivores that consume younger conspecifics and other competing herbivores, in addition to their original plant resource (i.e., intraguild predation *sensu* Polis *et al*., 1989). Here, we ask how aggression by older instars affects an aggressive herbivore species’ ability to invade a resident competing herbivore population. We considered a model where the older juveniles of species 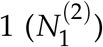 consume younger conspecifics 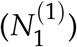 and younger juveniles of the competitor 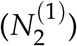. Both species’ juveniles consume resource *R* to mature into the next stage, whereas adults (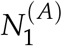 and 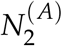) reproduce without relying on consuming *R*. We studied how four key parameters affect the invasion growth rates of the aggressive species (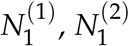, and 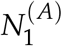; blue shaded box in Fig. 3a): the level of intraguild predation (*a*_21_) and its effect on the older instar’s mortality (*b*_21_), as well as the cannibalism level (*a*_11_) and its effect (*b*_11_). Together, these parameters allow us to distinguish how often aggressive interactions occur from how strongly they affect the demographic performance of the aggressive species.

**Figure 3:**
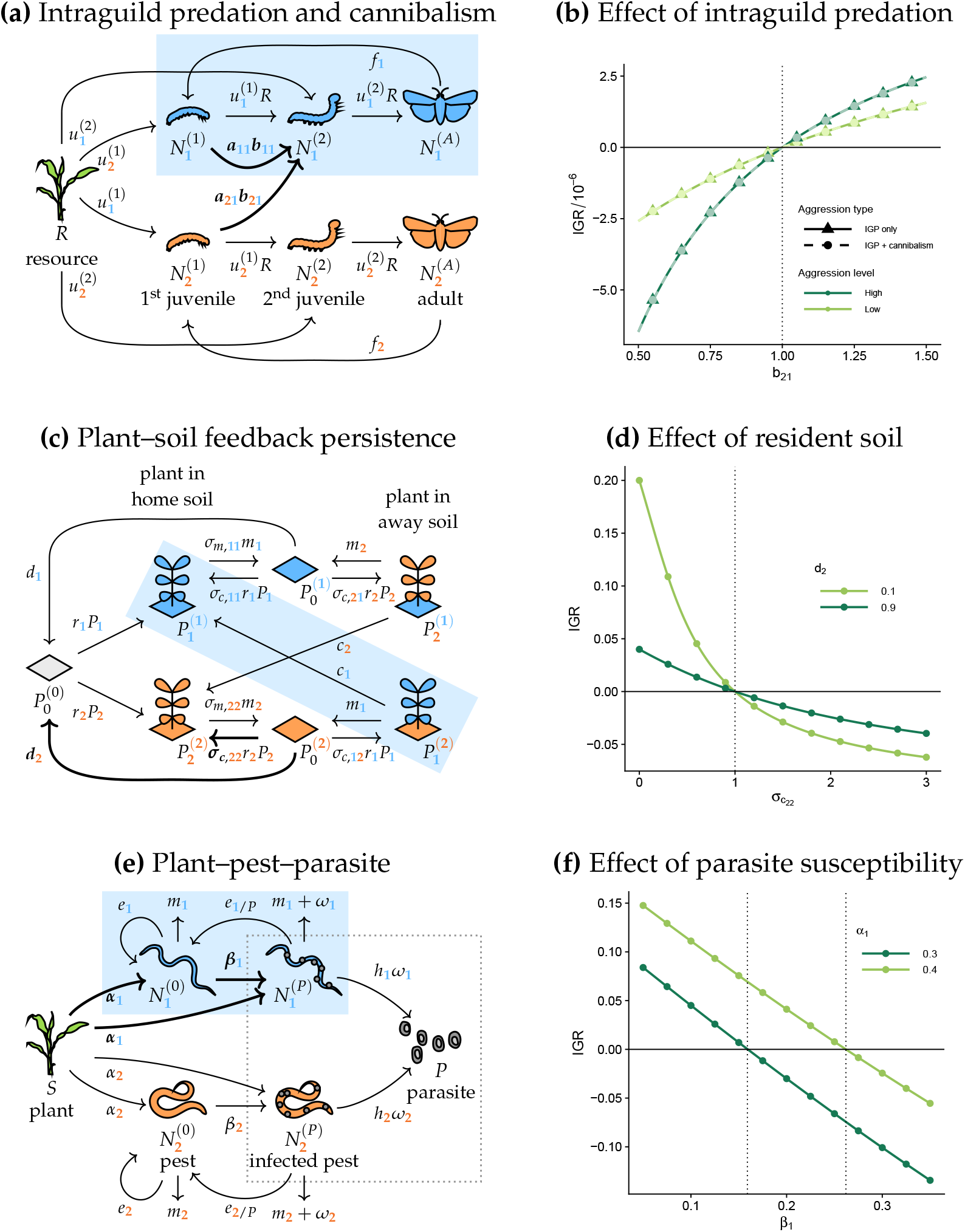
Applying invasion analysis to biologically distinct forms of population structure. The structured-population models include: **(a & b)** herbivores with developmental stage-dependent cannibalism and intraguild predation, for which we examine the effect of intraguild predation level (*a*_21_) and its demographic impact (*b*_21_); **(c & d)** a plant population structured across patches with different microbial legacies, for which we examine the strength of resident soil legacy (*σ*_*c*,22_) and its persistence (*d*_2_); and **(e & f)** pest populations structured by infection status in a plant–pest–parasite food chain, for which we examine the effects of the invading pest’s plant consumption rate (*α*_1_) and its susceptibility to the parasite (*β*_1_). In each model diagram (left panels), blue shading highlights the structured population, while focal parameters are indicated in boldface and by thick arrows; blue and orange colors indicate different species. Right panels show the corresponding effects of these parameters on invasion growth rate: solid lines are analytical results and points are numerical estimates (Appendix S3, Algorithm S3.1). Vertical dotted lines show analytical invasion boundaries and colors distinguish values of the second focal parameter. See Appendices S4–S6 for detailed analytical derivations. at which that interaction occurs: while intraguild predation level (*a*_21_) influences the magnitude of the invasion growth rate, its sign is determined solely by *b*_21_ (Fig. 3b). Moreover, aggressive behavior in the form of cannibalism does not affect the invasion outcome (overlapping circles and triangles in Fig. 3b). Although cannibalism may affect the equilibrium abundance once species 1 becomes common, it is biologically intuitive that cannibalism has little influence during invasion because encounters among conspecifics are negligible for the rare invader.

We derived the analytical invasion condition using all four methods described above. To isolate the effect of aggression, we assumed that the invader and the resident species are otherwise identical, so that all population differences between the two species are caused by aggression. We also assumed that adult-stage dynamics are fast compared with juvenile dynamics, allowing us to eliminate the adult stage and simplify the species to a two-stage population structure.

Under these assumptions, the invasion boundary for the aggressive species occurs simply at *b*_21_ = 1 (Appendix S4). Species 1 can invade only when consuming resident intraguild prey benefits its demographic performance (*b*_21_ > 1), whereas invasion fails when intraguild predation has a negative impact (*b*_21_ < 1). Our invasion analysis also separates the demographic consequence of a stage-specific interaction from the rate

### Case study 2: Resident soil legacy and rare-species invasion

Environmental states provide a less conventional example of population structure, in which individuals are structured by the patch types they occupy. Plant–soil feedback considers populations structured in this way: plants modify soil microbial communities, and these microbial effects can persist after plant death as a distinct soil state affecting later-arriving plants (Ke & Levine, 2021). Here, we ask how the persistence and strength of resident-conditioned soil determine the ability of another plant species to invade its competitor’s resident equilibrium. To address this question, we started from the fast-plant-conditioning patch occupancy model of Ke & Levine (2021), but extended it to a seven-variable model by allowing slow reconditioning when a plant grows on soil previously conditioned by another species. Specifically, state variables represent unoccupied unconditioned patches 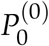, unoccupied patches conditioned by plant species 1 or 2 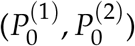, and occupied patches in which plant *i* grows on soil conditioned by plant 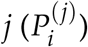. During plant 1’s invasion, it may grow on its own conditioned soil or on resident-conditioned soil, thereby being structured across different soil patches (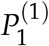 and 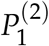; blue shade in Fig. 3c). We focused on two key parameters: *σ*_*c*,22_, the microbial effect modifying the resident plant 2’s colonization on its own conditioned soil, and *d*_2_, the decay rate of resident-conditioned soil. Together, *σ*_*c*,22_ and *d*_2_ allow us to examine how the strength and persistence of resident soil legacy shape the conditions for rare-species invasion.

Considering plant 1 invading the resident equilibrium of plant 2, we derived the analytical invasion condition using the methods described above (except for Method 2.1; Appendix S5). To isolate resident microbial effects on colonization, we set *σ*_*m,ii*_ = *σ*_*m,ij*_ and *σ*_*f* ,*ii*_ = *σ*_*f* ,*ij*_, assuming that conspecific and heterospecific soils have identical effects on a species’ mortality and fecundity. We also assumed that the two species have identical values of *m*_*i*_*σ*_*m,ii*_/*r*_*i*_*σ*_*f* ,*ii*_, thereby further removing species differences in microbe-mediated mortality and fecundity. Under these parameter assumptions, the remaining microbial effects are those on colonization, represented by *σ*_*c,ii*_; values greater or less than one indicate that soil microbes enhance or reduce the resident’s recolonization, respectively.

Under our simplified parameter assumptions, the analytical invasion criterion reduces to *σ*_*c*,12_ > *σ*_*c*,22_, a classic result suggesting that the resident must limit its own growth more than it limits the invader (Appendix S5). By setting *σ*_*c*,12_ = 1, the invasion boundary occurs at *σ*_*c*,22_ = 1 (Fig. 3d). Plant 1 can invade only when resident-conditioned soil weakens the resident’s ability to recolonize its own soil (*σ*_*c*,22_ < 1), and invasion growth rate increases when this soil legacy decays more slowly after host death (smaller *d*_2_; Fig. 3d). Biologically, slow decay maintains patches less favorable for resident recovery, creating an opportunity for the rare invader. In contrast, if resident-conditioned soil enhances resident colonization (*σ*_*c*,22_ > 1), the invader cannot invade and slow decay of the microbial effect further prevents invasion.

### Case study 3: Parasite-triggered stage structure

Our final case study considers population structure based on individual status arising from species interactions. For example, parasite infection can divide a pest population into healthy and infected states, analogous to compartmental models in epidemiology. Here, we ask how parasite infection creates a structured population that shapes invasion between two competing pest species (Price *et al*., 1986). We considered a tri-trophic food web in which plants (*S*) are consumed by two pest species and parasites (*P*) infect these pests, converting them from a healthy state 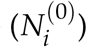 to an infected state (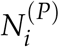 blue shade in Fig. 3e). As the infected pest lives for a period of time before eventually being killed by parasites, the two pest statuses differ in their mortality rates and conversion efficiencies. Interestingly, in this system the parasite can also be viewed as a stage-structured population, where the parasite is present either in the pest body or as free-living propagules (dotted box in Fig. 3e; Appendix S6). We studied how the parasite infection rate on pest species 1 (*β*_1_) and its plant consumption rate (*α*_1_) affect invasion of the two stage-structured populations described above. Together, these parameters allow us to examine how top-down control and resource competition jointly determine invasion in a system with interaction-generated population structure.

We examined the following two invasion scenarios: (i) the invasion criteria for the parasite invading a parasite-free equilibrium with just plants and one pest species, and (ii) the invasion criteria for pest species 1 (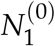 and 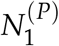) invading a plant–pest–parasite system of resident pest species 2. For both scenarios, we derived the analytical invasion condition using the methods described above (except for Method 2.1; Appendix S6). To distinguish between the two pest species, we assumed that pest species 1 has a lower consumption rate (*α*_1_ < *α*_2_) and a higher mortality rate (*m*_1_ > *m*_2_) compared to pest species 2. In compensation, it experiences a lower parasite infection rate (*β*_1_ < *β*_2_), i.e., a virulence–susceptibility tradeoff. Moreover, we assumed that parasite infection reduces the conversion efficiency of pests (*e*_*i*/*P*_ *≤ e*_*i*_). We present the invasion analysis for the parasite in Appendix S6 and focus on the invasion of pest species 1 into a resident equilibrium of species 2 below.

For the invasion of pest species 1, invasion growth rate decreases as the parasite infection rate *β*_1_ increases, reflecting stronger top-down control on the rare invader (Fig. 3f). In contrast, a higher plant consumption rate (larger *α*_1_) increases invasion growth rate by allowing pest species 1 to acquire more plant resources while rare. Consequently, increasing *α*_1_ shifts the invasion boundary toward higher values of *β*_1_; that is, pests with greater resource consumption can tolerate stronger parasite infection and still invade. Consistent with this interpretation, the analytical threshold for *β*_1_ increases with plant abundance *S*\* but decreases with parasite abundance *P*\* of the resident equilibrium (Appendix S6). Thus, biologically, invasion success reflects a balance between the invader’s advantage in resource acquisition and its susceptibility to top-down control.

## 6 Method 3: Simulation-based invasion analysis in complicated systems

So far, we have shown how to analytically solve invasion growth in systems with a stable resident equilibrium. However, theoretical (Chesson & Huntly, 1997) and empirical work (Letten *et al*., 2018) have demonstrated that fluctuations in the conditions experienced by invading species can affect coexistence. In this section, we extend our framework to fluctuating resident dynamics, where the invaders’ population transitions fluctuate over time and invasion growth rates are usually determined numerically. We first show how the concept of timescale separation (previously discussed in section 3) can guide numerical recipes for determining the invasion growth rate even when the resident population is not at equilibrium (Method 3.1). We then show how a standard mathematical technique known as Floquet theory rigorously extends the matrix population approach (introduced previously in section 4) to systems with cyclic resident dynamics (Method 3.2).

### Model extension with cyclic dynamics

As an example of non-equilibrium resident dynamics, we modified equation 1c so that species 2 also has two stages, juveniles 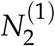 and adults 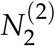 :

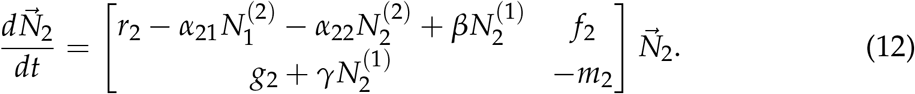

As highlighted in the model diagram (Fig. 4a, bolded terms), juveniles facilitate the juvenile-to-juvenile rate represented by *β*, and the maturation into adults represented by *γ*. When species 2 grows alone, these facilitative interactions can generate a stable limit cycle (Fig. 4b; see Appendix S7). Biologically, facilitation first allows juveniles to build up in abundance (dotted line), but population regulation is delayed because juveniles must mature into adults (dashed line) before exerting a negative effect on juveniles, which eventually causes both stages to decline. For our parameter values, the period of these oscillations is *T* = 0.778 (used in Fig. 4 as the unit of the *x*-axis).

**Figure 4:**
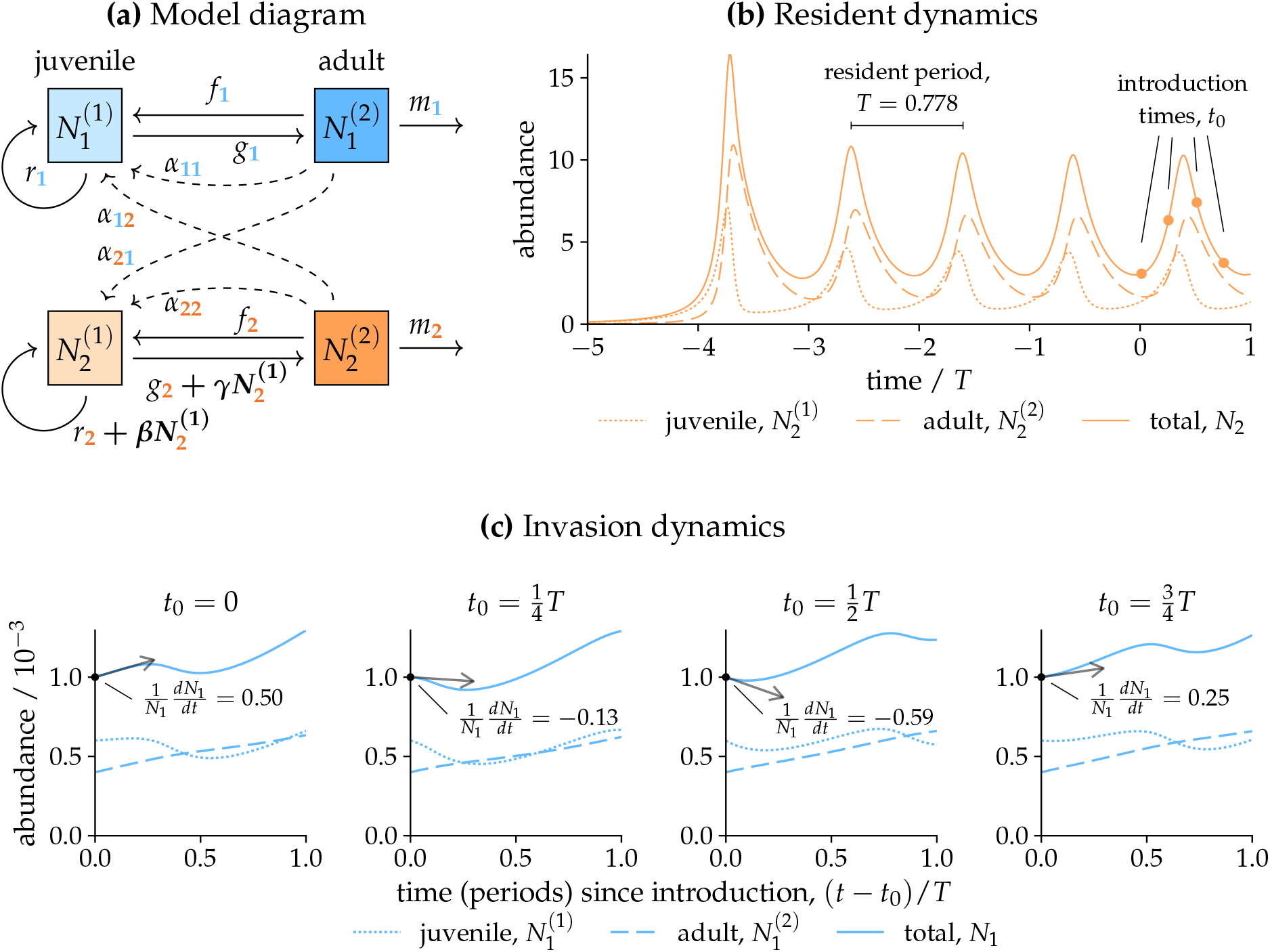
Model extension with cyclic resident dynamics. **(a)** Model diagram of the extended toy model in which both species 1 (blue) and 2 (orange) exhibit stage structure: juveniles 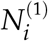 and adults 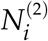 undergo maturation *g*_*i*_, juvenile replenishment *r*_*i*_, reproduction *f*_*i*_, and mortality *m*_*i*_ (solid arrows). Each species’ juvenile-to-juvenile transition is negatively affected by competition (*α*_*ij*_) from adults of both species (dashed arrows). Additionally, juveniles of species 2 facilitate the rates of their juvenile-to-juvenile transitions and maturation according to coefficients *β* and *γ*, respectively. **(b)** Time series of species 2 when growing alone as the resident, exhibiting a stable limit cycle with period *T* = 0.778. Time (*x*-axis) is scaled by *T*, such that one unit represents one complete resident cycle, and zero indicates the beginning of invasion trials (line segments indicate the four introduction times used in panel (c)). **(c)** Initial dynamics of species 1’s abundance (*y*-axis) when introduced at different points in the resident’s cycle: *t*_0_ = 0, *T*/4, *T*/2, or 3*T*/4 (from left to right; gray tangent arrows indicate the initial rate of change of the total population). Here, the *x*-axis shows time since introduction, (*t − t*_0_)/*T*, scaled by the resident period. In all time series, line types represent different population stages: juveniles 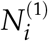 (dotted), adults 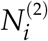 (dashed), and total *N*_*i*_ (solid). The initial population is as in Figure 1, 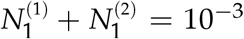, and all simulations start with 40% adults. Parameter values: *f*_2_ = 3, *g*_2_ = 1, *m*_2_ = 5, *α*_12_ = 0.5, *α*_21_ = 1, *α*_22_ = 5, *β* = 5, *γ* = 3; all other parameters are as in Figure 1.

The non-equilibrium dynamics of the resident species result in a further complication to invasion analysis. In addition to depending on initial population structure (Fig. 1b), species 1’s instantaneous growth rate depends on when in the resident’s population cycle it is introduced (shown for four different points in Fig. 4c). Because species 2’s adult abundance 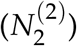 is responsible for regulating species 1’s growth rate, species 1 grows rapidly if introduced when species 2’s adults are at their lowest abundance (*t* = 0) but may initially decline if introduced at points in the cycle where adults are highly abundant (*t* = *T*/4 and *T*/2).

### Method 3.1: Simulating convergent invasion dynamics

Just as in the simple model (Fig. 1b), our simulations reveal that transient differences in the invader’s population structure rapidly disappear during invasion (Fig. 5a). Here, however, trajectories do not approach a fixed population structure, but rather converge to the same fluctuating cycle as long as the invader remains rare (circa 2 *≤ t ≤* 10). As a result, the invader’s percapita growth rate also converges to a stable cycle (Fig. 5b), and averaging this growth rate after the initial transient has decayed gives the long-term invasion growth rate that determines persistence. We can perform this calculation by identifying the interval during which species 1’s stage structure has already converged but its population remains sufficiently rare (Appendix S3, Algorithm S3.1); for our example, we find that IGR_1_ = 0.456 (Fig. 5b; dashed line shows the moving average).

**Figure 5:**
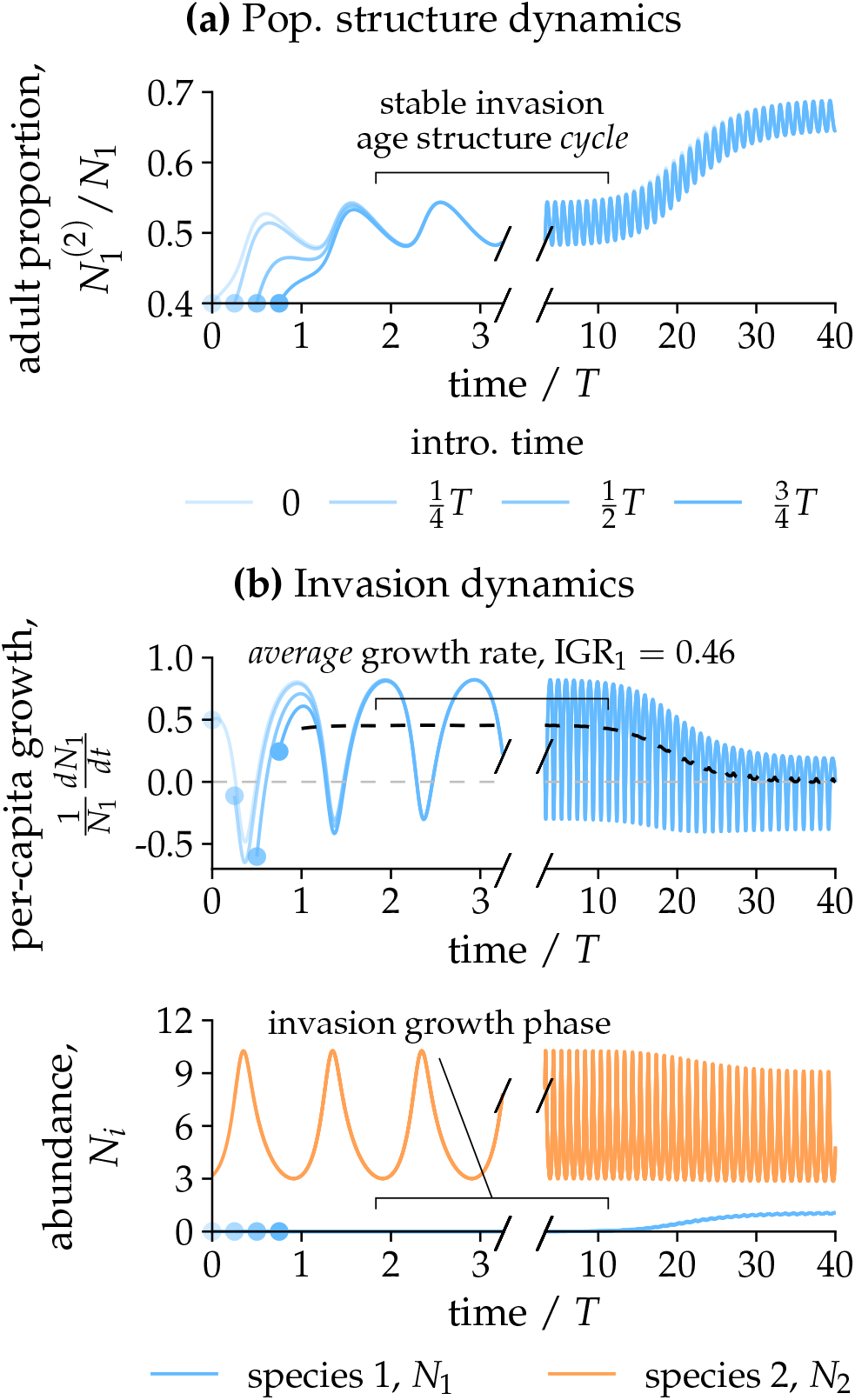
Fast and slow dynamics in the cyclic model. As in Figure 2, we plot the temporal trajectory of (a) population structure (the proportion of adults within the entire population) and (b) per-capita growth rate for the invader population (top) and the total species abundance (bottom). Parameters and invasion settings are as in Figure 4; blue colors indicate different invasion timing (light to dark blue lines: early to late invasion timing). Note that the resolution differs before (i.e., during the initial invasion) and after (i.e., subsequently approaching long-term cyclic dynamics) the break in the axis (after *t*/*T* = 3). The period when the invader reaches the same cyclic dynamics while remaining sufficiently rare gives the relevant invasion growth rate (IGR = 0.46; dashed line in (b) indicates the moving-window average of the per-capita growth rate).

As an alternative, we can estimate the same quantity using a version of the standard algorithm for calculating the *dominant Lyapunov exponent* (Benettin *et al*., 1980), of which the invasion growth rate is a special case. Briefly, we intermittently rescale the invader population size to a small value while allowing its population structure to evolve freely (see Appendix S3, Algorithm S3.2 and illustration in Appendix S8, Fig. S8.5). This numerical procedure has been used in structured population models (e.g., Werner *et al*., 2025), but the biological reason why it recovers the appropriate invasion growth rate is often left implicit. Our framework clarifies the underlying logic: rescaling provides a practical way to allow transient effects of the initial population structure to decay while maintaining the invasion scenario, such that the invader’s dynamics remain simple in the environment created by the resident. This numerical recipe is not restricted to cyclic resident dynamics. As we demonstrate in Appendix S8 for a chaotic Lotka–Volterra system (Sprott, 2010), the same approach can estimate the exponential growth rate of a rare species invading a resident community with chaotic dynamics, even when the resulting full community subsequently remains chaotic.

#### Rigorous theory for periodic systems: Floquet theory

Just as in equilibrium systems, methods based on matrix population models (Method 2) provide an efficient and mathematically rigorous way to quantify invasion growth rates in systems with cyclic resident dynamics. In this case, transitions between the invader’s stages now vary periodically as competitive pressure from the resident fluctuates over time. Fortunately, a mathematical technique known as Floquet theory shows that these periodic dynamics are closely related to the constant linear dynamics introduced in Box 2, providing a powerful technique for analyzing models of diverse natural phenomena (e.g., Barone *et al*., 1977; Klausmeier, 2008). In Box 3 and Appendix S9, we introduce the key results of this theory and show how to apply them to calculate invasion growth rates. Crucially, Floquet theory shows that the long-term dynamics of an invader (Fig. 6a, left panel) are dominated by an exponential growth term (middle panel), modulated by periodic changes in population structure around this long-term growth (right panel). Thus, Floquet theory provides the periodic analog of the matrix-population result (Box 2) by showing why population structure converges to a stable limit cycle during invasion. Moreover, it reduces the complicated dynamics of the full system to two matrices: the monodromy matrix **M** (Fig. 6b, left panel), which describes the population’s growth over discrete periods, and the Floquet exponent matrix **B** (right panel), which gives the corresponding continuous-time population growth. As we show in a worked numerical example below, calculating the leading eigenvalues of these matrices is a mathematically exact (and often much faster) method to find the invasion growth rates which we obtained by simulation in Method 3.1.

**Figure 6:**
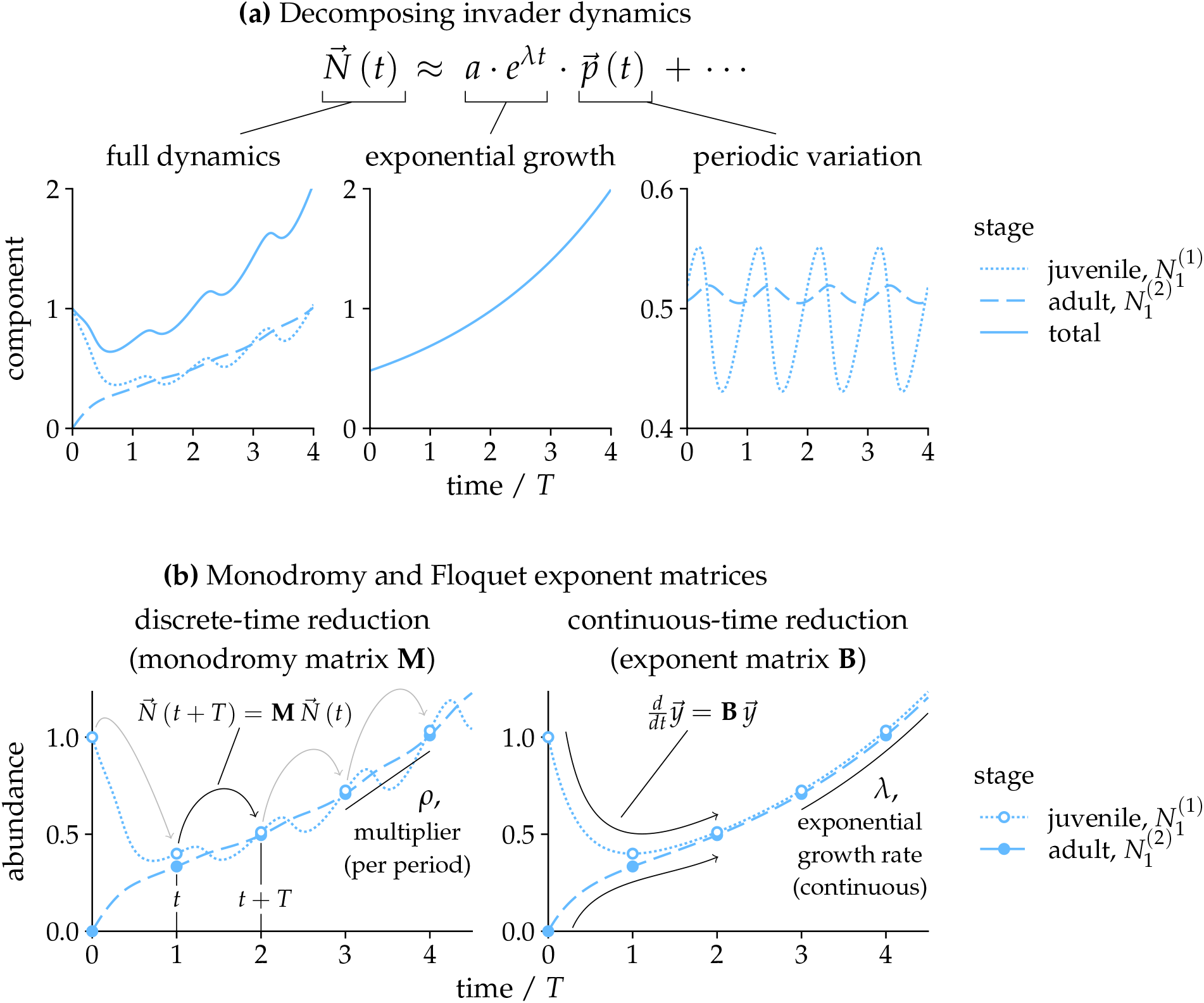
Applying Floquet theory to the cyclic model. Model and parameters as in Figure 5, except that the initial invader population (*t*_0_ = 0) consists of juveniles only. **(a)** Using the procedure in Box 3 (eq. B3.3), we can decompose invasion dynamics (left panel) into exponential growth at a rate *λ* (middle panel) and a periodic component 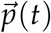 (right panel), which captures periodic population fluctuations around the overall exponential trend. **(b)** Visualization of the key mathematical objects in Floquet theory. The left plot shows how the full dynamics (lines; same as those shown in **(a)**) reduce to a simple recurrence relation (Box 3, eq. B3.2) when only considering abundances after a whole number of periods (points). Multiplication by the monodromy matrix **M** moves the system forward (arrows) by one period *T*. In the long term, each step multiplies abundances by *ρ*, the dominant Floquet multiplier (leading eigenvalue of **M**). The right plot shows the linear system (lines) associated with the matrix **B**. The system passes through the same points in the previous plot, but now advances in continuous time (arrows). In the long term, this linear system (and thus the original system as well) grows at a continuous exponential growth rate *λ*, the dominant Floquet exponent (leading eigenvalue of **B**). Components of the Floquet analysis are calculated according to Appendix S3, Algorithms S3.5–S3.8.

##### Box 3. Review: Floquet theory for growth in periodic environments

A well-established body of theory based on the work of Floquet (1883) allows us to extend the results of Box 2 to systems where transitions between population stages vary cyclically over time. Here, we summarize the application of *Floquet theory* to invasion analysis; see also Appendix S9 and Klausmeier (2008) for other ecological applications of the theory.

**Periodic linear differential equation system** Rather than a linear dynamical system with a constant transition matrix, we allow the transition matrix **C**(*t*) to vary cyclically with time such that dynamics obey a *periodic linear system*:

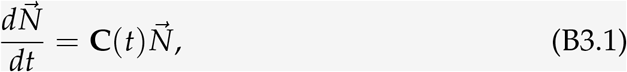

for some period *T* such that **C**(*t*) = **C**(*t* + *T*).

**Discrete-time reduction: the monodromy matrix and Floquet multipliers** Floquet (1883) showed that the complicated dynamics of the time-dependent system (eq. B3.1) are closely related to the solutions of the constant linear systems we already know how to solve (Box 2). In particular, population change over entire period lengths (e.g., *t* = 0, *T*, 2*T*, …) can be represented by a constant matrix **M** (Fig. 6b, left plot),

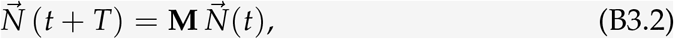

where *t* = *kT* for some whole number of periods *k*. Because the matrix **M** represents the combined effect of population transitions after the system has “run around” one full period, it is known as the *monodromy matrix* (from Greek roots meaning “single” and “run”). Effectively, eq. B3.2 represents dynamics in discrete time—much like classic discrete-time matrix population models (Lefkovitch, 1965; Leslie, 1945). Thus, the long-term population dynamics are determined by the leading eigenvalue *ρ* of **M** (known as the dominant *Floquet multiplier*), giving the factor by which the population grows over one period.

**Continuous-time decomposition and the Floquet exponent** Because eq. B3.2 dictates that solutions grow exponentially over entire periods, the continuous-time dynamics of eq. B3.1 also grow exponentially in the long run, but fluctuate within each period. The Floquet–Lyapunov theorem (Floquet, 1883; Lyapunov, 1907) shows that dynamics can be reduced to a continuous-time linear system ^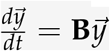^ (Fig. 6b, right plot), with transition matrix **B**, modified by some cyclically varying transformation with period *T*. Accordingly, after transient dynamics decay, the long-term dynamics can be decomposed into two components (Fig. 6a): underlying exponential growth or decline at some long-term growth rate *λ* (analogous to eq. B2.2 in Box 2) times fluctuation in abundance captured by a periodically varying vector 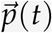:

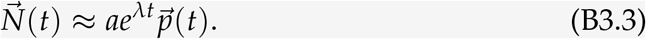

Here, *λ* is the leading eigenvalue of **B**, known as the *Floquet exponent* of the system.

**Relating the Floquet exponent to the Floquet multiplier** For invasion analysis, we are primarily interested in the Floquet exponent *λ*, which gives the invader’s long-term growth rate. Though it is possible to reconstruct the matrix **B** from the monodromy matrix **M** (Appendix S3, Algorithm S3.7), we can avoid this extra step if we are only interested in its eigenvalue *λ*. Since discrete-time growth with factor *ρ* per period describes the same system as continuous-time growth with rate *λ* (i.e., *ρ*^*k*^ = *e*^*λkT*^), we can directly relate the Floquet exponent *λ* to the Floquet multiplier *ρ*:

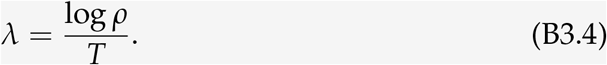

**Numerically solving for invasion growth rates** Floquet theory provides a simple and efficient method to numerically compute the monodromy matrix **M** (Appendix S3, Algorithm S3.5). For a system with *n* stages, we need information from at least *n* initial conditions to fully reconstruct **M**. The most convenient choice is to use a series of unit vectors 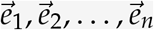 (the standard basis vectors), where a single stage has population one, and the remaining stages are absent. We obtain the monodromy matrix by integrating each of these *n* starting conditions from time *t* = 0 to *T* and pasting together the resulting vectors:

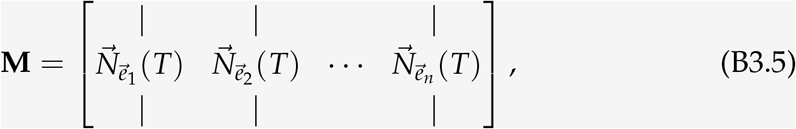

where 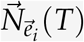 denotes the outcome when starting with only stage *i*. The invasion growth rate is then obtained by calculating the leading eigenvalue of **M** (the dominant Floquet multiplier *ρ*) and converting it to the Floquet exponent *λ* (eq. B3.4; see also Appendix S3, Algorithm S3.6).

### Method 3.2: Calculating invasion growth rates using Floquet theory

For invasion analysis, we calculate the Floquet exponent *λ* in Box 3. Applying the recipe in Box 3 for the specific model and parameters in Fig. 6, we construct the monodromy matrix **M** from numerical simulations of the invasion system over one period (from *t* = 0 to *T*). Specifically, we define a new system where the invader does not affect the resident’s dynamics (see Appendix S3, Algorithms S3.3–S3.4), and simulate this system for one period starting from two different initial conditions. With an initial population of only juveniles 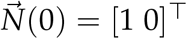, we obtain the first column: 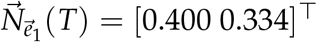. Starting from only adults 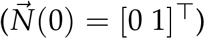, we obtain the second column 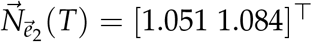. Combining these columns (eq. B3.5), we find that the monodromy matrix for time *t* = 0 is:

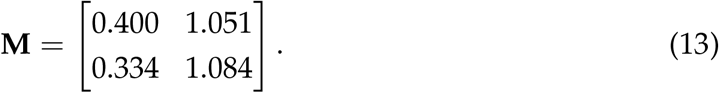

The leading eigenvalue of **M** gives the dominant Floquet multiplier *ρ* = 1.426, which describes the population growth over one period. Finally, applying eq. B3.4, the continuous-time invasion growth rate is given by the associated Floquet exponent, *λ* = log *ρ*/*T* = 0.456, which indeed matches the invasion growth rate estimated from direct simulation (Method 3.1; Fig. 5b). As calculating **M** only requires integration over a single period, this approach is faster and more accurate than simulating many periods until convergence (approximately 40*×* faster in our code).

## 7 Discussion

Invasion analysis has been a cornerstone for studying species coexistence as it provides a direct criterion for species persistence: a species with a positive invasion growth rate can recover from low abundance in the environment created by its competitors. Here, we developed a general framework for applying invasion analysis when the invading species is structured into multiple states. The organizing principle is that invasion growth in structured populations should be treated as a long-term property of the rare structured population, rather than as the instantaneous growth of any particular initial state distribution. Building on this insight, we presented two complementary analytical approaches: (1) timescale separation and variable transformations, which provide a biologically intuitive route to invasion growth rates, and (2) matrix population theory, which provides an eigenvalue-based mathematical description of the same convergence process. We also showed that the same logic can guide numerical estimation of invasion growth rates under complex dynamics where analytical expressions are unavailable. Together, these approaches provide an integrated toolkit linking insights from matrix-population theory to community-level ecological questions about species coexistence.

The power of invasion analysis comes from simplifying the full community dynamics: otherwise complicated models are reduced to a smaller and much more tractable subsystem (i.e., the invasion scenario). The pieces of our analytical toolkit give mathematically equivalent answers, but researchers can choose an appropriate method based on familiarity, arithmetic convenience, and the particular information required to answer their questions. In many studies, the key information is not the exact magnitude of the invasion growth rate, but rather its sign (Ellner *et al*., 2020; Pande *et al*., 2020). In this spirit, both analytical approaches presented here include shortcuts for deriving invasion criteria directly, without solving the full expression of the invasion growth rate (see also Appendix S2, subsection S2.1.2 for another mathematical shortcut). Nonetheless, if the focal population has relatively few stages, and a researcher is specifically interested in the exact population structure realized during invasion, direct analytical timescale separation (Method 1.1) may be useful for its conceptual simplicity. Meanwhile, a researcher working on a population with many stages must focus on deriving a simple invasion condition, since the exact expression for invasion growth rate may not be analytically solvable. Here, a reasonable approach would be to search first for a simple strategic linear combination (Method 1.2) and, if this fails, to fall back on the full stability conditions (Method 2.2). A well-chosen analytical approach pays off: our case studies highlight that even in complicated models, it is often possible to identify surprisingly simple rules for population persistence. For example, in the cannibalistic herbivores model, while the level of aggression matters for the magnitude of the invasion growth rate, invasion success itself is determined solely by the demographic effect of consuming intraguild prey (*b*_21_). Thus, the power of analytical approaches lies in their ability to translate ecological complexity into interpretable invasion criteria.

By rigorously extending invasion analysis to structured populations, our framework reinforces the role of invasion growth rates as a common currency for ecological success and persistence (Grainger *et al*., 2019). Indeed, the same approaches apply to systems beyond those in the main text, including chaotic dynamics (Appendix S8) and models in discrete time (Appendix S1). Appendix S10 illustrates the latter by deriving simple invasion conditions for plants experiencing discrete growth seasons, a workhorse system for studies on coexistence theory (e.g., Song *et al*., 2021; Werner *et al*., 2025). This opens the door to a variety of analyses. As a starting point, invasion analysis enables researchers to apply the well-known mutual invasibility criteria for coexistence (Box 1). Further decomposed, invasion growth rates can be used to partition coexistence outcomes into contributions from different ecological mechanisms (Barabás *et al*., 2018; Ellner *et al*., 2019; Letten *et al*., 2018) or summarize them into niche and fitness differences (Chesson, 2000; Spaak & De Laender, 2020). As current work extends and compares alternative partitioning approaches across increasingly complex communities (Song & Spaak, 2024; Spaak *et al*., 2023; Wan *et al*., 2025), starting from simple, interpretable expressions for invasion growth rates will be particularly important. Analytical solutions for the invasion of structured populations may also help resolve long-standing questions in coexistence theory. For instance, while many well-known mechanisms contributing to coexistence involve population structure, e.g., annual–perennial life-history differences (Werner *et al*., 2025), previous analytical treatments have relied on strong simplifying assumptions (DeMalach *et al*., 2021; Song *et al*., 2021). Finally, we caution that there are exceptional cases where the mutual invasibility criterion is not directly applicable. Despite this limitation, invasion analysis still forms the basis for newer approaches addressing these shortcomings, including in communities with positive density dependence (i.e., strong Allee effects: Schreiber *et al*., 2019; Walker & Gilbert, 2025) or complicated assembly rules (i.e., the invasion graph approach: Brennan & Schreiber, 2026; Hofbauer & Schreiber, 2010; Spaak & Schreiber, 2023).

Our framework is closely related to established mathematical procedures in diverse fields, including complex systems, epidemiology, and evolutionary biology. Our numerical procedure for estimating invasion growth rate in cyclic or chaotic resident dynamics is closely related to standard algorithms for estimating dominant Lyapunov exponents (Benettin *et al*., 1980), reflecting the fact that the invasion growth rate is a special case of this more general metric. In permanence theory, these exponents characterize the long-term growth rate of a rare structured population in a fluctuating environment and therefore provide a natural persistence criterion (Hofbauer & Schreiber, 2010; Roth & Schreiber, 2014). Meanwhile, our linear algebra-based approach mirrors a widely used approach from disease ecology and epidemiology, the next-generation matrix (Diekmann *et al*., 1990, 2009). Developed to derive the basic reproduction number in compartmental epidemic models, this technique evaluates the matrix of transition rates between infected states at the disease-free equilibrium. Finally, when population composition closely tracks environmental change, timescale separation techniques have also been used to derive effective population-level responses of structured species to changing environments (Kussell & Leibler, 2005; Letten *et al*., 2024).

By placing our work within a broader family of approaches that reduce complex state-dependent dynamics to interpretable criteria for long-term population growth, these connections point towards new syntheses between community ecology and other subdisciplines. For instance, because invasion growth rate is also the fundamental measure of fitness in evolutionary biology (Yamamichi *et al*., 2022), invasion analysis of stage-structured mutants provides important general insight into eco-evolutionary feedbacks (Bukkuri & Brown, 2023; Iritani & Day, 2026). Beyond the core of population biology, invasion analysis is being applied to an ever-expanding set of questions, from biological invasions (Iles *et al*., 2016; MacDougall *et al*., 2009) to biogeochemical cycling (Smith & Wan, 2019; Xu *et al*., 2026) and responses to climate change (Usinowicz & Levine, 2018). As such, we anticipate a growing number of research programs will incorporate population-level biological detail into models of species interactions. Amid this increasing complexity, ecologists must retain the interpretability that has made simple community models so powerful. We hope that general theory from population ecology, as synthesized here, will continue to provide ecologists with a guiding thread towards general understanding.

## Acknowledgments

We thank Adam Clark, Takeshi Miki, and members of the Ke lab for discussions.

## Statements and Declarations

### Author Contributions

JW and PJK conceived the study. JW led the analysis, with contributions from HCL, GYC, SCH, JZL, CTT, and PJK. JW and PJK wrote the manuscript with input from all authors.

### Data Availability

No original data were presented in this manuscript. All code and inputs required for the simulations and re-analyses in this manuscript are publicly available at https://github.com/joe-wan/structured-invasion-code/.

## Supporting Information for

### S1 General framework: matrix population models

When individuals of a species differ from each other in some way, we must model the population in more detail than allowed by a single state variable (main text, Box 2). One way to do so is to consider transitions of species *i* between *m*_*i*_ different discrete states, representing age, size, habitat, disease status, or some other condition(s) of interest. In this Appendix, we describe the general application of matrix population models to these systems.

#### S1.1 Continuous-time dynamics

Representing the abundance of species *i* in state *k* as 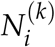, we can write a continuous-time system as

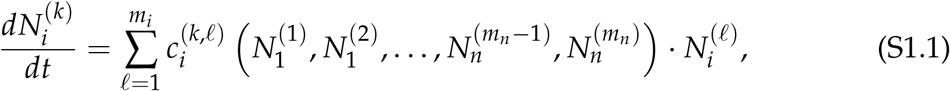

where 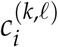 is the transition rate from state *l* to state *k* of species *i*, which may depend on the abundance of all states of all species in the system. We can also write the state distribution of species *i* as an *m*_*i*_-dimensional vector

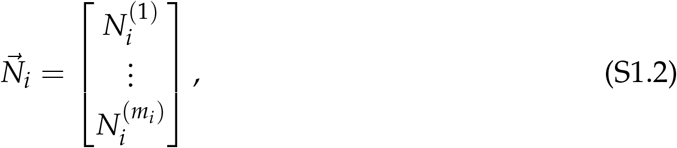

such that the population dynamics of species *i* can be represented by the vector differential equation

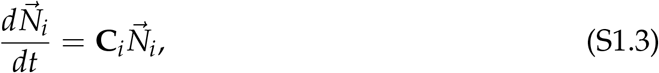

where **C**_*i*_ is the *m*_*i*_ *× m*_*i*_ transition matrix between states of species *i*, dependent on the state distributions of all species in the system:

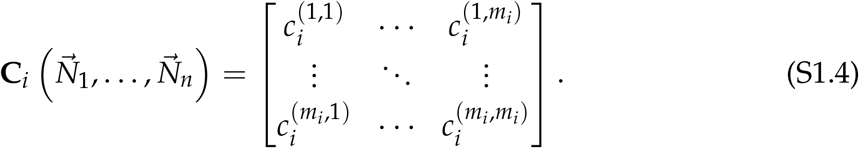

When conducting invasion analysis, we consider some resident state where 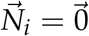 and the other species are at an equilibrium 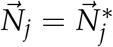 for all *j≠ i*. Evaluating the transition matrix at this resident equilibrium gives

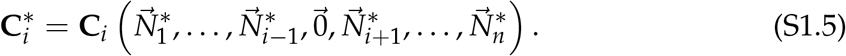

When we are near this invasion state, **C**_*i*_ is near some value which does not depend on the invader abundance, which we will denote 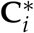. In other words, the transition matrix is independent of the invader’s dynamics because it is too rare to exert feedbacks on its own population dynamics. Equivalently, 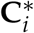 can be viewed as the Jacobian matrix for the invading species, evaluated at the resident equilibrium. In the following text, we will focus on a particular species *i* in the invasion condition, writing 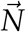 for 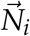 and **C** for 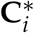 in the interest of brevity.

#### S1.2 Discrete-time dynamics

Though the main text focuses on continuous-time systems, it is worth also considering the discrete-time analog of this problem. Writing 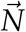 for 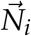 as above, we can write the abundances of a species’ stages at the next time step *t* + 1 in terms of its abundances at the current time *t*:

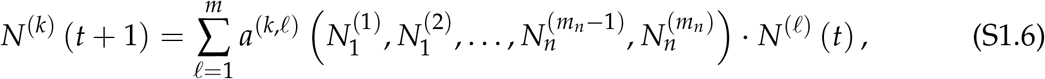

or in matrix form,

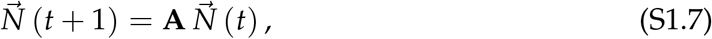

where **A** follows the notation used for this class of models in Caswell (2001). Here, considering the invasion scenario as in equation S1.5, *m* = *m*_*i*_ is the number of stages of the invader, and *a*^(*k,l*)^ is the expected contribution to stage *k* at the next time step per individual in stage *l* at the current time step, evaluated at the resident equilibrium. This class of discrete-time models is well-studied in population ecology. In particular, if the classes *k* = 1, 2, …, *m*_*i*_ represent age groups, **A** is known as the Leslie matrix (after Leslie (1945)); more broadly, if *k* instead represents developmental, size, or life-history stages, **A** may be termed a Lefkovitch matrix (after Lefkovitch, 1965). Standard textbooks (e.g., Caswell, 2001) treat the analyses of these models in terms of **A** and its eigenvalues in depth.

Note that this **A** has a different interpretation from our continuous-time **C**: elements on the diagonal represent the members of a particular class which *stay* in that class between times *t* and *t* + 1, as opposed to in the continuous-time formulation above, where diagonal elements reflect change in a particular class. As a formulation more compatible with the continuous-time equation above, we can rewrite the system as a difference equation,

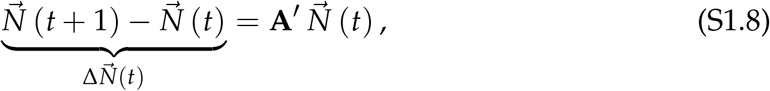

where **A**′ = **A** *−* **I** is obtained by subtracting the identity matrix (i.e., subtracting one from each diagonal entry) to account for the members remaining in this class. Indeed, taking the limit of this difference equation model as the time step becomes small recovers the ordinary differential equation formulation above. To focus on connections between the discrete- and continuous-time formulations, it will be convenient to use discrete-time **A**′ as an analogue of **C** to highlight similarities between analyses of the two types of models.

#### S1.3 Eigenvalue analysis of state transition matrices

The eigenvalues of **C** are the values *λ* = *λ*_1_, *λ*_2_, … such that

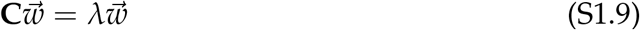

for some non-zero vector 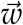. The value of 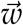 is associated with a particular eigenvalue *λ* and is known as its associated eigenvector. In general, if **C** is an *n × n* matrix, there will be up to *n* unique eigenvalues of **C**, which may be complex-valued. To solve for the eigenvalues, a convenient procedure is to solve the equation

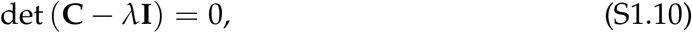

which gives a polynomial equation of degree *n* in *λ*, known as the *characteristic polynomial* of **C**. Biologically, an eigenvalue/eigenvector pair *λ*_*i*_, 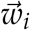 represents a solution to equation S1.3 where the elements of the population vector 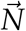 remain in a fixed proportion determined by 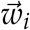, but its magnitude grows or shrinks at an exponential rate *λ*_*i*_;

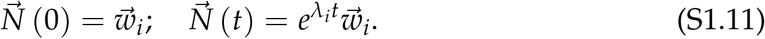

Even if our starting population state 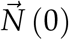 is not an eigenvector of **C**, we will usually be able to represent it as some combination of eigenvectors. The fact that equation S1.3 is a linear system under the invasion scenario in equation S1.5 means that the time-dependent population trajectory is also a combination of the trajectories determined by individual eigenvalues/eigenvectors,

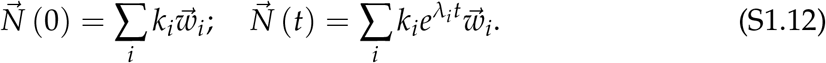

In fact, if we assume the eigenvalues are ordered by decreasing real parts, with *λ*_1_ being the eigenvalue with the greatest real part (known as the leading eigenvalue), we see that

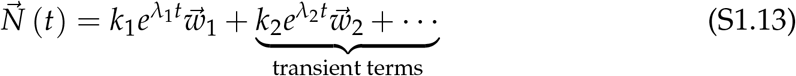

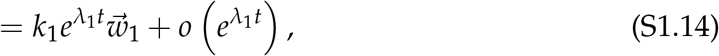

where the terms associated with the non-leading eigenvalues are transient, because their influence becomes small relative to the first term as time *t* becomes large. Here, the little-o notation 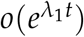 indicates that the transient terms grow more slowly than 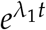. Accordingly, the long-term growth rate is determined by the leading eigenvalue *λ*_1_ (for brevity, we denote it as *λ* when the context makes it clear, as in the main text) and the long-term population structure approaches 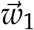 (similarly abbreviated 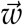 in the main text), giving the result in eq. B2.2 of the main text. If *λ*_1_ is positive, the population maintains positive growth.

There is an important mathematical complication: generally, some eigenvectors may have components with negative signs. Does this imply the long-term stage structure is not biologically meaningful because it contains negative abundances? Fortunately, it turns out that the special properties of **C** guarantee that this problem cannot arise: because the off-diagonal elements of **C** are nonnegative, solutions which start positive will remain positive (Box S1.B1). Furthermore, due to the Perron–Frobenius theorem (see Box S1.B2), the leading eigenvalue *λ*_1_ is real and its eigenvector 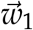 has nonnegative components. For sensible population models, **C** is irreducible (Box S1.B3) and we furthermore have that 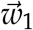 is uniquely defined up to a constant; i.e., the population’s stages approach a single well-defined set of proportions.

In discrete time, we can write

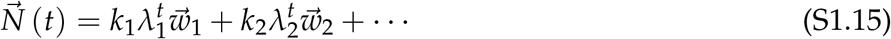

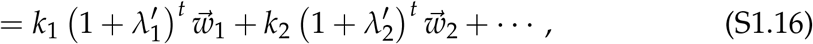

where *λ*_1_, *λ*_2_, … are the eigenvalues of **A**, 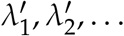 are the eigenvalues of **A**′, and 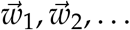 are the associated eigenvectors. In an exact analog to the continuous-time result, the population maintains positive growth if the leading eigenvalue 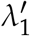 of **A**′ is positive (or equivalently, if **A**’s dominant eigenvalue *λ*_1_ is greater than one).

Box S1.B1. Special properties of C

##### Box S1.B1.

Special properties of **C**

In biologically realistic models, a particular individual cannot give rise to a negative abundance of individuals in a different stage. This means the off-diagonal elements of the matrix **C**, *c*^(*k,l*)^ for *k≠ l*, should be nonnegative. Biologically, this condition ensures that trajectories starting from nonnegative populations remain nonnegative for all stages. This follows from the fact that **C** is a member of a special class of matrices known as Metzler matrices.

**Metzler matrices** A matrix **M** where all off-diagonal elements are nonnegative (i.e., (**M**)_*ij*_ *≥* 0 for all *i≠ j*) is known as a Metzler matrix. Named after Metzler (1945), who first applied them in economics, this class of matrices naturally describes the evolution of many systems whose elements take nonnegative values. Consider the system of linear ordinary differential equations associated with a matrix **M**,

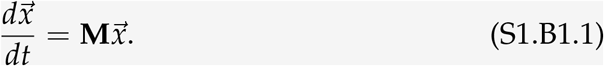

Ważewski (1950) proved that all solutions starting from nonnegative 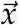 remain nonnegative if and only if **M** is a Metzler matrix. To see this, consider the following necessary and sufficient condition for the system to avoid becoming negative: when *x*_*i*_, the *i*th element of 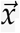, is zero, its derivative *dx*_*i*_/*dt* must be nonnegative. Writing out the element of the matrix-vector product, this condition is ∑_*j*_ (**M**)_*ij*_ *x*_*j*_ *≥* 0; furthermore, since *x*_*i*_ = 0, we can write ∑_*j≠i*_ (**M**)_*ij*_ *x*_*j*_ *≥* 0. This holds generally for all nonnegative 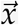 if and only if all non-diagonal entries of **M** are nonnegative; (**M**)_*ij*_ *≥* 0 for *i≠ j*.

**Discrete-time nonnegativity** In discrete time, let 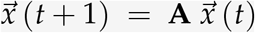. Here, 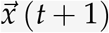 remains nonnegative for all nonnegative 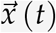 if and only if **A** is a non-negative matrix (i.e., all elements of **A** are nonnegative). Equivalently, since **A**′ = **A** *−* **I**, all non-diagonal elements of **A**′ must be nonnegative, and the diagonal elements must be at least *−*1, i.e., (**A**′)_*ii*_ *≥ −*1 for all *i*. Thus, for the system to remain nonnegative, it is a necessary (but not sufficient) condition that **A**′ is a Metzler matrix.

##### Box S1.B2. The Perron–Frobenius theorem

A foundational result in linear algebra, known as the Perron–Frobenius theorem (Frobenius, 1912; Perron, 1907), states that a matrix with nonnegative entries has a real, nonnegative eigenvalue *λ*\* (1) whose magnitude is greater than or equal to the magnitude of any other eigenvalue, and (2) whose associated eigenvector 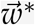 can be chosen to have all nonnegative components. Furthermore, if the matrix is irreducible (Box S1.B3), 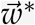 is uniquely defined up to a constant.

**Leading eigenvalues** We apply this to show our result for **C**, which can have negative elements on its diagonal. Define a new matrix **D** whose elements are all nonnegative by letting **D** = **C** + *b***I**, where *b* > 0 is a large enough positive constant. Because 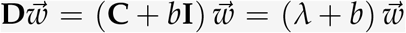, each eigenvalue *λ* of **C** corresponds to an eigenvalue *λ*′ of **D** according to *λ*′ *− b* = *λ*; moreover, if 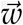 is an eigenvector of **C** associated with eigenvalue *λ* (i.e., 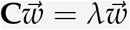), then it is also an eigenvector of **D** associated with the eigenvalue *λ*′.

Because **D** has all nonnegative elements, the Perron–Frobenius theorem guarantees there is a real nonnegative *dominant* eigenvalue *λ*^\**′*^ whose magnitude is at least as great as that of any other eigenvalue *λ*, i.e., *λ*^\**′*^ *≥* |*λ*|. Because *λ* and *λ*^\**′*^ are distinct eigenvalues, any other eigenvalue *λ* must have a strictly lesser real part than this dominant eigenvalue, Re (*λ*) < *λ*^\**′*^. Thus, *λ*^\**′*^ is also the *leading* eigenvalue of **D**. Moreover, it is guaranteed that we can choose an eigenvector 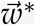 associated with *λ*^\**′*^, whose components are all nonnegative.

Since the eigenvalues of **C** have simply been shifted by *−b*, **C** is also guaranteed to have a leading eigenvalue *λ*\* = *λ*^\**′*^ *− b*. (Note that, unlike *λ*^\**′*^, this eigenvalue may be negative, and need not be the dominant eigenvalue.) Furthermore, the nonnegative eigenvector 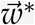 of **D** is also the eigenvector of **C** associated with the leading eigenvalue *λ*\*.

##### Box S1.B3. Irreducible matrices

An *n × n* matrix **M** is irreducible if, for any two distinct indices *i* and *j* such that 1 *≤ i, j ≤ n*, we can find a sequence of distinct indices *s*_1_, *s*_2_, …, *s*_*m*_ such that *s*_1_ = *i, s*_*m*_ = *j*, and 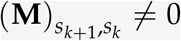 for all *k* such that 1 *≤ k ≤ m −* 1. In other words, if we view each element of the matrix 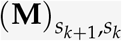 as representing the transition of individuals from stage *s*_*k*_ to stage *s*_*k*+1_, irreducibility of **M** is equivalent to saying that individuals of any stage can eventually end up in any other stage after some sequence of population transitions—a natural condition for considering all stages as part of the same population. Thus, this condition will naturally be satisfied in biologically realistic models of transitions within a single population.

**Properties of irreducible Metzler matrices** If the continuous-time population transition matrix **C** is irreducible, the shifted matrix **D** is also irreducible, since only the off-diagonal entries (which remain unchanged) matter for the definition of irreducibility above. In this case, a stronger version of the Perron–Frobenius theorem states that the leading eigenvalues of **C** and **D** are *algebraically simple*. Among other properties, this means that they are associated with a positive eigenvector which is unique up to a constant. Precisely stated, if *λ*\* is the leading eigenvalue of **C**, then there is an eigenvector 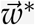 whose elements are all positive satisfying 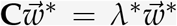 ; furthermore, the only positive vectors satisfying this condition are positive multiples of 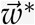. The same argument holds for discretetime **A**′. Biologically, this means that population dynamics converge to the same long-term set of non-zero state proportions, regardless of the starting conditions.

#### S1.4 Eigenvalue analysis with stability conditions

To evaluate whether continuous-time **C** or discrete-time **A**′ admits long-term population growth (i.e., persistence of the invader population), we need to know the sign of the leading eigenvalue. In systems with three or more stages, however, directly solving for this eigenvalue may often be infeasible. As an alternative, we can apply mathematical conditions from *stability analysis* which diagnose whether the matrix has a nonnegative leading eigenvalue. This is closely related to invasion analysis: a system where invasion is possible is an unstable system, because the small perturbation caused by introducing the invader grows in magnitude.

In this section, we present several recipes for stability analysis in systems of varying complexity. Box S1.B4 presents frequently used stability conditions which can directly be applied to small systems with two to five stages, while Box S1.B5 gives the general Routh–Hurwitz criteria from which these conditions are derived.

Finally, in very complex cases, it may be preferable to only focus on the region of parameter space near the invasion boundary. In this case, the invasion condition can be derived using the *invasion determinant*, summarized in Box S1.B6. This approach is naturally suited for evolutionary invasion analyses, where the system is always near the invasion boundary because the traits of the invader and resident differ only infinitesimally (Iritani & Day, 2026). More generally, however, caution is warranted because this approach may recover spurious boundaries in addition to the correct invasion condition.

##### Box S1.B4. Stability conditions for small matrices

For an *n × n* matrix **M**, find the characteristic polynomial using eq. S1.10. Then, we can write the coefficients as *a*_0_, *a*_1_, *a*_2_, …, *a*_*n*_ such that the polynomial is

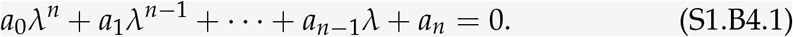

Note that some texts use the opposite convention for the subscripts of *a*, where *a*_*i*_ is the coefficient of the *λ*^*i*^ term. Assume that the leading coefficient is one, *a*_0_ = 1. We can always rewrite the polynomial in this form by dividing the polynomial through by the highest-order coefficient (alternatively, solving det (*λ***I** *−* **M**) = 0 will always give a highest-order coefficient of 1). Routh (1877) and Hurwitz (1895) provided a simple general test to determine when all eigenvalues of **M** have negative real parts, summarized in Box S1.B5. Here, we give the frequently-encountered special cases for *n* = 2 through *n* = 5.

**Two-dimensional matrices** In two-dimensional systems, the conditions for stability are particularly simple. In particular, all eigenvalues have negative real parts if and only if

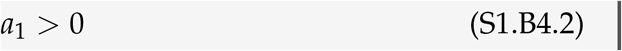

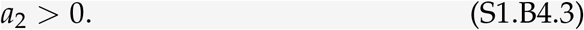

Furthermore, by expanding the determinant in det (*λ***I** *−* **M**), we see that *a*_1_ = *−* tr **M** and *a*_2_ = det **M**. Noting that the trace is the sum of the eigenvalues and that the determinant is the product, the stability conditions can also be rewritten as *λ*_1_ + *λ*_2_ = tr **M** < 0 and *λ*_1_*λ*_2_ = det **M** > 0.

**Three-dimensional matrices** In three-dimensional systems, the eigenvalues all have negative real parts if and only if the following three conditions hold:

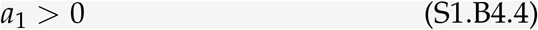

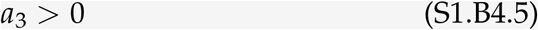

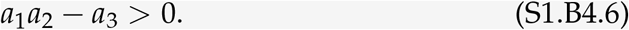

**Four-dimensional matrices** In four dimensions, the conditions become

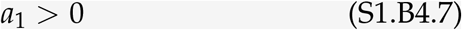

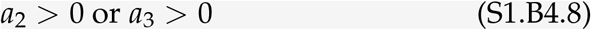

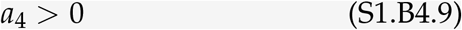

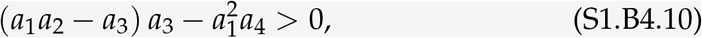

where we may choose *a*_2_ > 0 or *a*_3_ > 0 depending on which one is easier to calculate, and the last condition may also be written as 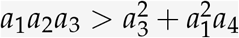 .

**Five-dimensional matrices** In five dimensions, it suffices to show that the following conditions hold

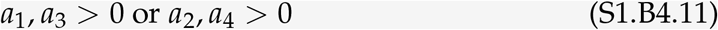

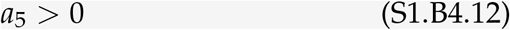

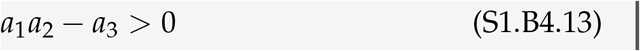

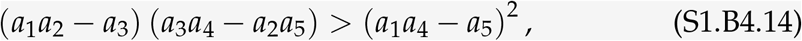

where we can choose whether to check positivity of *a*_1_, *a*_3_ or of *a*_2_, *a*_4_.

##### Box S1.B5. General Routh–Hurwitz conditions

Say we have a general characteristic polynomial

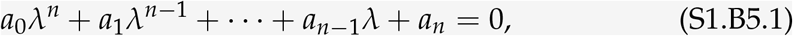

where *a*_0_ > 0. Define the Hurwitz matrix **H**, an *n × n* matrix such that

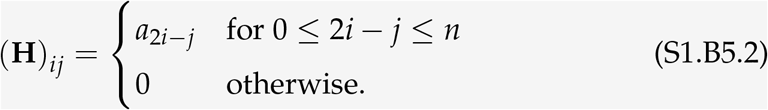

The resulting matrix has the structure

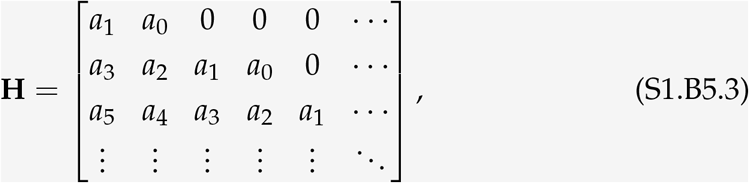

where we note that the indices in each row decrease by one, the indices in each column increase by two, and entries where the index would be outside the range from 0 to *n* are replaced with zeros. Note that some texts (e.g., the original work of Hurwitz (1895)) define **H** as the transpose of the above matrix instead, but this does not affect the stability calculation.

Now, let **H**_*k*_ denote the *k × k* square matrix obtained by only taking the first *k* rows and columns of **H** (i.e., the *k*th leading principal submatrix of **H**). Then, assuming that *a*_0_ > 0, stability occurs if and only if all of these submatrices have positive determinants, i.e.,

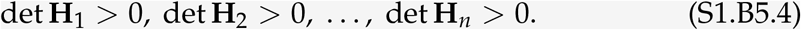

Generally, it is a necessary condition for stability that all *a*_*i*_ are positive. Indeed, Liénard & Chipart (1914) showed that the number of determinants which need to be calculated can be greatly reduced if we know the signs of some of the *a*_*i*_ coefficients. The simplified Liénard–Chipart conditions show that stability can be proven by showing that approximately half of the *a*_*i*_ coefficients and half of the determinants are positive. In particular, stability occurs if and only if two conditions hold: first, at least one of the following coefficient conditions is satisfied:

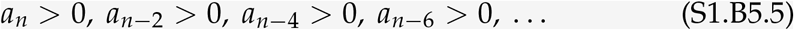

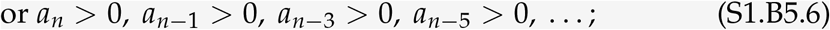

and second, at least one of the following determinant conditions is satisfied:

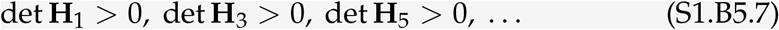

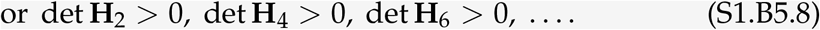

In other words, we only need to check if (1) all odd-numbered or all even-numbered coefficients, as well as *a*_*n*_, are positive, and (2) all odd-numbered or all even-numbered determinants are positive. Applying these conditions (always choosing the set of determinants with the smallest maximum order) and letting *a*_0_ = 1, we straightforwardly obtain the three-, four-, and five-dimensional criteria in Box S1.B4.

##### Box S1.B6. Iritani & Day (2026)’s invasion determinant

A necessary condition for stability is that all *a*_*i*_ coefficients are positive in the characteristic polynomial

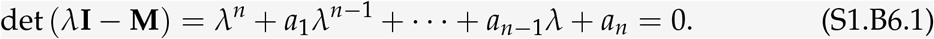

Furthermore, note that *a*_1_ = tr (*−***M**) and *a*_*n*_ = det (*−***M**) for any *n*. Thus, two necessary conditions for stability are

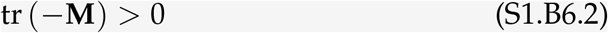

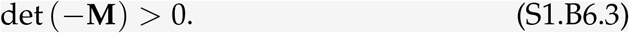

In the two-stage case above (eqs. S1.B4.2 and S1.B4.3), these conditions are also sufficient for stability. More generally, the trace and determinant conditions alone do not suffice to show stability. When conducting invasion analysis, however, there are special cases where it may suffice to only consider the determinant condition (eq. S1.B6.3) instead of fully evaluating all stability conditions (Boxes S1.B4 and S1.B5).

**Invasibility near the invasion boundary** Say we are only interested in deriving the boundary where changes in some underlying parameter(s) cause the invasion growth rate to cross continuously from negative to positive. In other words, we assume that the leading eigenvalue of the matrix **C** is near zero. Furthermore, as before, assume the special properties of **C** required for biological realism hold as this transition occurs: that it has only nonnegative off-diagonal elements (i.e., is a Metzler matrix; Box S1.B1) and that it is irreducible (Box S1.B3). Then the transition from negative to positive invasion growth (i.e., stability to instability) occurs when the leading eigenvalue *λ*\* of **C** guaranteed by the Perron–Frobenius theorem (Box S1.B2) crosses from negative to positive, with the real part of all other eigenvalues remaining negative. Since *λ*\* is an algebraically simple leading eigenvalue and the determinant can be understood as the product of all eigenvalues, this implies that det (*−***C**) changes sign from positive (a necessary condition for stability) to negative at the invasion boundary.

**The invasion determinant condition** Thus, the condition for invasibility *near the invasion boundary* under our assumptions is

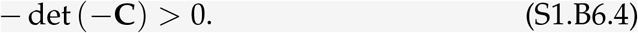

The left-hand side of this expression, termed the *invasion determinant* by Iritani & Day (2026), serves as a useful proxy when performing analytical investigations: near the invasion boundary, it shares the same sign as the true invasion growth rate. Nonetheless, some caution is warranted in general invasion analyses: while a positive value of the left-hand side always implies invasibility (i.e., the system must be unstable because det (*−***C**) = *a*_*n*_ < 0), negative values may also occur in an invasible system due to a non-leading eigenvalue crossing zero. Thus, solving the boundaries defined by inequality S1.B6.4 may recover spurious boundaries in addition to the correct invasion boundary. In these cases, comparison to numerical simulations or additional analyses may be needed to confirm the results of determinant-based calculations.

## S2 Detailed analyses for simple stage-structured model

We consider the simple model of a stage-structured invader species from the main text (eq. 1 and Fig. 1).

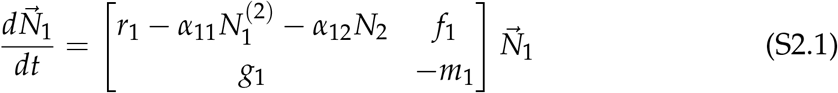

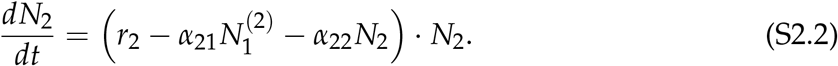

The resident equilibrium of species 2 is

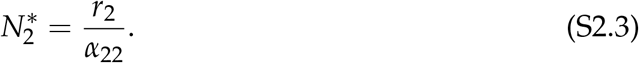

Since we are interested in species 1’s invasion condition, we set 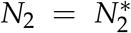 for all subsequent derivations in this Appendix.

### S2.1 Method 1.1: Direct timescale separation

#### S2.1.1 Timescale separation using transformations in the main text

We will choose a new pair of variables: *N*, representing the slow dynamics of population growth, and *Q*, representing the fast dynamics of population stage structure. Since competition only depends on adults (stage 2), we choose 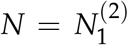 as a convenient choice for the population (slow) variable. Accordingly, a convenient choice of the stage structure (fast) variable is the ratio of juveniles (stage 1) to adults (stage 2), 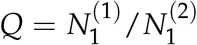. Then 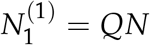 and we can rewrite the system as:

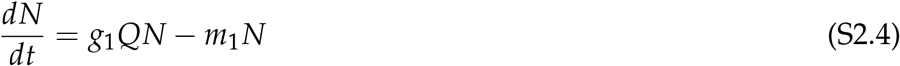

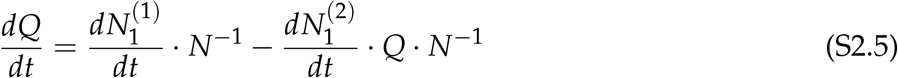

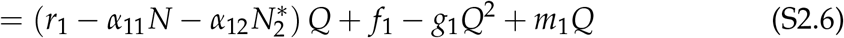

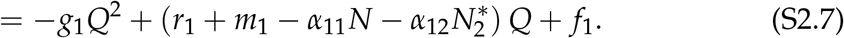

Note that many other choices of *N* and *Q* would be valid here. While the choice above simplifies the algebra considerably, we could also have selected *N* to be the total population and *Q* to be the proportion of adults within the total population (demonstrated in subsection S2.1.3).

First, we solve for the quasi-equilibrium of *Q* in the invasion condition. In this case, *N ≈* 0, allowing us to neglect the term *−α*_11_*N*. Solving using the quadratic formula gives the single biologically meaningful root,

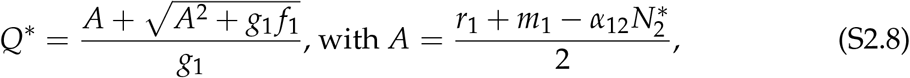

where we note that the other root is necessarily negative and therefore not biologically meaningful (since *Q* is a ratio of populations). Next, we find the exponential growth rate of *N* when *Q* = *Q*\*. We find that the growth rate of *N* is expressed as:

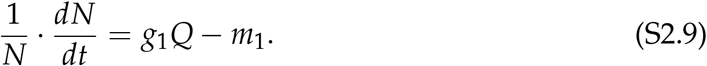

Substituting *Q* = *Q*\* into the per-capita growth rate, we find

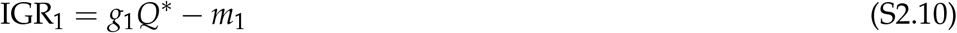

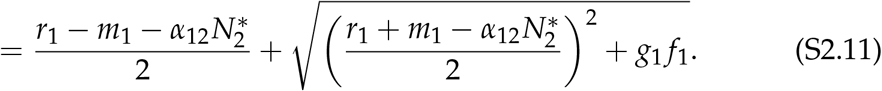

Finally, we solve for the conditions under which the invasion growth rate is positive. Noting that

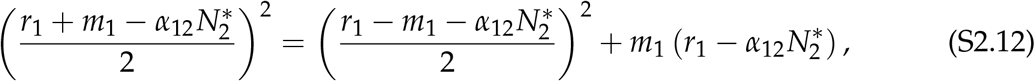

we can rewrite

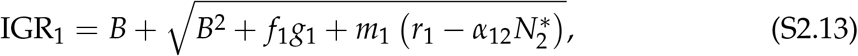

with

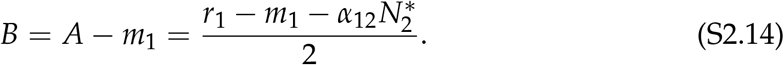

We note that the invasion growth rate is positive (1) when 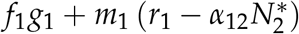 is positive, which rearranges to

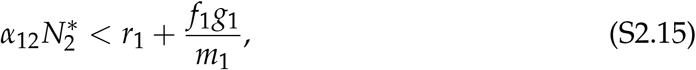

or (2) when *B* is positive, which occurs when 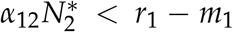. Since the second condition also implies the first, the first condition is necessary and sufficient for positive invasion growth.

#### S2.1.2 Simplified derivation: the *λ*-trick

There is also a convenient alternative to the full calculation of the quasi-equilibrium stage ratio above. Applying the chain rule to differentiate 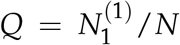 (eq. S2.5), we see that the result can be written in terms of the per-capita growth rate of *N*, i.e., (1/*N*) *· dN*/*dt*:

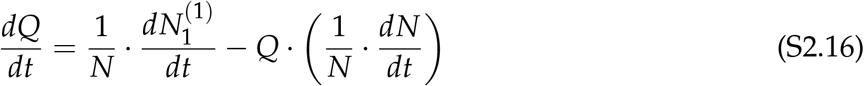

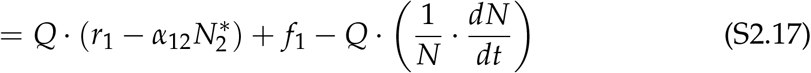

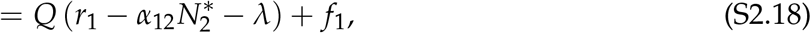

where we applied the invasion condition in the second line, and in the last line, we have denoted *λ* = (1/*N*) *· dN*/*dt* for convenience. We note that this per-capita growth *λ* is precisely the invasion growth rate in which we are interested. Accordingly, we can solve the quasi-equilibrium *Q*\* in terms of the unknown *λ*, finding that

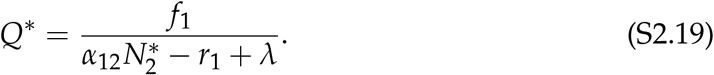

Substituting this into the expression for invasion growth rate (eq. S2.9), we find that

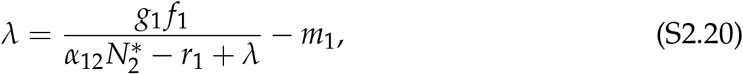

where we have also re-expressed the per-capita growth rate on the left-hand side as *λ*. Now we could rearrange this equation to a quadratic in *λ* and solve to obtain the same result as above. However, in the case where we are only interested in the boundary between successful and unsuccessful invasion, we can take the shortcut of considering when the parameters in the system and the value of 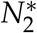 cause the system to be on this invasion boundary. To do so, we substitute *λ* = 0 into the above expression and rearrange to find that the boundary occurs when:

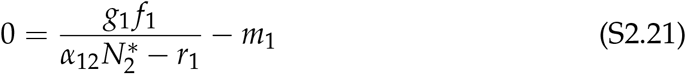

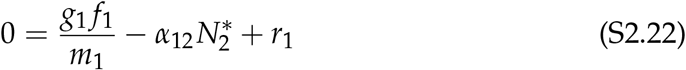

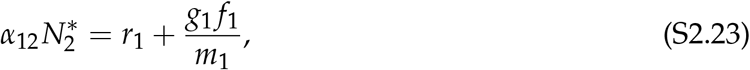

where in the second line we have multiplied through by the quantity 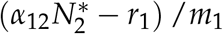. Biologically, we note that increasing the term on the left-hand side corresponds to increasing competitive pressure from residents, and therefore harms the invader’s growth, while increasing the terms on the right-hand side benefits the invader (e.g., increasing its reproduction and maturation, or decreasing its mortality). Accordingly, we conclude that the invasion condition is

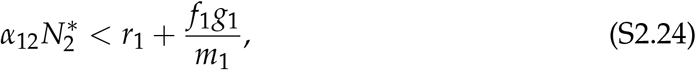

matching the result from the full calculation.

Alternatively, we can reason about the inequality from the mathematical properties of the invasion growth rate expression. We can write the invasion growth rate condition in *λ* (eq. S2.20) as *p*(*λ*) = 0, where

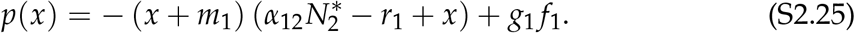

Here, *p*(*x*) is a quadratic polynomial with negative leading coefficient, such that *λ* is the biologically meaningful root of *p*. We note that the only biologically relevant root of *p* is the one corresponding to greater *Q*\* (as shown above) and thus is the greater of the two possible roots. Therefore, due to *p*’s negative leading coefficient, a positive value of *p*(*x*) for some value *x* implies that the root *λ* is greater than *x*. Thus, *p*(0) > 0 implies *λ* > 0 and means that species 1 can successfully invade, which we can rearrange (noting that 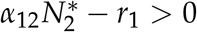 near the invasion boundary) to give the inequality for the invasion condition above.

#### S2.1.3 Derivation with alternate variable transformations

To illustrate that there are multiple valid options for *N* and *Q*, we use the alternate and somewhat more intuitive transformations, as mentioned in the main text:

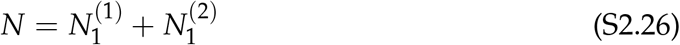

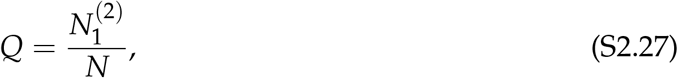

where *N* is the total abundance of both stages, and *Q* is the fraction of that total occupied by the adult stage 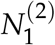. Though the algebra is significantly messier, we find the same results. First, using the transformations 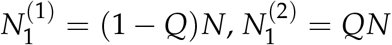, we find that the per-capita dynamics of *N* follow

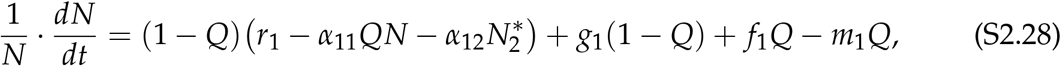

where we can remove the *−α*_11_*QN* term when considering the invasion dynamics (*N ≈* 0). Next, using the chain rule, we find that

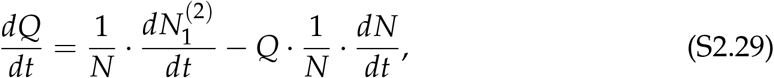

and substituting the expressions for 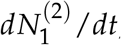, (1/*N*) *dN*/*dt*, and 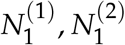, we find (again neglecting the *α*_11_ term due to the invasion condition):

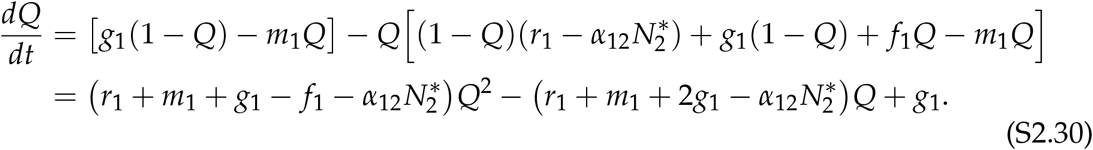

Solving the quadratic, we find that

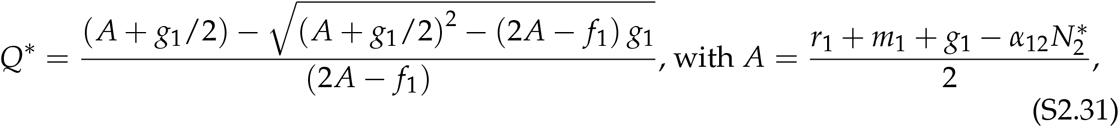

and it is now the root with *−* which is biologically meaningful (since *Q* cannot be greater than 1). Finally, we substitute back into the per-capita growth rate of *N* to find the invasion growth rate. Simplifying 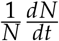 and applying the invasion condition (i.e., dropping the *α*_11_ term as *N ≈* 0 and setting species 2 to the resident equilibrium 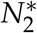), we find:

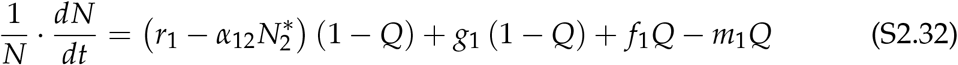

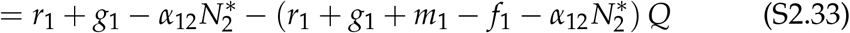

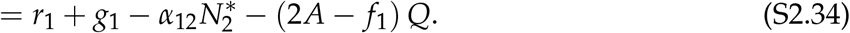

Thus, substituting the quasi-equilibrium stage structure *Q*\*, we find that

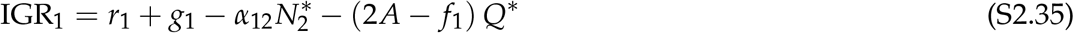

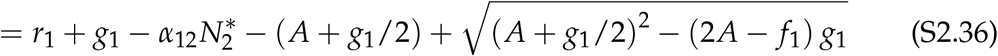

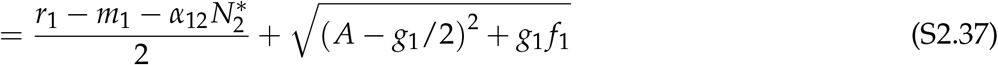

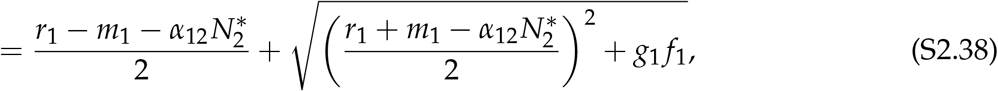

matching eq. S2.11 above. Thus, these alternate transformations give identical results, though the algebra is notably more tedious.

### S2.2 Method 1.2: Strategic linear combination

Let 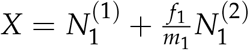. Then

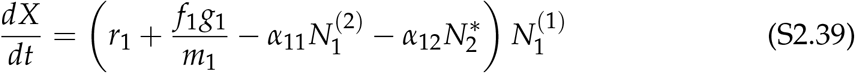

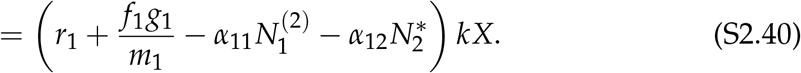

Then the invasion growth rate is

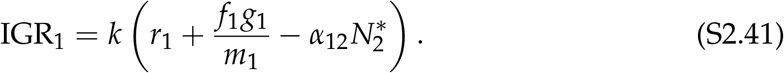

Given that *k* is a positive constant, the criterion for a positive IGR matches the expression in eq. S2.15:

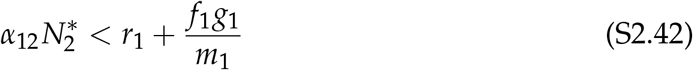

### S2.3 Method 2.1: Direct solution for leading eigenvalue

Via linear algebra, we consider the eigenvalues of the matrix

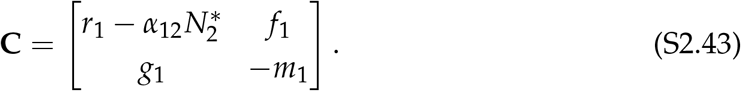

Solving the characteristic equation det (**C** *− λ***I**) = 0, we obtain

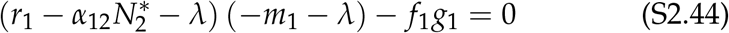

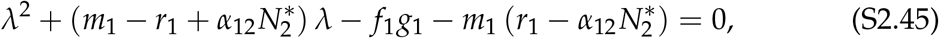

which has the leading eigenvalue:

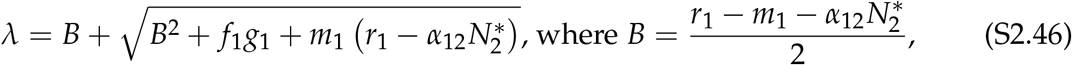

and we note that, following the logic in Method 1.1, *λ* is positive when the term 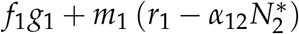 is positive, which rearranges to the same expression in eq. S2.15:

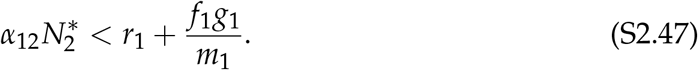

### S2.4 Method 2.2: Applying stability conditions

Applying the Routh–Hurwitz stability conditions, we find that *λ* is positive if one of the following conditions holds. First, a negative determinant implies exactly one eigenvalue with a positive real part,

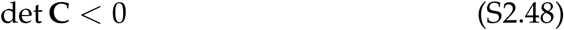

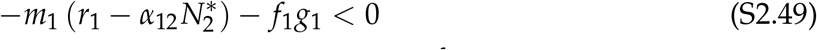

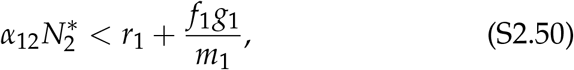

which matches the expression in eq. S2.15. Second, if this condition does not hold, the eigenvalues share a sign and a positive trace implies that both eigenvalues are positive:

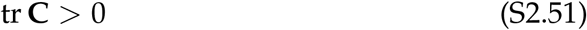

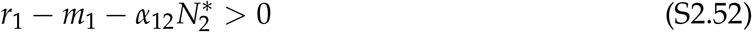

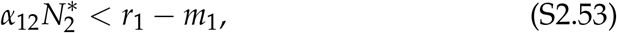

but as we noted above, this condition also implies the first.

## S3 General numerical recipes

### S3.1 Simulation-based algorithms for invasion growth rates

#### Algorithm S3.1.

Direct simulation for invasion growth

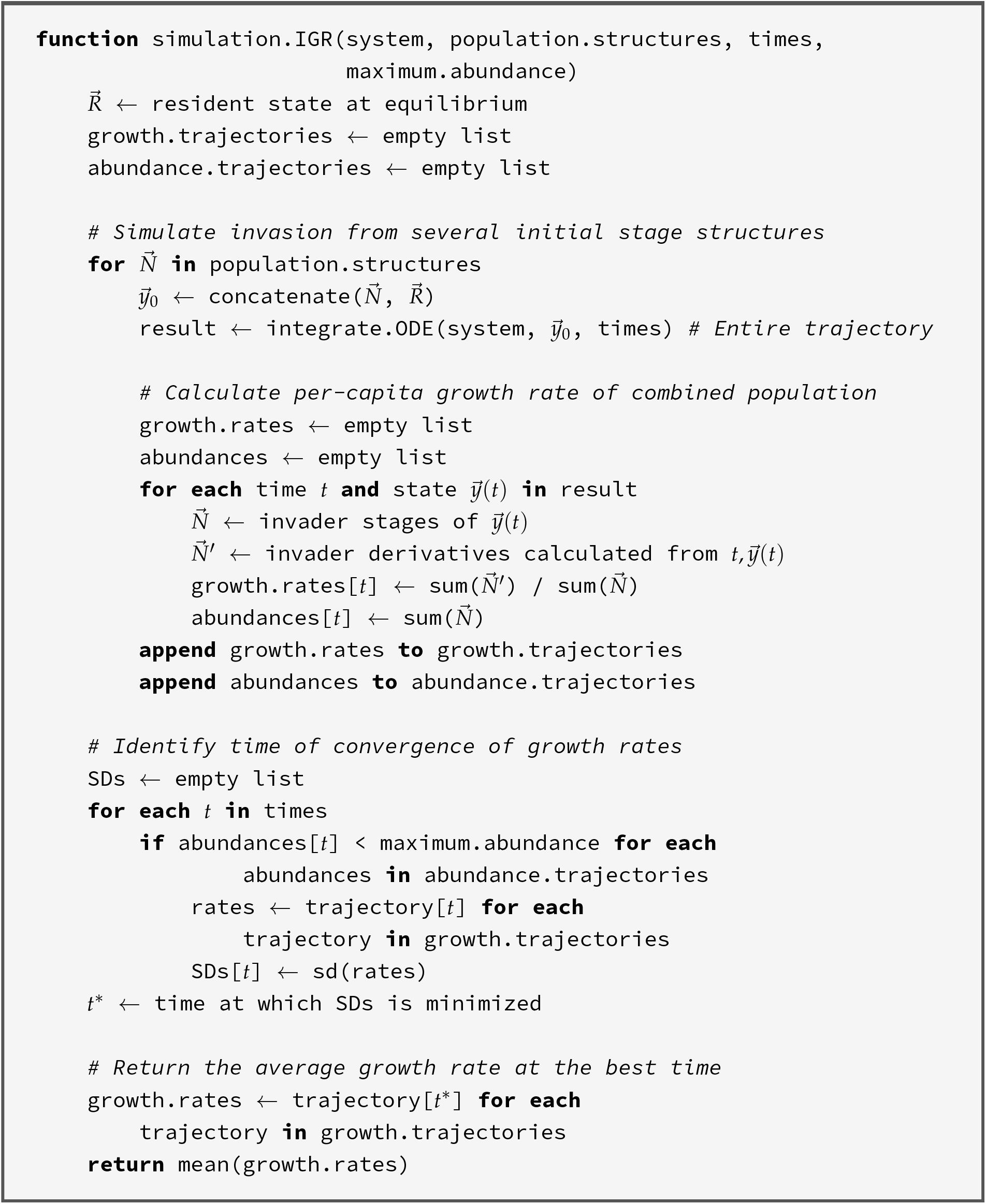

#### Algorithm S3.2.

Rescaling (Lyapunov exponent) algorithm for invasion growth

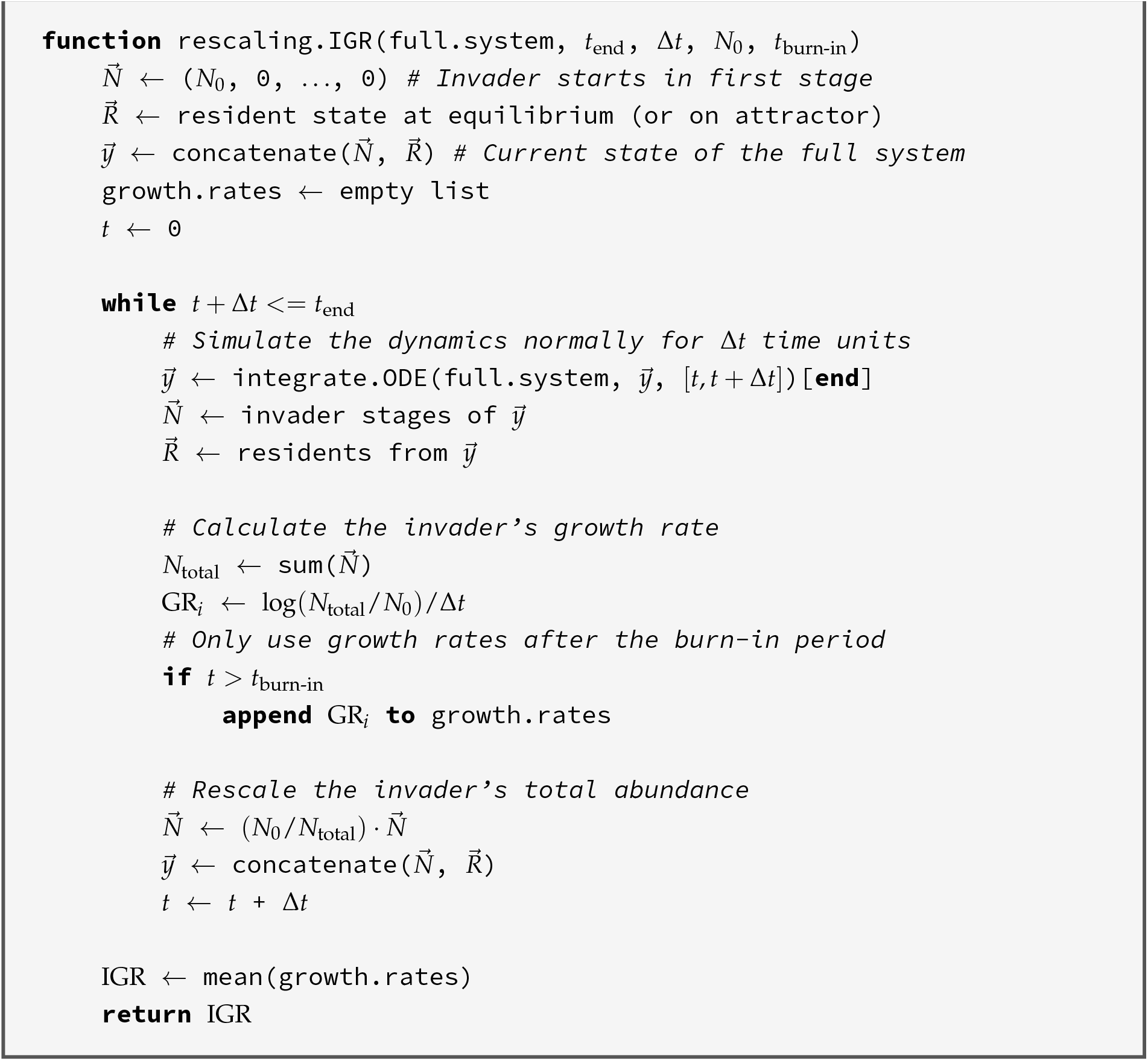

### S3.2 Linearization of invader dynamics

#### Algorithm S3.3.

Linearization of invader dynamics

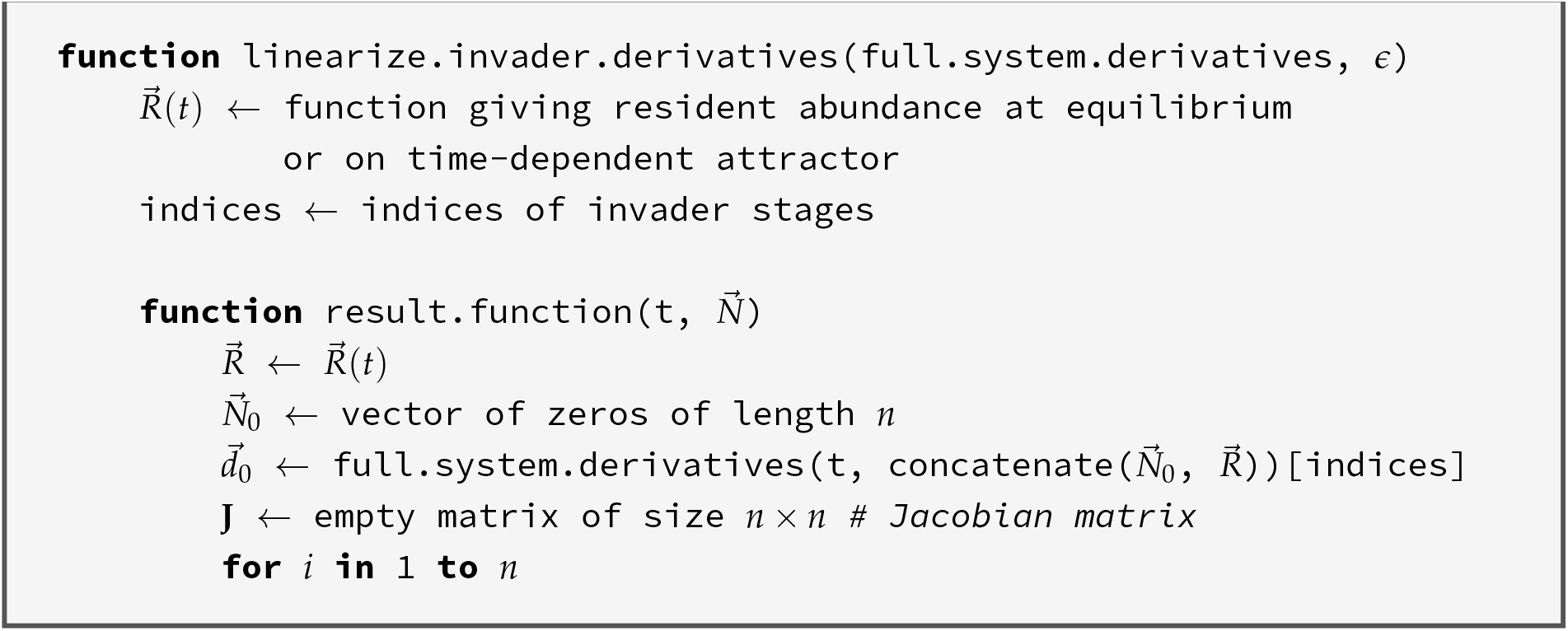
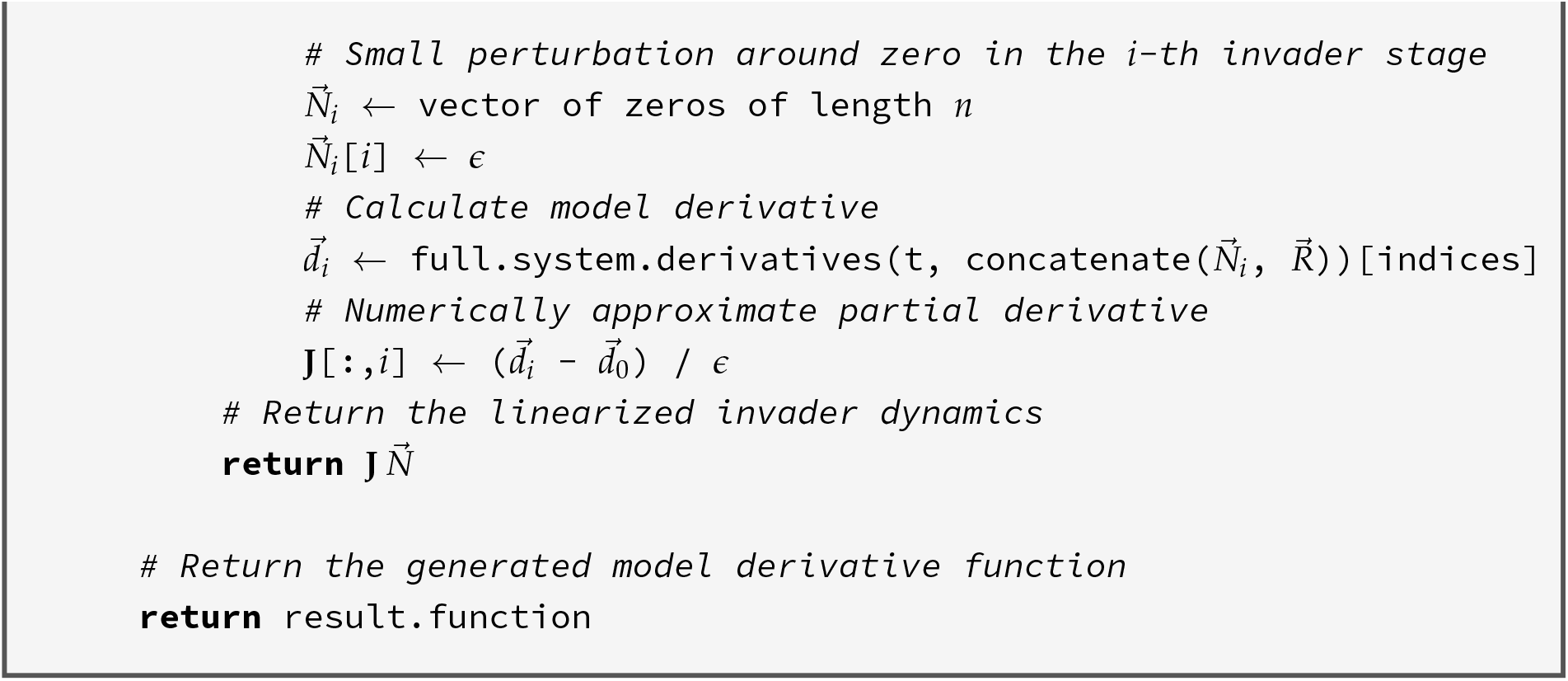

#### Algorithm S3.4.

Invasion growth with linearized invader dynamics

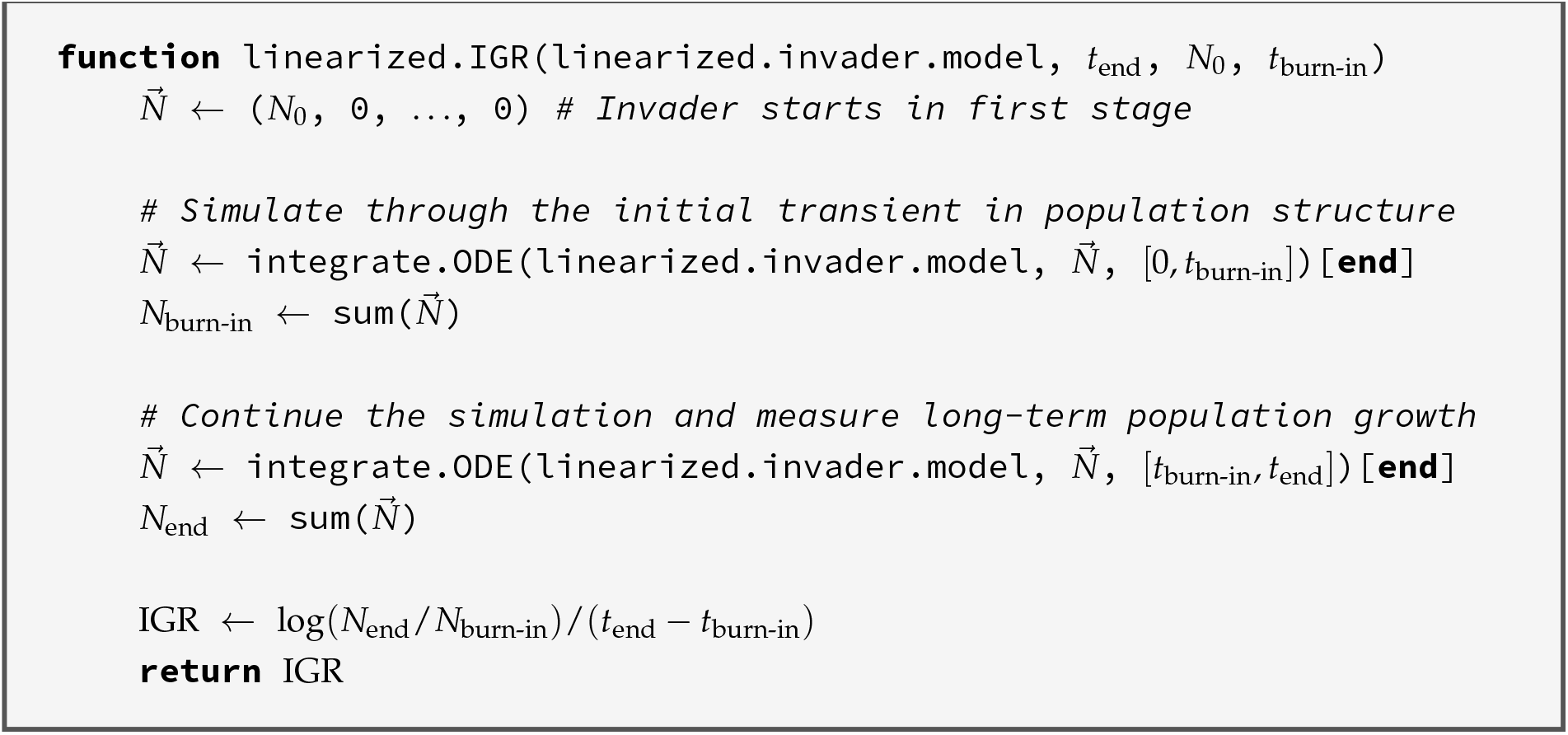

### S3.3 Floquet theory

#### Algorithm S3.5.

Monodromy matrix

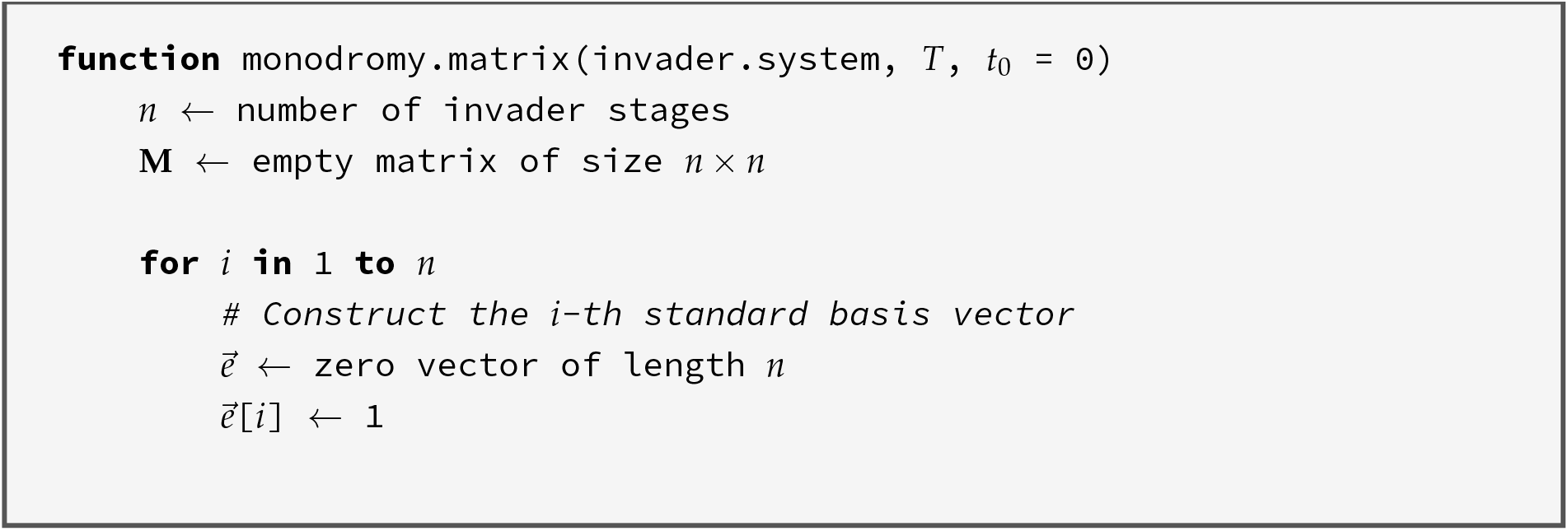
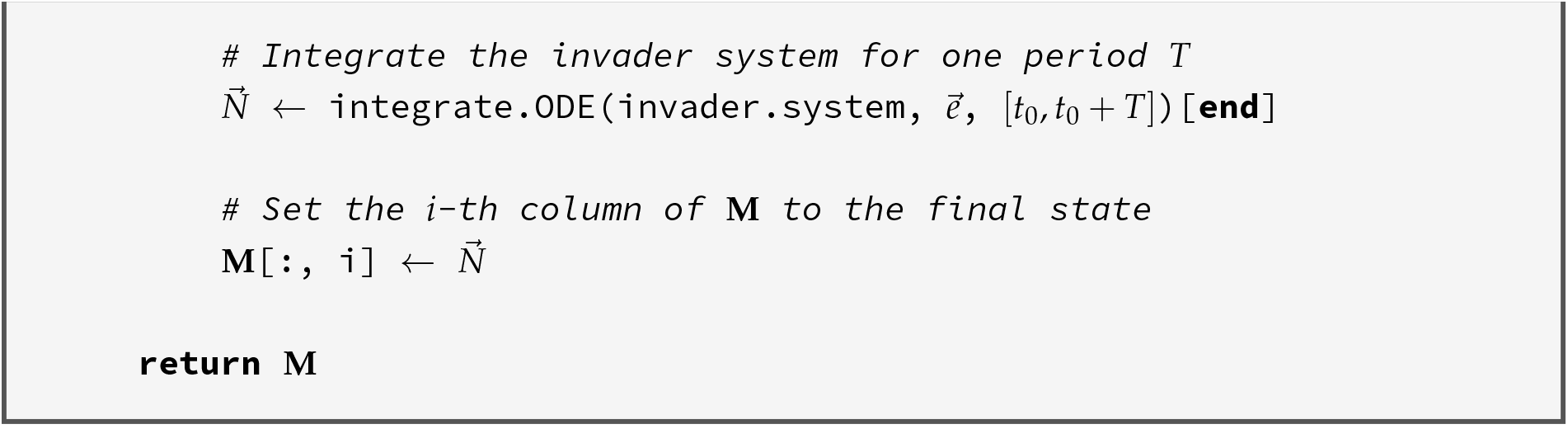

#### Algorithm S3.6.

Floquet calculation of invasion growth

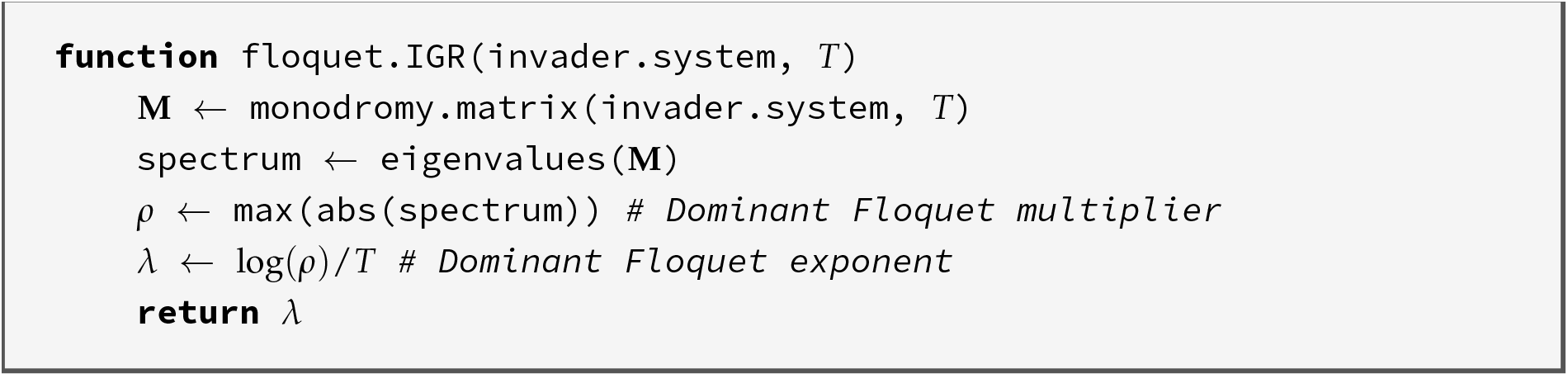

#### Algorithm S3.7.

Finding the Floquet exponent matrix B

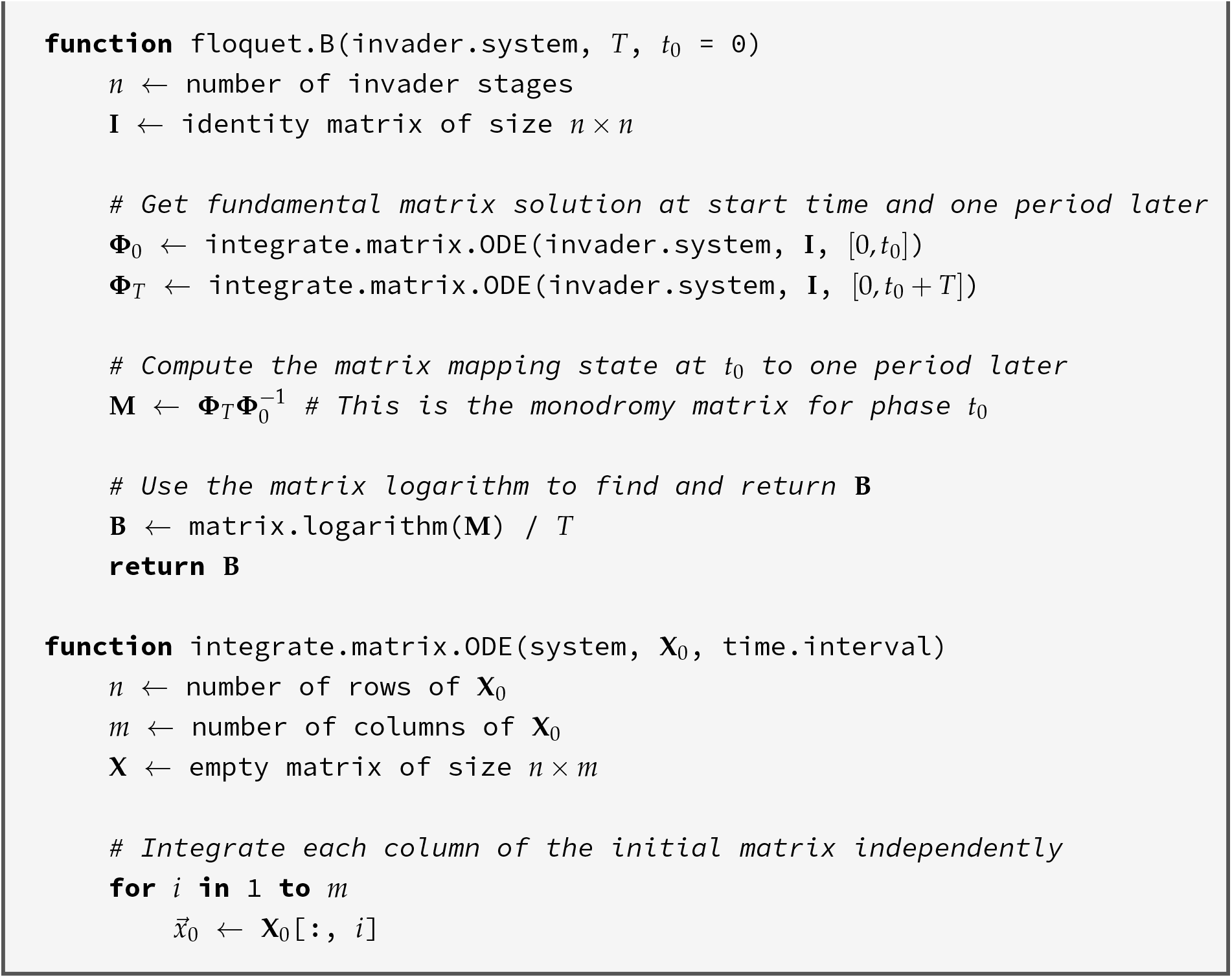
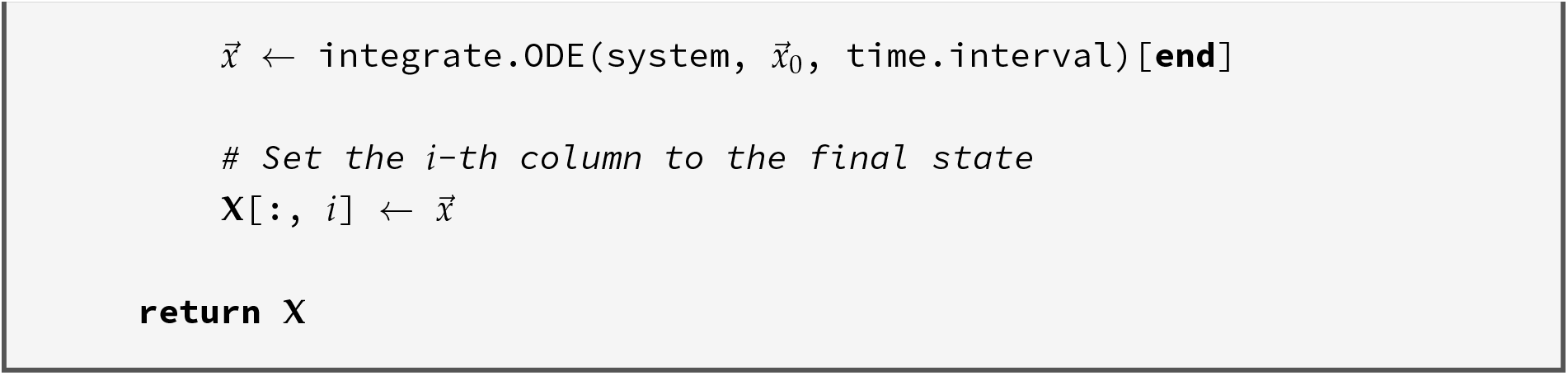

#### Algorithm S3.8.

Periodic Floquet transformation

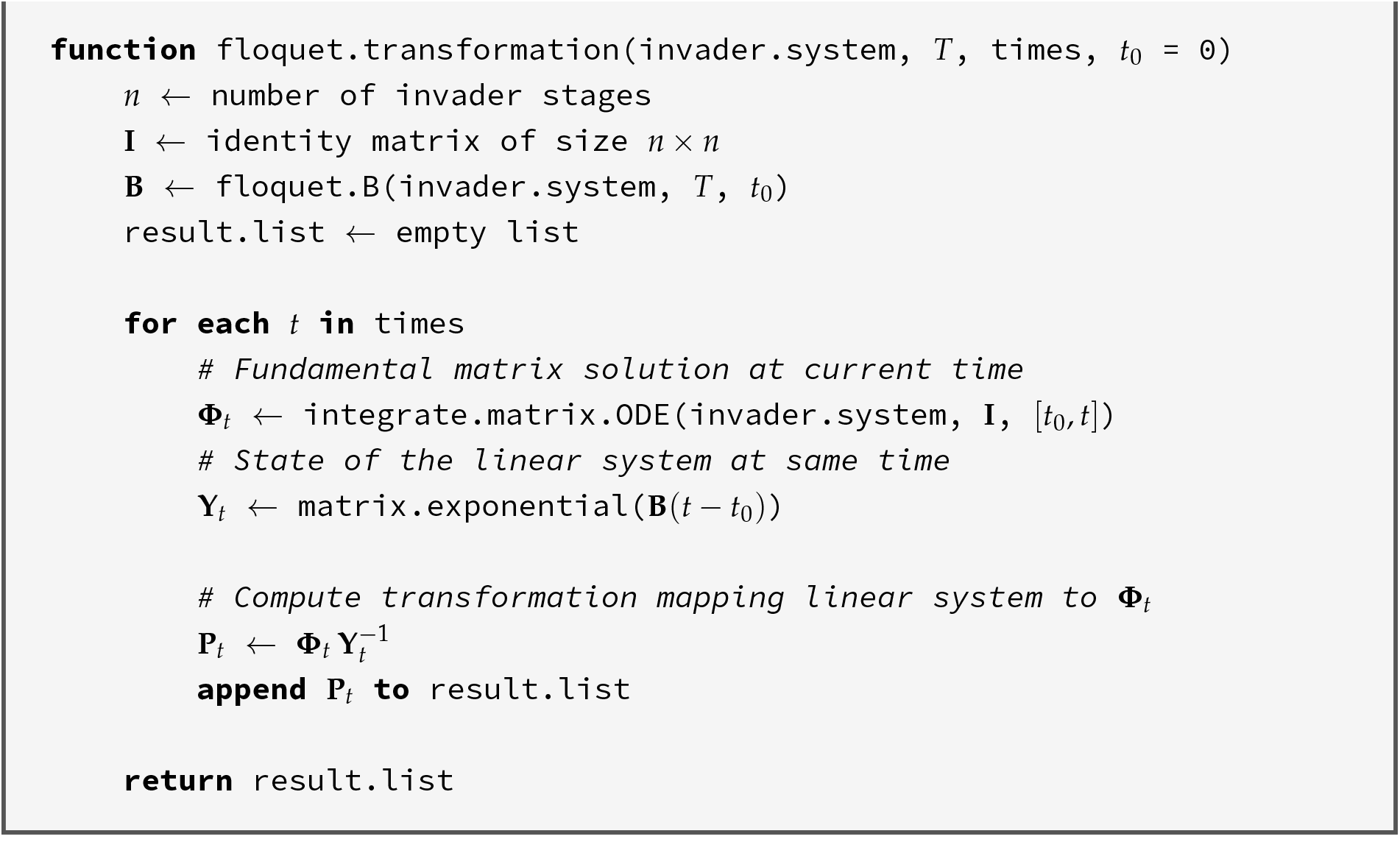

## S4 Intraguild predation and cannibalism model

We consider a set of insect populations with three stages 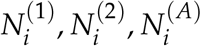 competing for a shared logistically growing resource *R*.

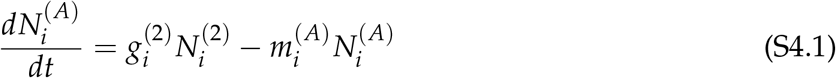

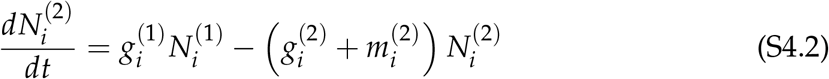

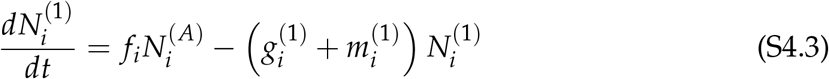

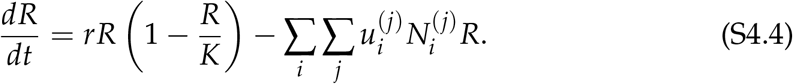

In this general model, assume development 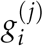, mortality 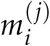, and fecundity *f*_*i*_ are functions of the state variables (superscript *j* = 1, 2, *A*). In particular, they may be coupled to resource uptake 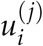 *R*. On the other hand, stages may die due to predation by other stages, or the development of predatory stages may change as a result of their consumption of prey stages.

### Three-stage intraguild predation model

From this general framework, we analyze a specific version of the model with intraguild predation (IGP) in two species whose development depends on resource uptake. Assume species 1’s second juvenile stage 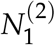 performs predation at density-dependent rate *a*_*i*1_ on first-stage juveniles 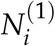 of both species (*i* = 1, 2, representing cannibalism and IGP, respectively) which (i) affects the mortality of the predator individual, and (ii) causes mortality of the prey individual. Thus, the mortality of the first stage of both species is given by a baseline mortality 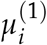 plus mortality due to predation by 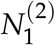 ,

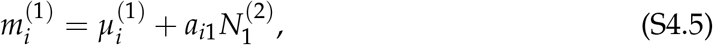

where *i* = 1, 2. Furthermore, for the predatory stage 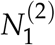, assume that its mortality depends not only on resource uptake 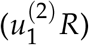 but also on the rate at which it predates prey species *i*, i.e., 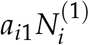 according to the following function

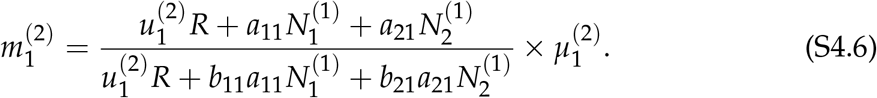

Now, consider resource dynamics. Assume that juvenile states’ development rates 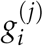 increase linearly with resource level according to the uptake constant 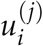. Then,

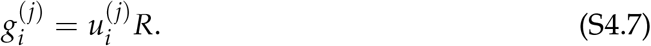

Finally, we assume fecundity is a constant rate *f*_*i*_.

Putting these together, the dynamics of the two species, where species 2 is preyed upon by its competitor but does not itself engage in predation, are given by:

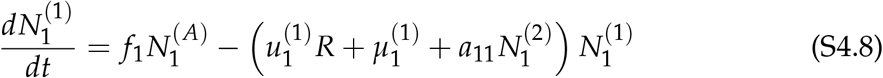

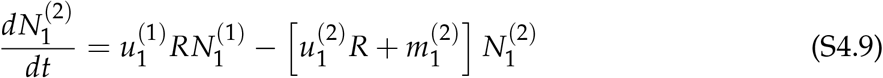

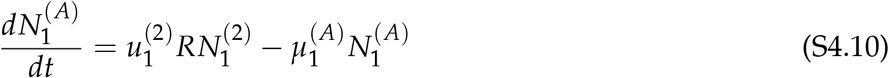

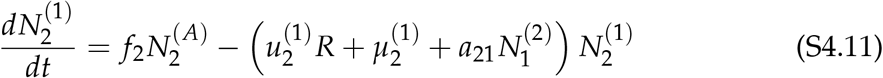

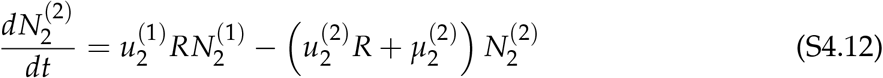

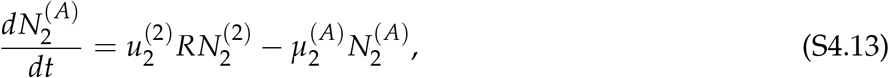

with 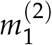 as the function defined above in eq. S4.6, and resource dynamics given by the same equation as in the general model above.

### Two-stage simplification

Note that the dynamics of adults are relatively simple: assuming the other variables are at equilibrium or changing slowly, adult abundance of species *i* reaches the quasi-equilibrium value,

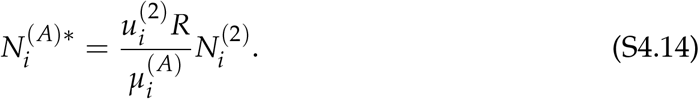

Then, with the assumption that adult population dynamics are fast, we can simplify the dynamics of the two species to:

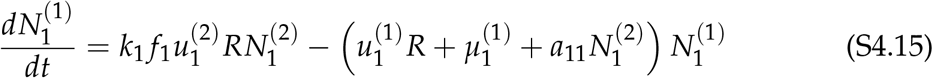

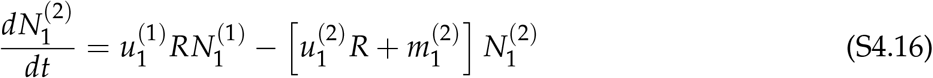

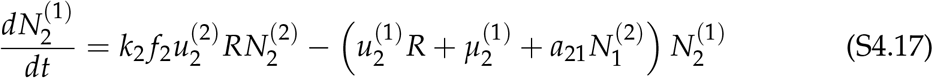

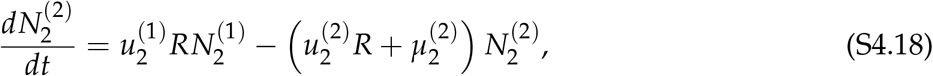

where 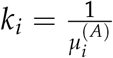 and 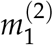 is a function of resource and 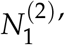 s consumption of first-stage juveniles (eq. S4.6); resource dynamics are as previously given.

### Monoculture equilibrium of species 2

Consider the invasion of species 1 (the predator species) into a monoculture of species 2 (the prey). We first solve for the monoculture equilibrium of species 2 alone, thereby ignoring the 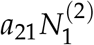 term. The resulting equilibrium (*R*\*, 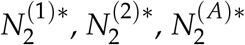) fulfills the following:

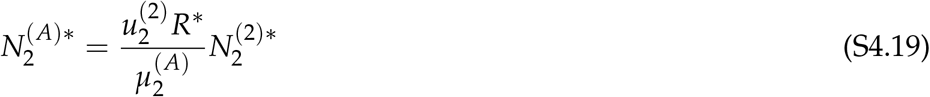

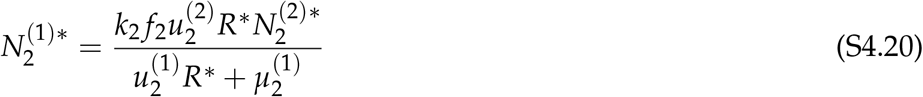

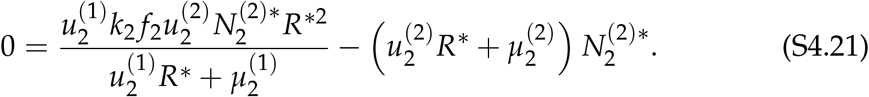

Dividing the third equation above by 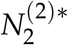 and multiplying by the denominator, we arrive at:

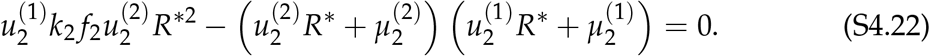

### S4.1 Method 1.1: Direct timescale separation

We will choose a new pair of variables: *N*, representing the slow dynamics of population growth, and *Q*, representing the fast dynamics of population stage structure. Since direct predation only depends on the second juvenile stage, we choose 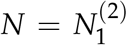 as a convenient choice for the population (slow) variable. Accordingly, a convenient choice of the stage structure (fast) variable is the ratio of first-to second-stage juveniles, 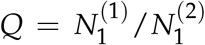. Then, 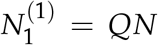, and we can rewrite the system considering the invasion scenario:

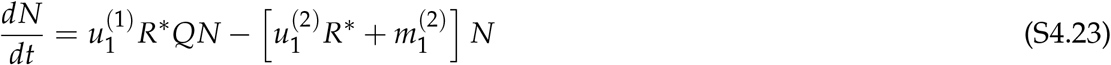

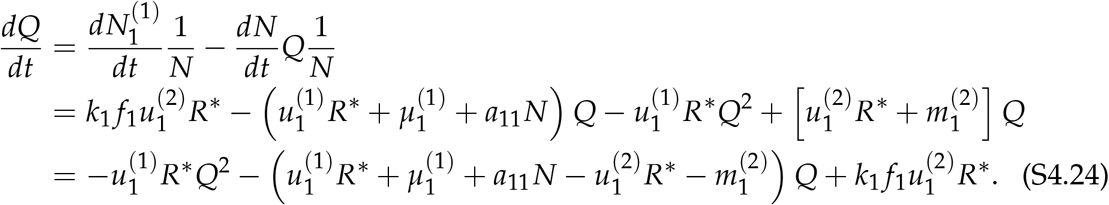

We proceed to solve for the quasi-equilibrium of *Q* in the invasion condition. In this case, *N ≈* 0, allowing us to neglect the term *a*_11_*N* (i.e., the cannibalism term representing the consumption of first-stage juveniles by second-stage juveniles). Solving using the quadratic formula gives the single biologically meaningful root,

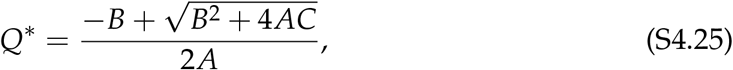

where

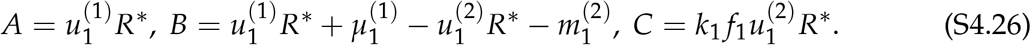

We note that the other root is necessarily negative and therefore not biologically meaningful (since *Q* is a ratio of populations). Next, we find the per-capita growth rate of *N* when *Q* = *Q*\*, i.e., the invasion growth rate IGR_1_:

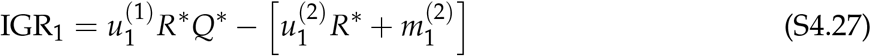

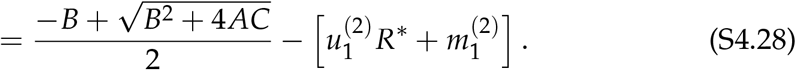

Finally, we solve for the conditions under which the invasion growth rate is positive:

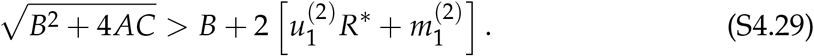

Noting that 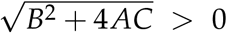 and the right-hand side of the above inequality 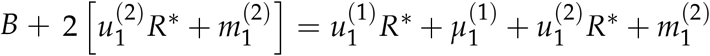 is also positive, we can rewrite:

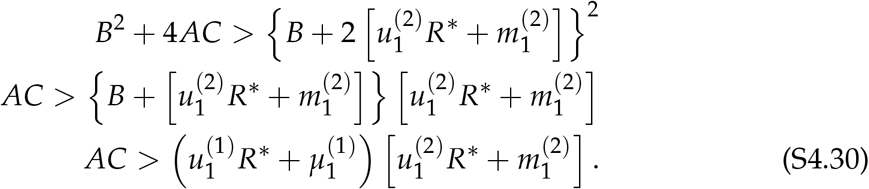

Substituting the expressions for *A* and *C*, followed by subtracting 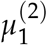, and finally dividing the whole inequality by 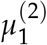, we get:

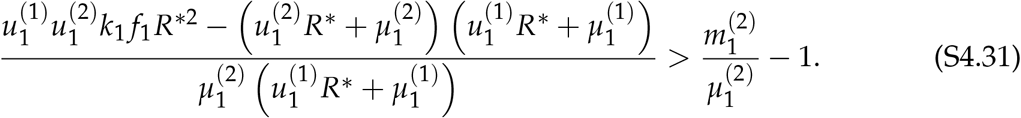

### S4.2 Method 1.2: Strategic linear combination

At the invasion equilibrium, let 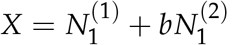, with 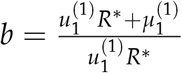. Then:

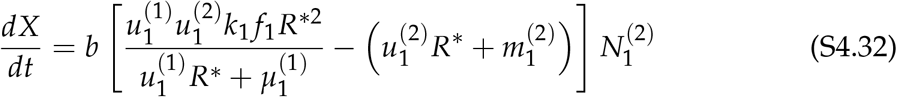

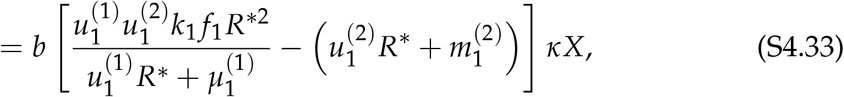

with *κ* > 0. The invasion growth rate is thus 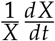, which is positive when:

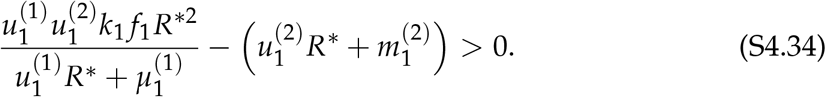

With the same analytical trick as above, this criterion can be written as follows, which is identical to the condition in eq. S4.31:

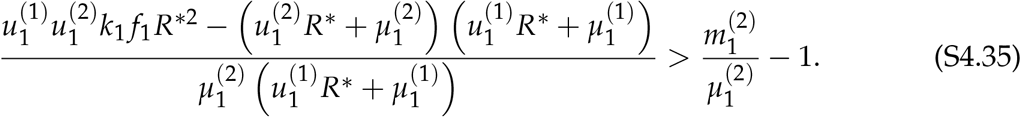

### S4.3 Method 2.1: Direct solution for leading eigenvalue

For the same two-stage simplification as before, the dynamics of species 1 during invasion can be written in the linear matrix form:

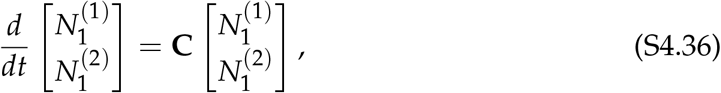

where the matrix **C** is given by (note that the term 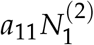 is omitted due to low abundance during invasion):

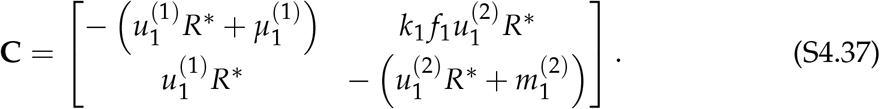

Writing the characteristic equation det (**C** *− λ***I**) = 0, we obtain

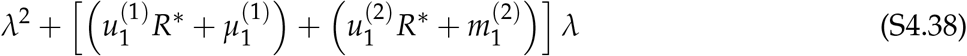

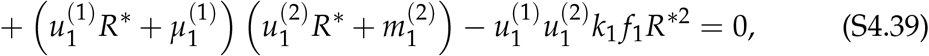

which has the leading eigenvalue,

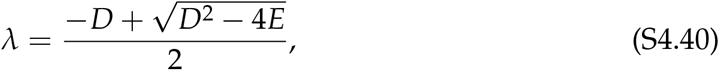

where

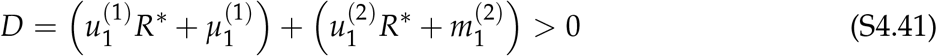

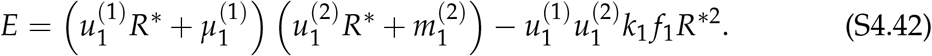

Given that *D* > 0, the leading eigenvalue, and thus the invasion growth rate, is positive if *E* is negative, giving

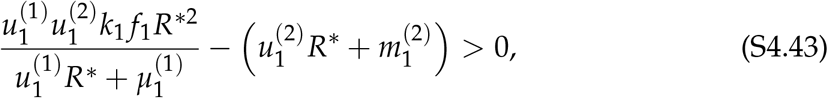

which is identical to the previous invasion criterion in eq. S4.34 and can be further arranged to give eq. S4.31.

### S4.4 Method 2.2: Applying stability conditions

Applying the Routh–Hurwitz stability conditions, we find that *λ* is positive if one of the following conditions holds. First, a negative determinant implies a strictly positive dominant eigenvalue,

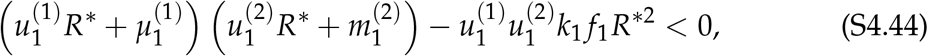

which matches the expression in eq. S4.31 and eq. S4.34. Second, if the determinant is positive, the eigenvalues share a sign and a positive trace implies that both eigenvalues are positive:

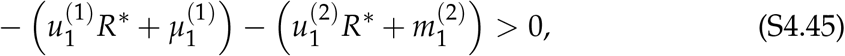

which cannot be fulfilled since all the parameters are strictly positive. Therefore, the determinant criterion solved above is the necessary and sufficient condition for the invasion of species 1.

### S4.5 Parameter effects on invasion

From the invasion criterion in eq. S4.31, we can also derive a pure parameter-based invasion condition by substituting the expression for 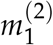 into the invasion condition. Considering the scenario that the two species are identical except for the aggressive behavior of species 1, the numerator of the left-hand side of eq. S4.31, i.e.,

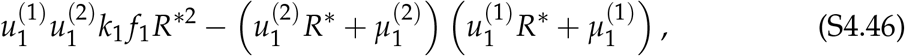

equals

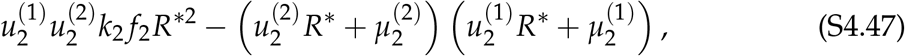

which equals zero given the monoculture condition in eq. S4.22. Therefore, the necessary and sufficient condition for species 1 to invade is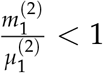, which can be written out in full at invasion (i.e., 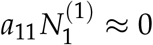) as:

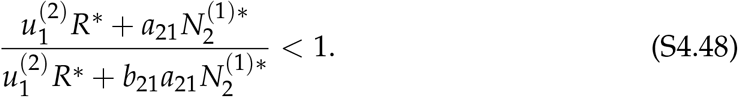

Therefore, if IGP is beneficial to species 1 (*b*_21_ > 1), which reduces the mortality of the predator, species 1 can always invade. Alternatively, if IGP is not beneficial (e.g., species 2 is toxic and increases species 1’s mortality), then species 1 cannot invade. Importantly, one can also note that cannibalism does not affect the invasion criterion, which is understandable as cannibalism is inconsequential when species 1’s abundance is low.

## S5 Plant–soil feedback model

We consider a simplification of the patch occupancy model used by Ke & Levine (2021) to study the effects of soil legacy persistence on plant competition. As in their “fast conditioning” models, we consider the transition to the conditioned state to be instantaneous when a plant grows on unconditioned soil (termed “conditioning” for short). However, we allow the transition to the conditioned state to be slow when a plant grows on a patch which already has a conditioned microbial community (termed “reconditioning”). This reflects the assumption that reconditioning should take longer than conditioning because it requires regression of the existing microbial legacy in addition to conditioning of the plant’s own microbial community.

Our model treats seven state variables, representing the dynamics of the proportion of patches occupied by combinations of plants and soil states. Unoccupied patches may host no microbial community 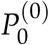 or be conditioned by either plant species 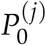 (where *j* = 1, 2). However, patches with living plants (*i* = 1, 2) always have a conditioned community (*j* = 1, 2), and patches with the soil community of the opposite species 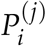 (*j≠ i*) transition to patches where the plant species has conditioned its own community 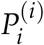. Thus, the model can be written as the system of differential equations

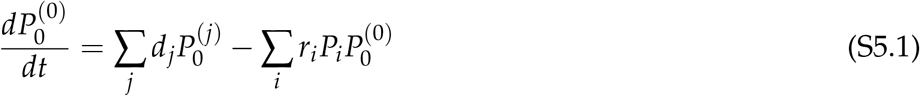

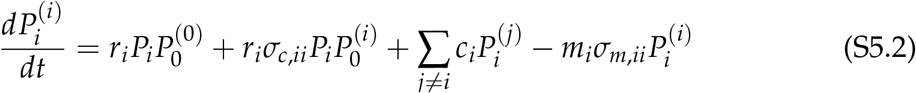

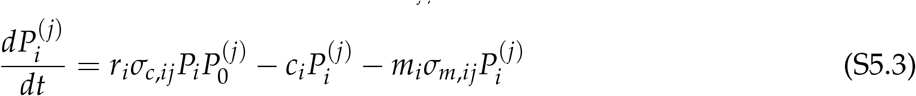

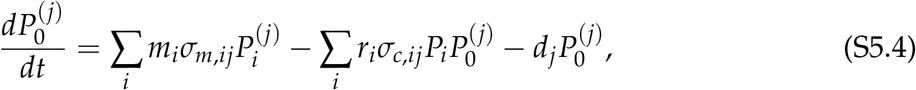

where *i,j* ∈ {1,2}.

Here, *r*_*i*_ is the colonization rate of plant *i*, which is modified by a colonization feedback factor of *σ*_*c,ij*_ for patches with microbial community *j≠* 0. Colonization furthermore depends on the total fecundity of species *i*, which we define as 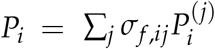, where *σ*_*f* ,*ij*_ is the fecundity feedback factor of plant *i* when growing in soil *j*. Upon colonization, unconditioned patches 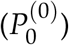 immediately transition to the 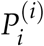 state, while patches with conditioned microbial communities 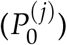 transition to the 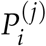 state. Meanwhile, plants in occupied patches die at a mortality rate *m*_*i*_, which is modified by a mortality factor *σ*_*m,ij*_. Finally, plants of species *i* gradually recondition soil to their own community at a rate *c*_*i*_, while unoccupied patches of soil type *j* (i.e., 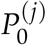) slowly decay to unconditioned soil 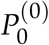 at a rate *d*_*j*_.

Written out in full for two species, the model is:

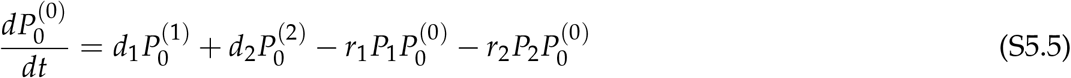

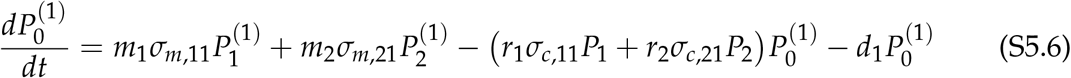

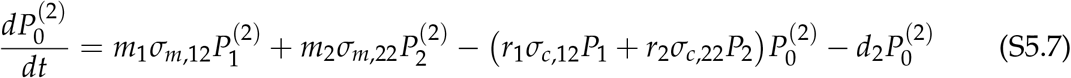

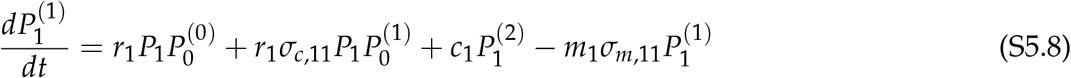

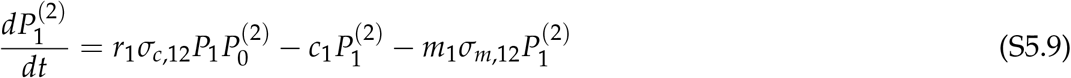

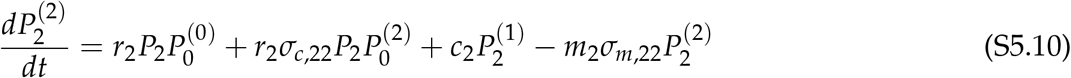

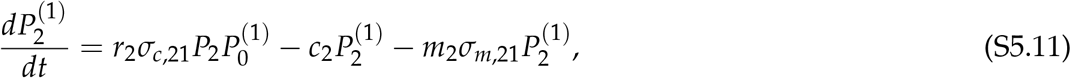

where the fecundity-weighted total abundance terms *P*_1_, *P*_2_ used for calculating colonization are given by

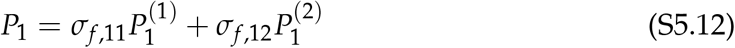

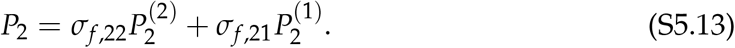

Note that we have written *c*_*i*_ instead of *c*_*ij*_ because there is only one reconditioning rate per species.

### Monoculture equilibrium

We first solve for the monoculture equilibrium of plant *I* growing alone. Dynamics reduce to:

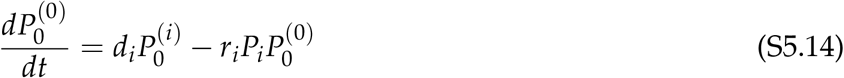

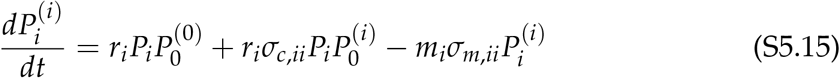

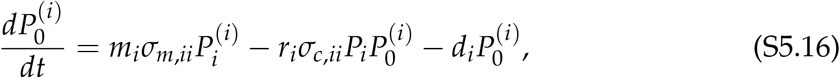

where we have dropped any terms involving other species *j*. At equilibrium,

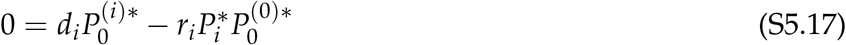

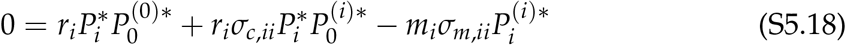

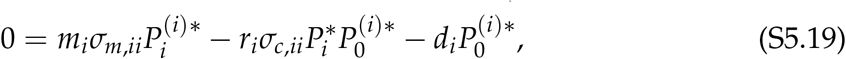

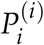 can be factored out of 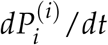, so we solve ^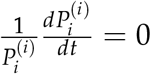^ for 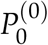 to find:

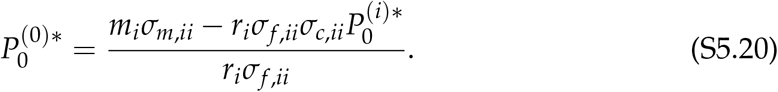

Letting *β*_*i*_ = *σ*_*m,ii*_/*σ*_*m,ij*_, *γ*_*i*_ = *σ*_*f* ,*ii*_/*σ*_*f* ,*ij*_, 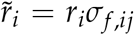 and 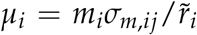, with *j≠ i*, we can simplify this to:

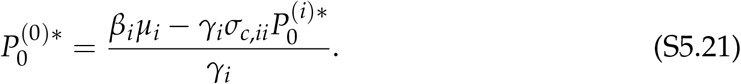

Using 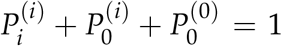, and letting 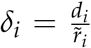, we can set 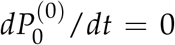 to obtain a quadratic equation for 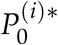 :

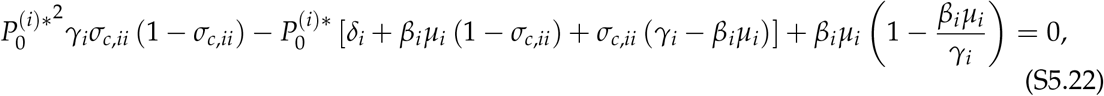

which is a quadratic function of 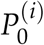. To ensure feasibility, we constrain *r*_*i*_*σ*_*f* ,*ii*_ > *m*_*i*_*σ*_*m,ii*_ in further calculations. Substituting 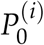 with 0 gives 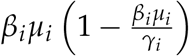, which is positive, and substituting 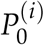 with 1, we obtain 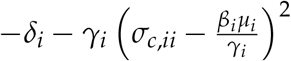, which is negative. Thus, the quadratic function must have exactly one root in the interval (0, 1). Assuming

negative colonization feedback (*σ*_*c,ii*_ < 1), the polynomial is a concave-up quadratic equation, with the biologically meaningful root

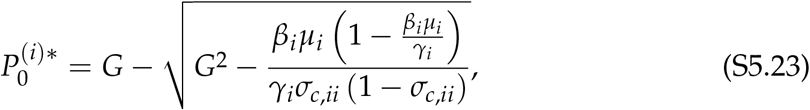

where

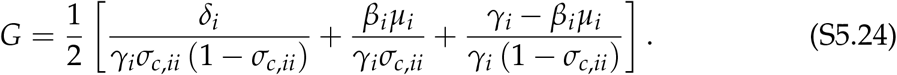

Note that we have taken the root with *−* because there are two positive roots, but only one in the interval (0, 1), so the root with + must be greater than 1 and thus not biologically meaningful. In contrast, when assuming positive colonization feedback (*σ*_*c,ii*_ > 1), the polynomial becomes a concave-down quadratic. Here, we should take the root with + since there is only one positive root, and the criterion makes *γ*_*i*_*σ*_*c,ii*_ (1 *− σ*_*c,ii*_) negative, so the root with *−* must be negative.

### S5.1 Method 1.1: Solve for the invasion growth rate using the *λ* trick

The easiest way to solve for the invasion growth rate using separation of timescales is to use the “*λ* trick” from the simple model analysis (section S2.1.2). We consider the case where species 1 is invading the resident equilibrium of species 2 (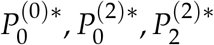). We define the ratio variable:

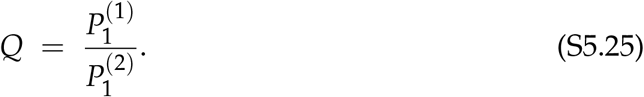

Taking the time derivative gives:

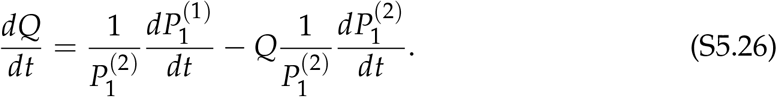

Substituting the dynamical equations for 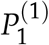 and 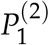 and considering the invasion scenario yields (with 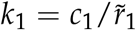):

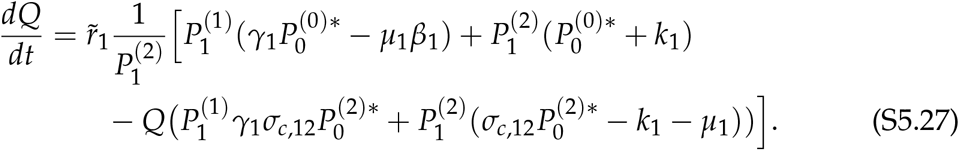

Rearranging terms gives:

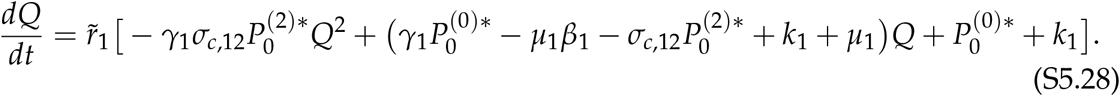

To apply the *λ* trick, we define:

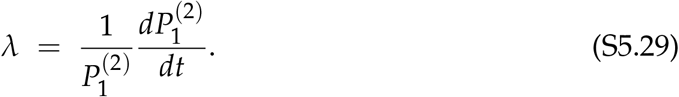

Then the equation for *Q* becomes:

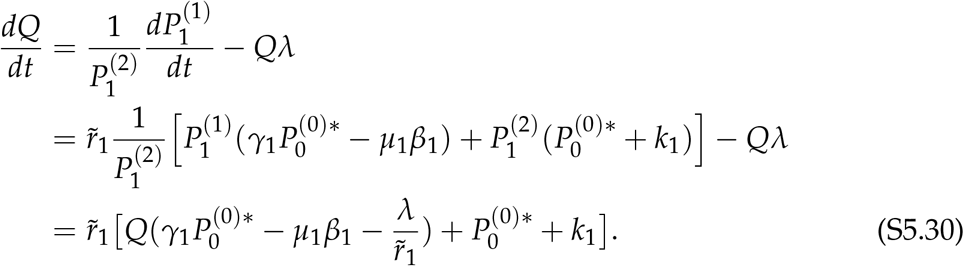

Setting ^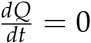^ yields the quasi-steady-state solution

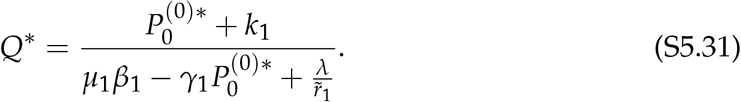

The growth rate *λ* is given by

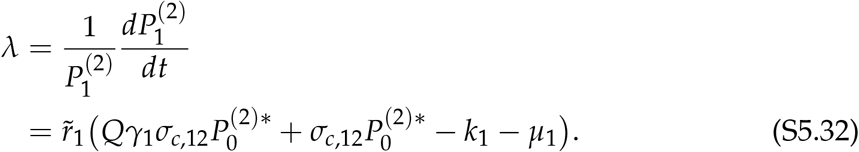

Substituting *Q*\* into *λ* gives

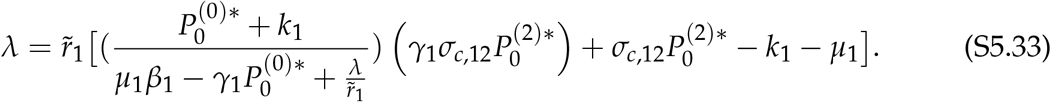

Setting *λ* = 0 at the invasion threshold yields

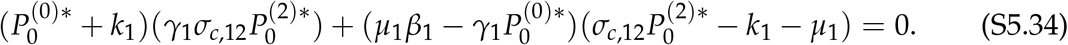

Expanding and rearranging terms leads to the following equation for the invasion threshold:

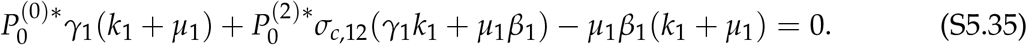

By substituting 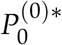 with the relationship in eq. S5.21 and solving the above equation at zero, we can find the critical value of 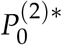 that would be needed to make IGR_1_ = 0 (i.e., 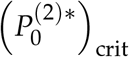):

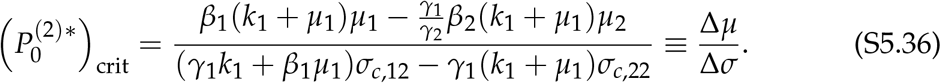

Biologically, we note that soil 2 favors the invader when Δ*σ* > 0 and the invasion condition is 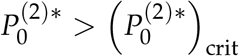 ; alternatively, soil 2 disfavors the invader when Δ*σ* < 0 and the invasion condition is 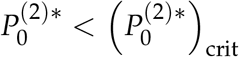. Both cases give the invasion condition for species 1 in terms of the resident equilibrium of species 2.

### S5.2 Method 1.2: Strategic linear combination

Let 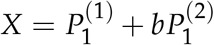, with 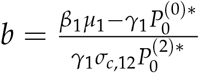. Then, we can write the dynamics of *X* under the invasion scenario as:

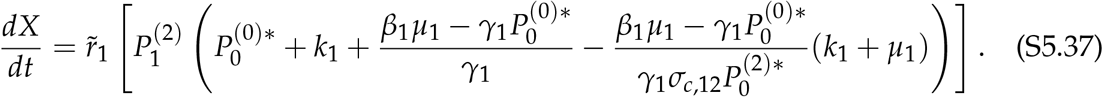

By substituting 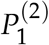 with ^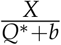^, which is derived from eq. S5.25, we arrive at

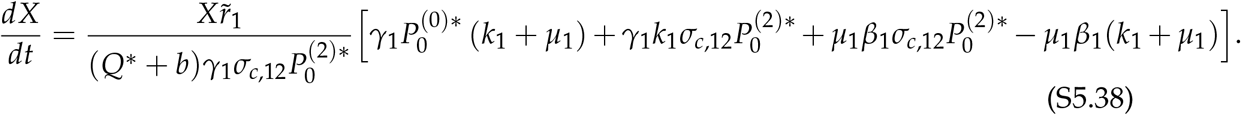

The invasion growth rate is:

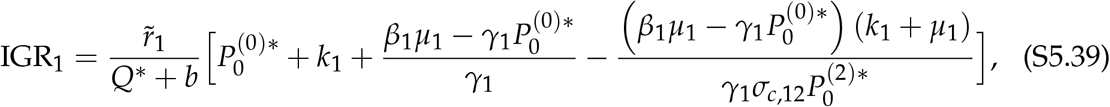

where *Q*\* is obtained from eq. S5.28 as

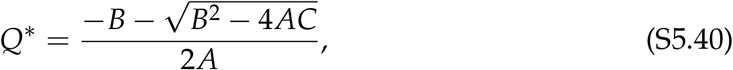

where

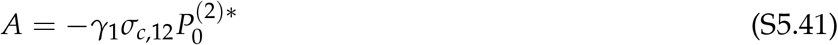

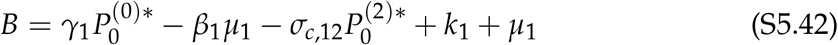

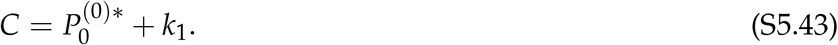

Assuming Δ*σ* in eq. S5.36 is positive, we can derive the condition for IGR_1_ > 0 as:

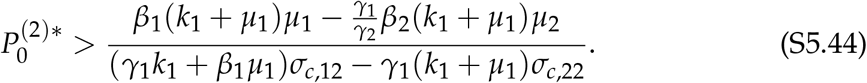

### S5.3 Method 2.2: Applying stability conditions

To determine whether species 1 can invade when rare, we linearize the dynamics of species 1 around the resident equilibrium of species 2. Throughout this analysis, all quantities with a superscript * denote values evaluated at the resident equilibrium. Using the notation introduced above, we define nondimensionalized parameters as follows: *β*_1_ = *σ*_*m*,11_/*σ*_*m*,12_, *γ*_1_ = *σ*_*f* ,11_/*σ*_*f* ,12_, 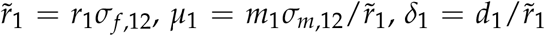, and 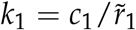. Furthermore, we rescale time to remove a factor of 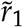 from the dynamics by letting 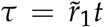. Thus, when species 1 is rare, its dynamics are governed by the linearized subsystem:

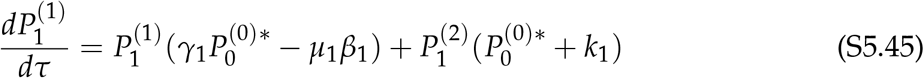

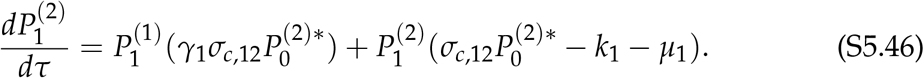

Rewriting these equations in matrix form yields

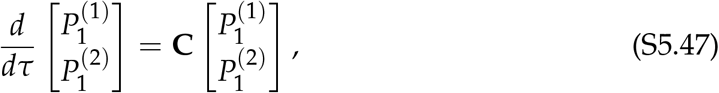

where the Jacobian matrix **C** is given by

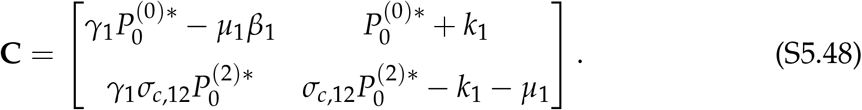

The invasion growth rate of species 1 is determined by the dominant eigenvalue of **C**. Rather than solving explicitly for the eigenvalues, we apply the standard stability conditions for 2 *×* 2 linear systems. In particular, invasion is possible if and only if the dominant eigenvalue of **C** is positive, which occurs when either

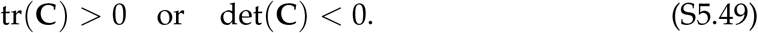

We first consider the determinant:

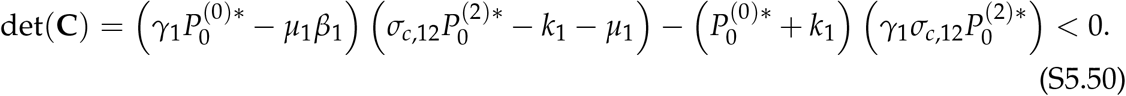

Expanding and rearranging the determinant condition det(**C**) < 0 gives:

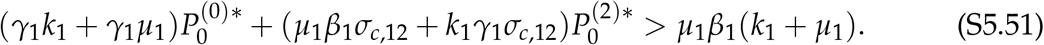

This inequality defines a linear constraint on the resident equilibrium state 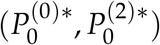 from eq. S5.21, under which the determinant of the Jacobian becomes negative. Biologically, this corresponds to the regime in which the resident equilibrium loses linear stability to perturbations introduced by species 1. Again, assuming Δ*σ* > 0 in eq. S5.36 and solving inequality S5.51 explicitly for the conditioned resident patch fraction 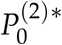 yields the invasion threshold:

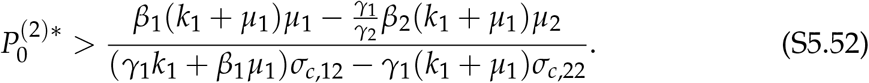

### S5.4 Parameter effects on invasion

From the above invasion criterion, we can also derive a pure parameter-based invasion condition by substituting the equilibrium expressions for 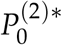 (with square roots) into the invasion condition. However, this is messy to solve. For simpler algebra, consider the boundary where the invader can just invade, i.e., IGR_1_ = 0. As previously mentioned, the critical value of 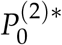 that would be needed to make IGR_1_ = 0 is:

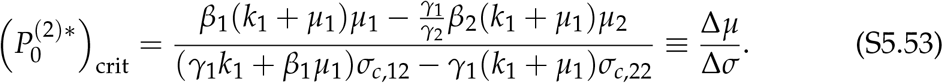

Substituting 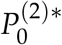 with this critical value into the quadratic expression in eq. S5.22 derived above gives:

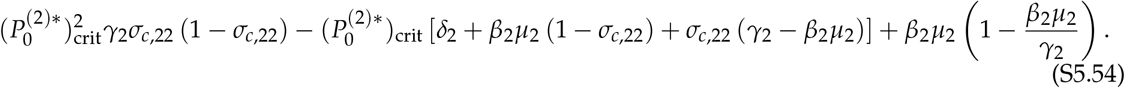

Note that this polynomial has a positive quadratic coefficient as *σ*_*c*,22_ < 1, is positive at 0, and the biologically relevant root is the smaller one. Thus, if the expression is positive at 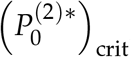, the actual equilibrium 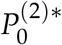 is higher than 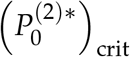, and if the expression is negative, 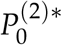 is lower than 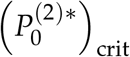. The above two situations correspond to the cases where Δ*σ* > 0 and Δ*σ* < 0, respectively. After some algebra, we can simplify the expression to:

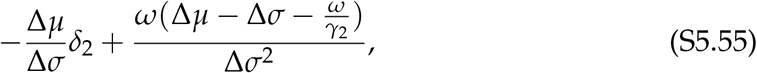

where

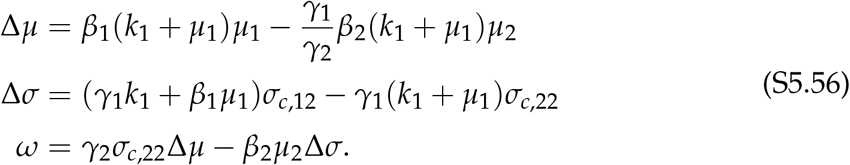

To get the final invasion condition, consider that if soil 2 favors the invader (Δ*σ* > 0), then invasion requires 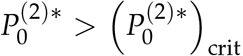, which implies expression S5.55 is positive; multiplying through once by Δ*σ* gives our final answer: species 1 can invade if and only if

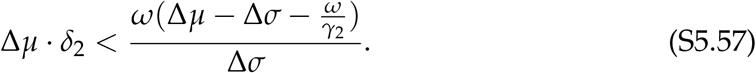

Finally, we note that if soil 2 disfavors the invader (Δ*σ* < 0), then invasion requires 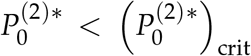 and the expression in eq. S5.55 to be negative. However, since Δ*σ* < 0, multiplying through once by Δ*σ* reverses the inequality and results in the same invasion criterion as above. To gain further insights, we consider the following special cases. First, if we assume that species receive identical microbial effects from both conspecifics and heterospecifics (i.e., *β*_*i*_ = *γ*_*i*_ = 1), the inequality S5.57 can be simplified to:

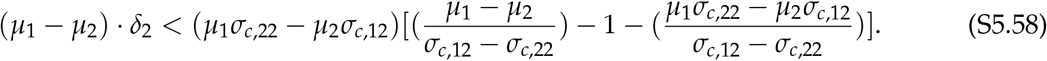

Furthermore, if we assume the realized mortality-to-colonization ratios of species 1 and 2 are identical (*µ*_1_ = *µ*_2_ *= µ*_*i*_), the inequality can be simplified to:

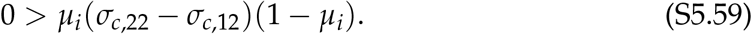

Imposing the feasibility condition, we constrain *µ*_*i*_ < 1 (i.e., *r*_*i*_*σ*_*f* ,*ij*_ > *m*_*i*_*σ*_*m,ij*_) to obtain the final criterion:

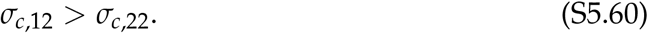

## S6 Plant–pest–parasite model

We model a plant–pest–parasite system where multiple pest species consume a shared host plant *S* and are infected by a common parasite *P*. The parasite infects healthy pests 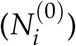 and transitions them to parasitized pests 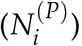, which can survive for a period of time during the infection phase before eventually dying as parasite offspring emerge from their bodies. We assume that the parasite is not vertically transmitted. This system can be described as follows:

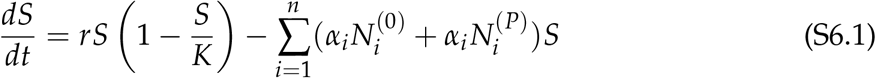

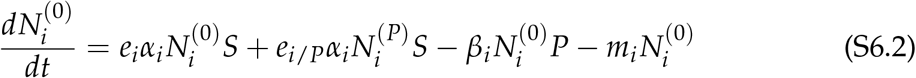

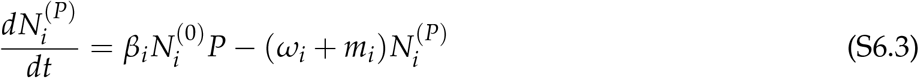

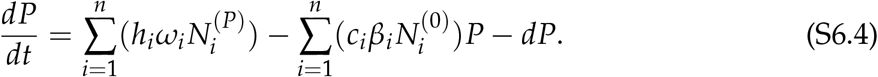

The plant (*S*) grows logistically and is consumed by uninfected pests (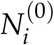; we consider two pest species, so *i* = 1, 2) and parasitized pests 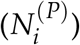 with consumption efficiency *α*_*i*_. As the parasite is not vertically transmitted, host plant consumption by both pest statuses contributes to the uninfected pest population size (with conversion efficiencies *e*_*i*_ and *e*_*i*/*P*_, respectively). Pest population decreases due to natural mortality (with intrinsic mortality rate *m*_*i*_) and parasite infection (with efficiency *β*_*i*_), converting pests into the parasitized state. In addition to their natural mortality, parasitized pests experience an additional parasite-induced mortality *ω*_*i*_. Finally, the free-living parasite population increases following the death of parasitized pests (*h*_*i*_ parasites emerge upon the death of a single parasitized pest), and decreases as they infect pests (with conversion parameter *c*_*i*_) and when they naturally die (with mortality rate *d*).

### S6.1 Invasion analysis for parasite

In this host–pest–parasite system, there are two stage-structured population dynamics: one involves the transition of parasite stages (i.e., free-living versus inhabiting within the pest body) and the other involves the transition of the pest stages (i.e., uninfected versus infected). In this section, we will perform the invasion analysis for the first stage-structured population dynamics; namely, the parasite can be either (i) free-living, represented by the original state variable *P*, or (ii) within the pest body, represented by the original state variable 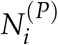. To be consistent with the notation used throughout this study, we represent the free-living parasite as *P*^(0)^ and the pest-infecting stage as *P*^(*N*)^. For simplicity, we consider the invasion of the parasite into a host–single pest equilibrium (*S*\*, *N*\*) and rewrite the system as follows (note we rename *N*^(0)^ as *N* to avoid cluttering the notation with too many superscripts):

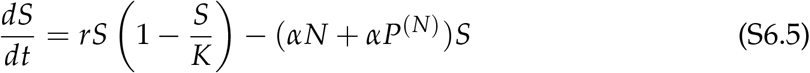

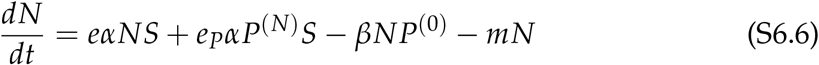

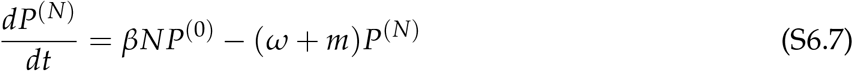

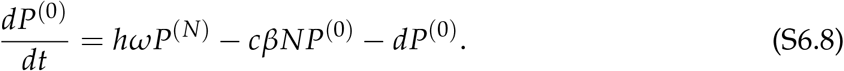

#### S6.1.1 Method 1.1: Direct timescale separation

We will choose a new pair of variables: *P*, representing the slow dynamics of population growth, and *Q*_*P*_, representing the fast dynamics of population stage structure. We choose *P* = *P*^(0)^ for the population (slow) variable. The stage structure (fast) variable is the ratio of parasitized pests to free-living parasites, *Q*_*P*_ = *P*^(*N*)^/*P*^(0)^. Then *P*^(*N*)^ = *PQ*_*P*_, and we can rewrite the system under the invasion condition as

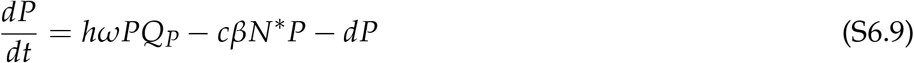

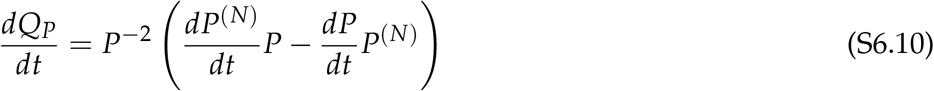

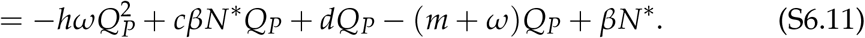

First, we solve for the quasi-equilibrium of *Q*_*P*_ in the invasion condition. Solving using the quadratic formula gives the single biologically meaningful root,

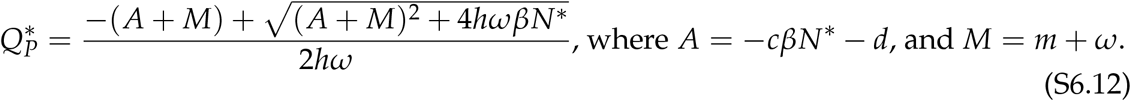

Next, we find the per-capita growth rate of *P* when 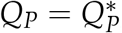, i.e., the invasion growth rate for the parasite:

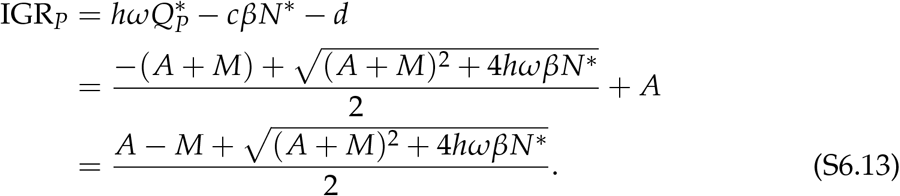

Finally, we solve for the conditions under which the invasion growth rate is positive:

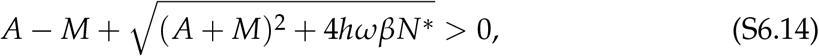

and we note that *M − A* is always positive, from which the following inequality can be derived:

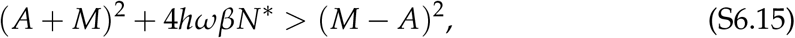

and we get the condition for the parasite to invade:

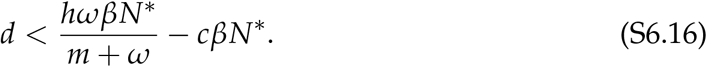

### S6.1.2 Method 1.2: Strategic linear combination

Let *X* = *P*^(0)^ + *bP*^(*N*)^, with 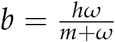. Then

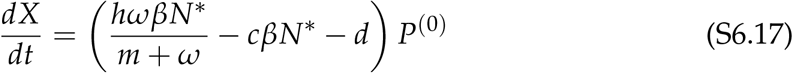

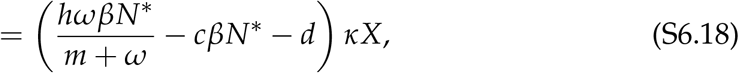

with *κ* > 0. Then the invasion growth rate is

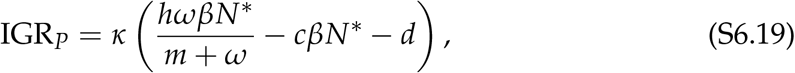

and the invasion criterion is identical to the condition in eq. S6.16:

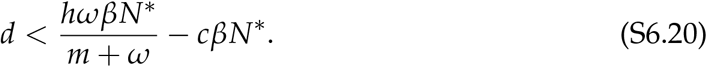

### S6.1.3 Method 2.2: Applying stability conditions

The dynamics of the parasite during invasion can be written in the linear matrix form:

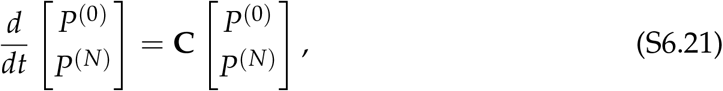

where the matrix **C** is given by

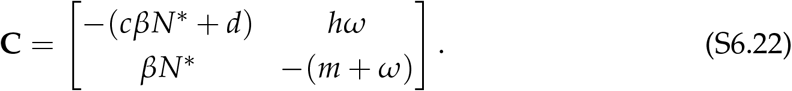

Considering the eigenvalues (*λ*) of **C** and applying the Routh–Hurwitz stability conditions, we find that the leading eigenvalue is positive if one of the following conditions holds. First, a negative determinant implies a positive dominant eigenvalue,

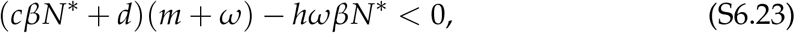

which matches the expression in eq. S6.16. Second, if this condition does not hold, the eigenvalues share a sign and a positive trace implies that both eigenvalues are positive:

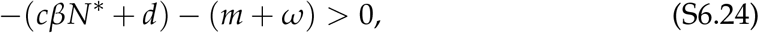

which cannot be fulfilled since all the parameters are strictly positive. Therefore, the determinant criterion solved above is the necessary and sufficient condition for the invasion of the parasite.

#### S6.1.4 Parameter effects on parasite invasion

The equilibrium abundances of the host–pest system (*S*\*, *N*\*) can easily be obtained by setting ^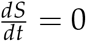^ and ^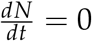^, which gives 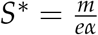 and 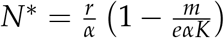. Substituting *N*\* into the inequality in eq. S6.16, the invasion criterion of the parasite can be obtained as:

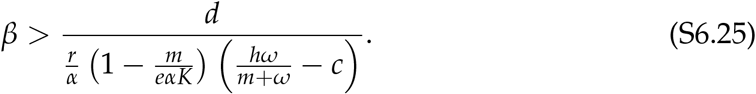

We find that positive invasion growth of the parasite requires that the infection rate of the parasite must be higher than a threshold determined by host, pest, and parasite traits. In other words, the higher the infection rate of the parasite, the more likely it is that the parasite can invade and persist when introduced to the host–pest system.

### S6.2 Invasion analysis for pest

Now, we turn our focus to the second stage-structured dynamics in the general model: pests can be either healthy 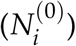 or infected 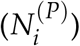 by parasites. Considering two pest species, and reverting back to the notation in the general model, the system is

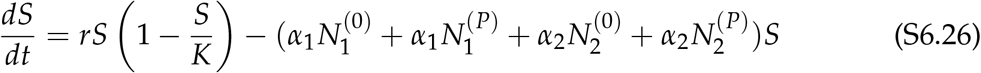

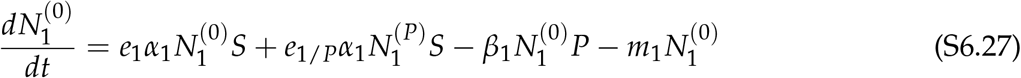

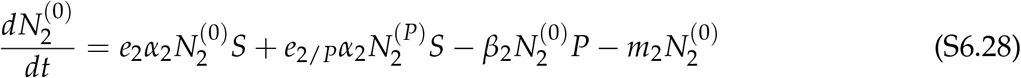

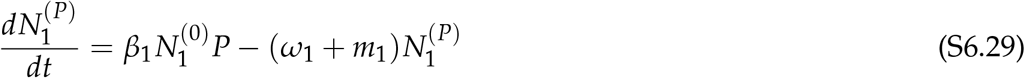

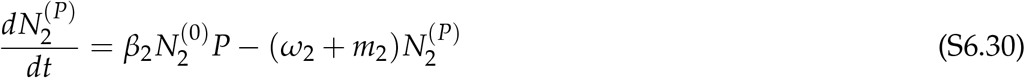

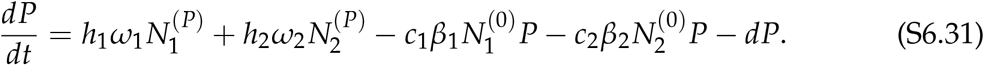

Below, we perform the invasion analysis for pest species 1, with stage-structured state variables 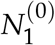 and 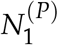, to invade the resident equilibrium of pest species 2 (*S*\*, 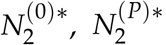, *P*\*).

#### S6.2.1 Method 1.1: Direct timescale separation

We will choose a new pair of variables: *N*_1_, representing the slow dynamics of population growth, and 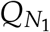, representing the fast dynamics of population stage structure. We choose 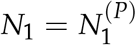for the population (slow) variable. Accordingly, the stage structure (fast) variable is the ratio of uninfected to parasitized pests, 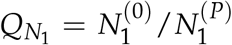. Then 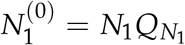 and we can rewrite the system considering the invasion scenario:

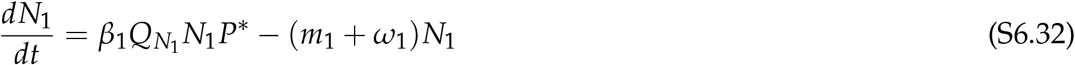

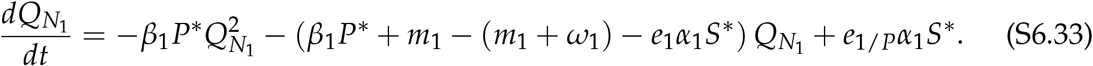

First, we solve for the quasi-equilibrium of 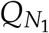 in the invasion condition. Solving using the quadratic formula gives the single biologically meaningful root,

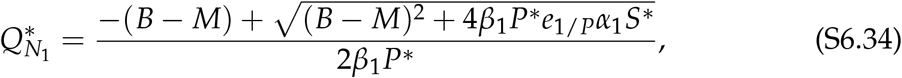

where *M* = *m*_1_ + *ω*_1_ and *B* = *β*_1_*P*\* + *m*_1_ *− e*_1_*α*_1_*S*\*. Next, we find the per-capita growth rate of *N*_1_ when the resident species are at equilibrium (*S*\*, 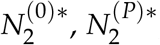, and *P*\*). We find that the invasion growth rate of *N*_1_ is expressed as

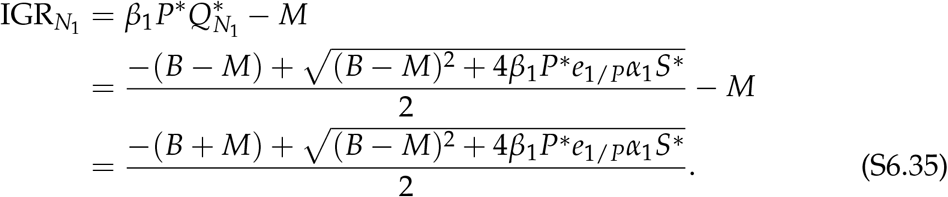

Finally, we solve for the conditions under which the invasion growth rate is positive:

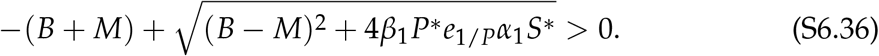

We now discuss the sign of 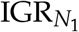 based on the sign of *B* + *M*. If *B* + *M* is negative (i.e., *e*_1_*α*_1_*S*\* *− β*_1_*P*\* *− m*_1_ > *m*_1_ + *ω*_1_), then 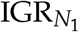 must be positive. Alternatively, if *B* + *M* is positive (i.e., *e*_1_*α*_1_*S*\* *− β*_1_*P*\* *− m*_1_ < *m*_1_ + *ω*_1_), we derive the following condition for positive invasion growth rate:

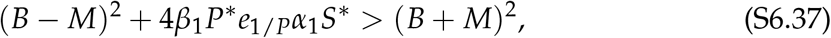

which can be simplified to the following:

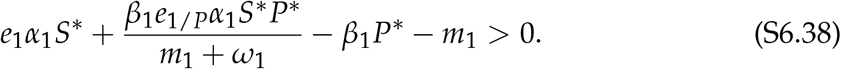

Given that the condition for *B* + *M* < 0 also implies the above condition, eq. S6.38 is the necessary and sufficient condition for pest species 1 to invade the monoculture of pest species 2 along with its associated host plant and parasite abundances.

#### S6.2.2 Method 1.2: Strategic linear combination

Let 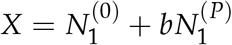, with 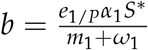. Then

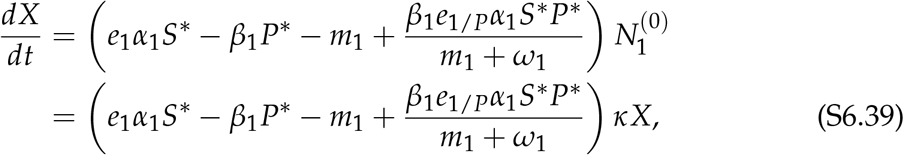

with *κ* > 0. Then the invasion growth rate is

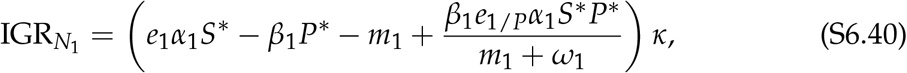

and the invasion criterion is the same as that in eq. S6.38:

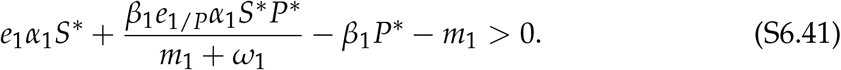

#### S6.2.3 Method 2.2: Applying stability conditions

The dynamics of pest species 1 during invasion can be written in the linear matrix form:

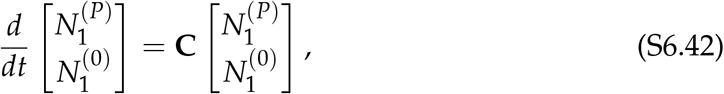

where the matrix **C** is given by

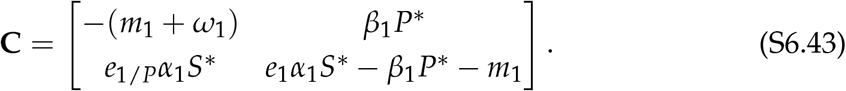

Considering the eigenvalues (*λ*) of **C** and applying the Routh–Hurwitz stability conditions, we find that the leading eigenvalue is positive if one of the following conditions holds. First, a negative determinant implies a dominant eigenvalue with nonnegative real part,

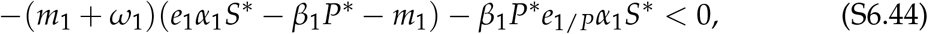

which matches the expression in eq. S6.38. Second, if this condition does not hold, the eigenvalues share a sign and a positive trace implies that both eigenvalues are positive:

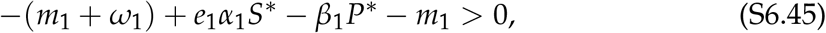

but as we noted above, this condition also implies the first; therefore, the determinant criterion solved above is the necessary and sufficient condition for the invasion of species 1.

#### S6.2.4 Parameter effects on pest invasion

From eq. S6.38, we find that for pest species 1 to invade, the parameters must satisfy

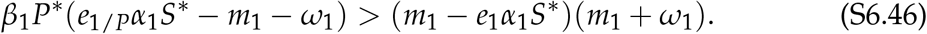

Let *M*_1_ = *m*_1_ + *ω*_1_, representing the mortality of infected individuals. If *e*_1/*P*_*α*_1_*S*\* > *M*_1_, we divide the above expression by the positive term with the parenthesis on the left-hand side, and rewrite the inequality as

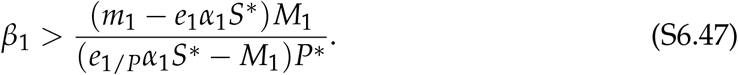

Since *e*_1/*P*_ *≤ e*_1_ (i.e., parasite infection reduces the conversion efficiency of pests),

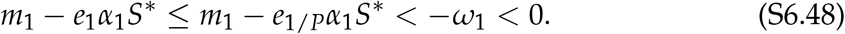

It follows that the right-hand side of equation S6.47 is less than zero, and the invasion criterion for *N*_1_ is always fulfilled since *β*_1_ > 0. Biologically speaking, when *e*_1/*P*_*α*_1_*S*\* > *M*_1_, the mortality of infected individuals is relatively low while their energy gain from host consumption is relatively high; therefore, *N*_1_ can always invade regardless of the pest’s susceptibility to parasites (*β*_1_).

On the other hand, if *e*_1/*P*_*α*_1_*S*\* < *M*_1_, i.e., the resource level is low and the price of infection is high, species 1 can only invade if the infection efficiency of parasites satisfies

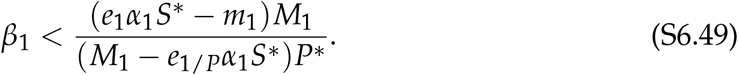

One can see that the above criterion is a decreasing function of *P*\*, meaning that a top-down control from higher resident parasite population makes invasion harder, and an increasing function of *S*\*, meaning that a bottom-up control from higher resident plant population provides more resource to make invasion easier. We can also rewrite the invasion criterion as

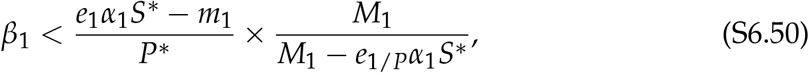

where the latter term increases when *M*_1_ decreases; thus, a lower mortality of infected individuals allows species 1 to endure a higher infection efficiency.

## S7 Stage-structured model with limit cycle

We consider the stage-structured resident species *N*_2_ growing alone in the cyclic model introduced in eq. 12 of the main text. Dropping the species subscripts (i.e., 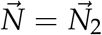, the subsystem is given by

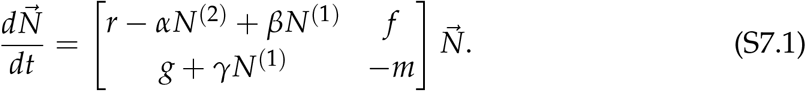

In this Appendix, we derive the conditions on facilitation *β, γ* for a limit cycle via a Hopf bifurcation.

### S7.1 Equilibrium

Let 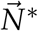 represent the resident equilibrium of the system and **C**\* represent the resident’s stage transition matrix (i.e., the matrix in eq. S7.1) evaluated at this resident equilibrium. Since 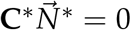 for a nonzero vector 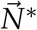, it must be the case that **C**\* is singular. Thus a necessary condition for equilibrium is det **C**\* = 0, which means the abundances lie on the line:

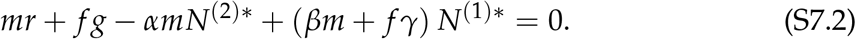

Next, setting *dN*^(2)^/*dt* = 0, we also have

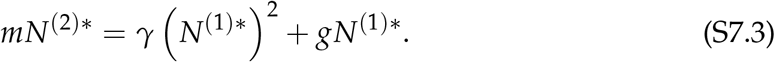

Substituting the expression for *N*^(2)*^ into the equilibrium condition gives

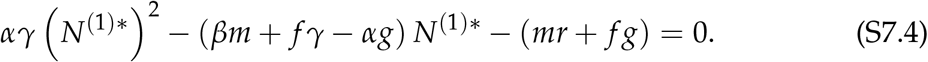

Then, we obtain

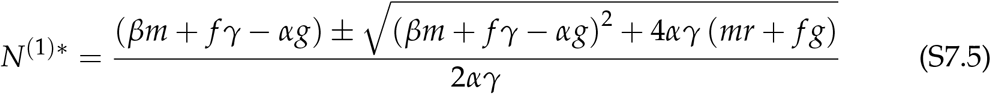

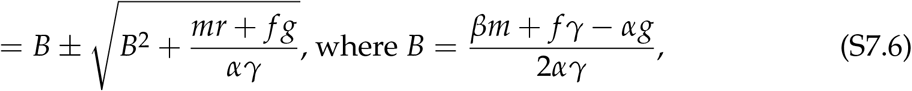

where we can ignore the *−* root because it is always negative and hence not biologically meaningful.

### S7.2 Hopf bifurcation

The first step to showing the emergence of a limit cycle is showing that the equilibrium loses stability through a Hopf bifurcation; i.e., as parameters are varied, the real parts of a complex conjugate pair of leading eigenvalues cross from negative to positive through zero.

Letting 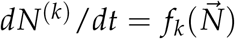, the Jacobian matrix is the matrix **J** such that the element in row *k* and column *l* is (**J**)_*kl*_ = *∂ f*_*k*_/*∂N*^(*l*)^. We have 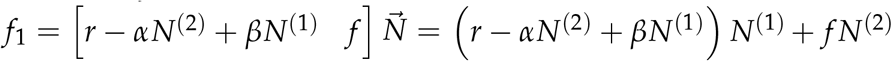 and 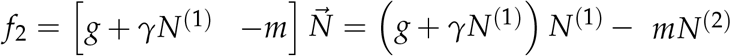 and thus, at equilibrium 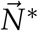 ,

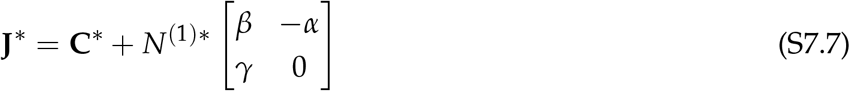

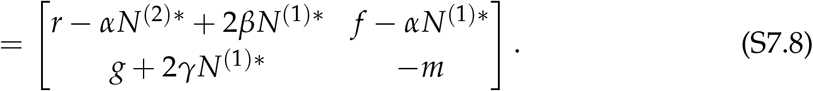

First, we derive the general criterion on **J**\* for a Hopf bifurcation in a two-dimensional system. Generally speaking, the characteristic equation is *λ*^2^ *−* (tr **J**\*) *λ* + det **J**\* = 0. A Hopf bifurcation occurs when the eigenvalues are a complex conjugate pair and the real part changes sign (i.e., they cross the imaginary axis while their imaginary parts remain nonzero). Thus, we have two conditions: (1) the real part changes sign at tr **J**\* = 0 for some parameter values, and (2) at those parameter values, the roots are complex, which occurs whenever det **J**\* > 0 (note that tr **J**\* = 0 here, so the sign of the polynomial’s discriminant is fully determined by the matrix’s determinant).

Now we consider our specific expression for the Jacobian. Leaving the elements of **C**\* unsubstituted at first,

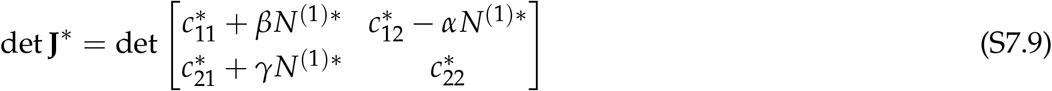

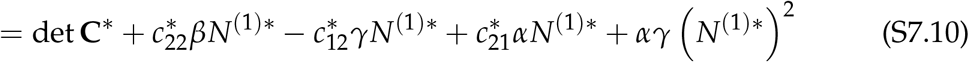

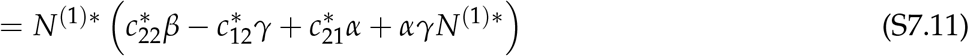

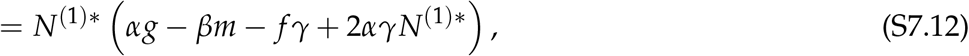

where we have used that det **C**\* = 0 at equilibrium and substituted in the expressions for elements of **C**\*. We note that *N*^(1)*^ is positive for a feasible equilibrium, so the other condition for the Hopf bifurcation is

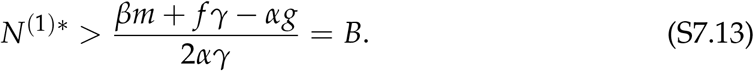

Based on our solution to the quadratic eq. S7.4, *N*^(1)*^ equals *B* plus some positive quantity, so this always holds. Next, note that the trace is

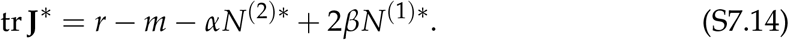

Then the trace becomes greater than zero, one condition for a Hopf bifurcation, when the inequality below goes from false to true:

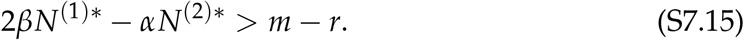

Using the condition for equilibrium, we can rewrite this as:

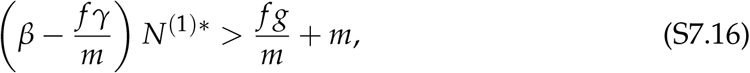

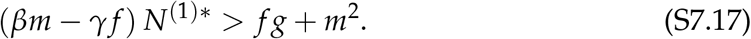

Feasibility requires *N*^(1)*^ > 0, and the right-hand side is always positive, so

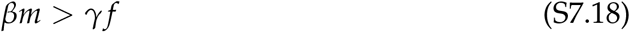

is a necessary (but not sufficient) condition for a Hopf bifurcation. Dividing by the positive quantity *βm − γf*, we find that a Hopf bifurcation occurs when the equilibrium abundance crosses above

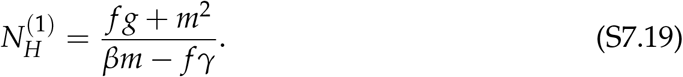

### S7.3 Boundedness

Let *N* = *N*^(1)^ and *Q* = *N*^(2)^/*N*^(1)^ as before. We will rewrite the system in terms of these transformed variables and, considering *Q* to be the fast variable, find its quasi-equilibrium *Q*\*. Rewriting the derivative, we find

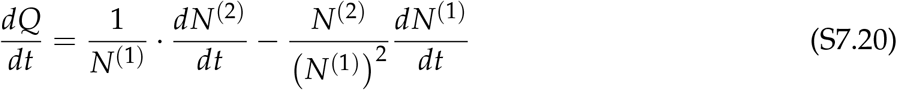

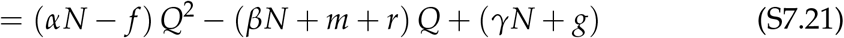

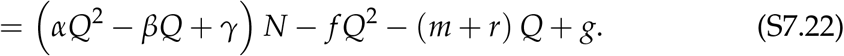

When *N →* ∞, the terms in *N* dominate *dQ*/*dt* and thus, the quasi-equilibrium *Q*\* obeys

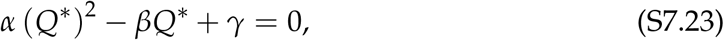

which has the solutions 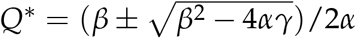. The discriminant *β*^2^ *−* 4*αγ* determines how many real roots there are. We use the discriminant to consider the effect of parameter values on the long-term behavior of *Q*; here, we ignore the boundary case where *β*^2^ *−* 4*αγ* = 0, where diagnosing long-term behavior requires a more rigorous analysis. If the discriminant is positive (*β*^2^ > 4*αγ*), both roots of *Q*\* are real and positive. Since *dQ*/*dt* is negative between the two roots, the smaller one is the stable solution; thus,

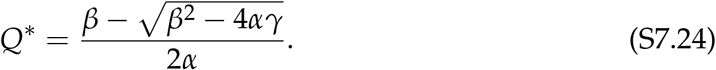

If the discriminant is negative (*β*^2^ < 4*αγ*), there are no real roots and *dQ*/*dt* is positive for all *Q*. This suggests that as long as *N* remains large enough, *Q* can grow arbitrarily large.

Now consider asymptotic population growth at high *N*. Writing *dN*/*dt* in terms of *Q* and *N*, we have

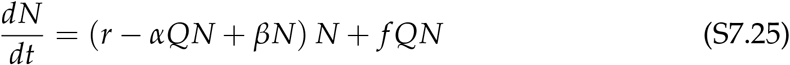

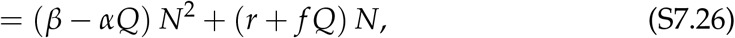

where the terms in *N*^2^ dominate as *N* becomes large. Thus, population growth at large *N* is determined by the coefficient *β − αQ*. Consider the two cases for the asymptotic behavior of *Q* derived above:

- If *β*^2^ > 4*αγ, Q* reaches a quasi-equilibrium *Q*\*. Substituting the expression for *Q*\* (eq. S7.24), we see that 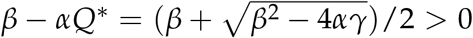. This means that *dN*/*dt* remains positive. Biologically we conclude that the resident settles on some fixed quasi-equilibrium population structure and grows indefinitely.
- If *β*^2^ < 4*αγ, Q* continues to grow: in terms of the original variables, adults *N*^(2)^ become increasingly abundant relative to juveniles *N*^(1)^. This eventually causes the coefficient *β − αQ* to become negative, thus making *dN*/*dt* negative as well. Thus, the system is driven back down to lower abundances. Biologically, in this case, as the resident population becomes large, it eventually becomes dominated by adults. Since adults hinder the population’s overall growth rate via their competitive effect on juveniles, this has the effect of preventing unbounded population growth.

Thus, our heuristic analysis suggests that

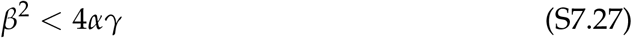

is required for population growth to remain bounded. Biologically, it ensures that the facilitative effect *β* of juveniles on their own replenishment rate is counterbalanced by the population regulation imposed by adults, as determined by the competition coefficient *α* and the *γ* parameter controlling facilitation of the juvenile-adult transition.

### S7.4 Parameter selection

Finally, we consider how to select parameters such that the system shows a stable limit cycle. The parameter *γ* is crucial: it regulates the population, preventing unbounded growth, but it must also be small enough that it doesn’t damp out oscillations. To see this, note that the necessary condition for the Hopf bifurcation is that

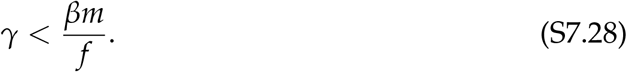

Boundedness, however, requires

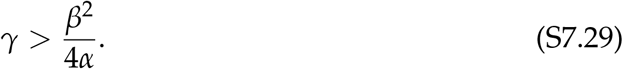

Thus, values of *γ* that allow stable limit cycles exist only when

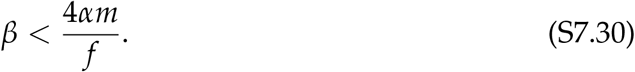

Having chosen a value of *β* satisfying this condition, we can then choose a *γ* value such that *β*^2^/4*α* < *γ* < *βm*/ *f*. Finally, we must ensure that the equilibrium juvenile abundance *N*^(1)*^ (eq. S7.6) is above the threshold 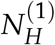 (eq. S7.19).

## S8 Stage-structured model with chaotic dynamics

Here, we analyze a multispecies extension of the model in equations 1a–1c of the main text. We consider a model of *n* interacting species with *n* + 1 state variables: the two stages of species 1, 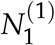 and 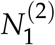, which we will analyze as the invading species, plus *n −* 1 non–stage-structured species *N*_2_, …, *N*_*n*_. These follow Lotka–Volterra dynamics,

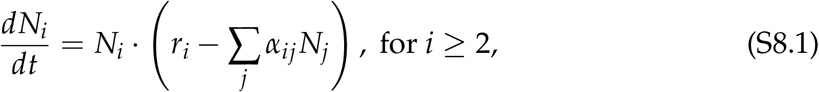

where, in the residents’ competition terms, we take *N*_1_ to represent species 1’s total abundance, 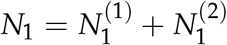. Here, *r*_*i*_ is the intrinsic growth rate of species *i*, and *α*_*ij*_ is the sensitivity of species *i*’s per-capita growth to competition from species *j*. Meanwhile, species 1’s dynamics obey

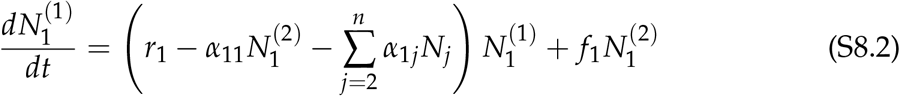

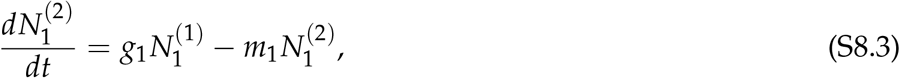

where *r*_1_ is the juvenile-to-juvenile transition rate (representing juvenile fecundity if positive, or mortality if negative), *f*_1_ is the fecundity of adults, *g*_1_ is the maturation rate of juveniles to adults, and *m*_1_ is the adult mortality rate. As in the stage-structured models in the main text (eqs. 1a–1c and eq. 12), competition affects the invader’s juvenile-to-juvenile transition rate (modifying the *r*_1_ term), and the invader’s competitive effect on itself (the *α*_11_ term) is mediated only by adult abundance, 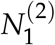. Note the different interpretations of *α*: the competitive effect caused by the invader (the *α*_*i*1_ terms) reflects both invader stages because it depends on the total abundance *N*_1_.

### Chaotic parameter values

We use 5 species (total 6 state variables). Species 1’s demographic parameters are as before: *r*_1_ = *−*1, *f*_1_ = 3, *g*_1_ = 1, *m*_1_ = 0.5. We couple this to a chaotic four-species Lotka–Volterra system represented by species 2 through 5, such that the complete set of *r*_*i*_ and *α*_*ij*_ coefficients is given by

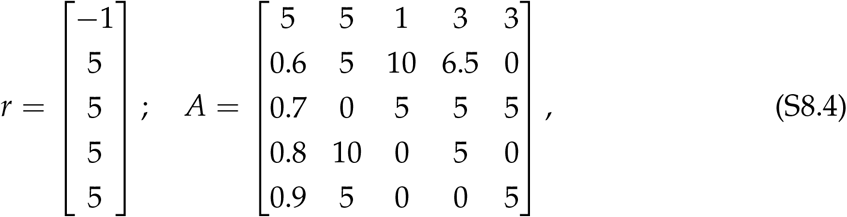

where (*A*)_*ij*_ = *α*_*ij*_. The Lotka–Volterra parameters for species 2 through 5 (i.e., *r*_2:5_ and *A*_2:5,2:5_) are based on the chaotic system discovered by Sprott (2010), scaled by a constant factor. As in the original system (see eq. 6.52 in Sprott, 2010), the non–stage-structured residents exhibit chaotic dynamics. Trajectories starting from near the unstable fixed point give rise to irregular fluctuations (Fig. S8.1a), and a phase plane shows an attractor with the complex interbraided structure typical of chaotic systems (Fig. S8.1b).

**Figure S8.1:**
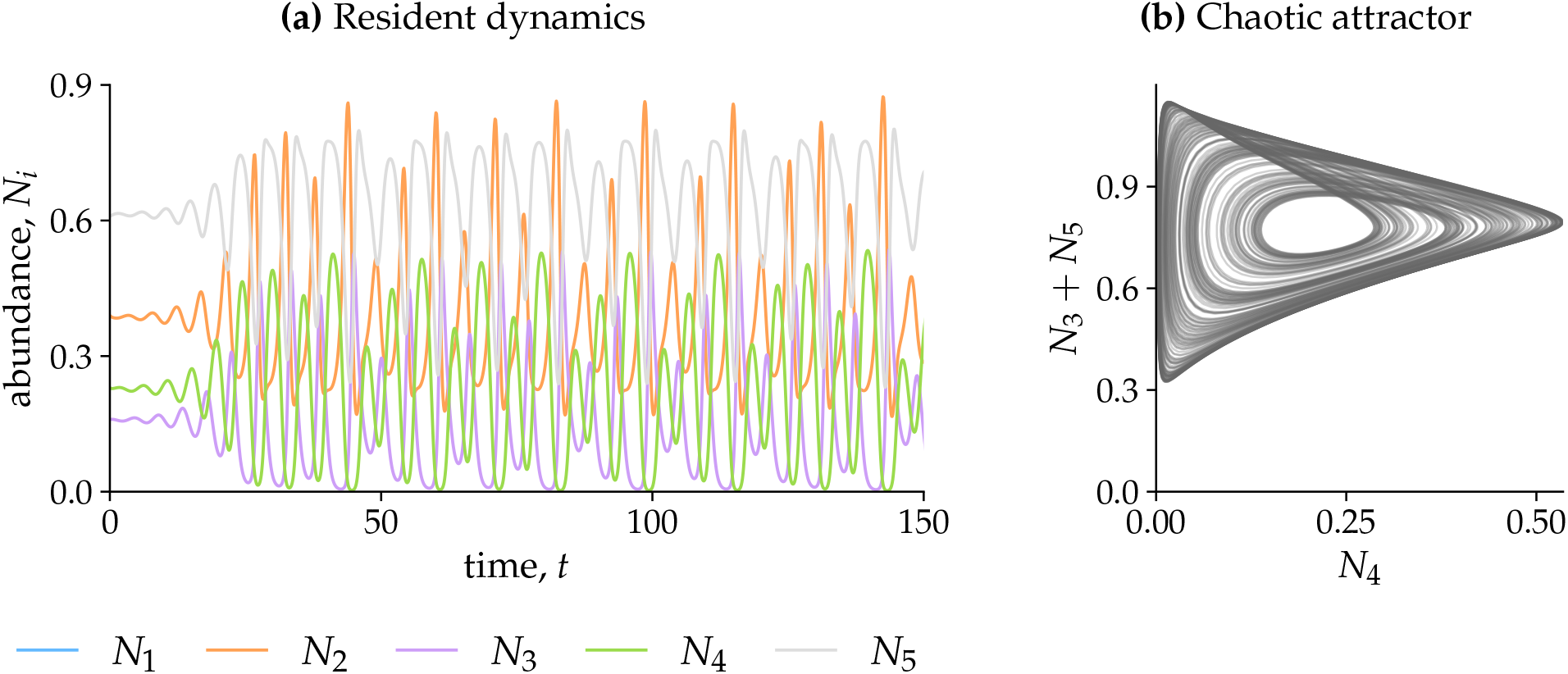
Chaotic resident dynamics for our parameter values. **(a)** Time series of species 2 through 5, starting from an initial condition near the unstable fixed point of the resident species’ dynamics. **(b)** Phase plane for resident species, based on *t* = 25 to 1000 of the same simulation.

### Invasion of stage-structured species

Just as in the simpler systems in the main text, invasion analysis can be used to study coexistence in this chaotic system. Introducing the stage-structured invader (species 1) to the system, we find that its total abundance increases successfully towards a persistent positive state (Fig. S8.2). Since its dynamics are driven by the residents’ chaotic fluctuations, the invader’s rate of growth also fluctuates irregularly. The total abundance *N*_1_ (thick blue line) fluctuates irregularly, but follows a broad trend of exponential growth before reaching a final state where it, along with all residents (colored background lines), can persist indefinitely with fluctuating abundance. In Fig. S8.3, we visualize the population structure of species 1 during this invasion trajectory. When the invader remains rare (i.e., until about *t* = 600), the proportion of its adults fluctuates chaotically around a constant average 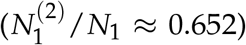. As the invader becomes more common, its population structure continues to fluctuate, but becomes more adult-dominated on average. Ultimately, the average proportion of adults reaches a plateau 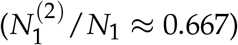 as the invader reaches its final fluctuating abundance in the community (after about *t* = 900).

**Figure S8.2:**
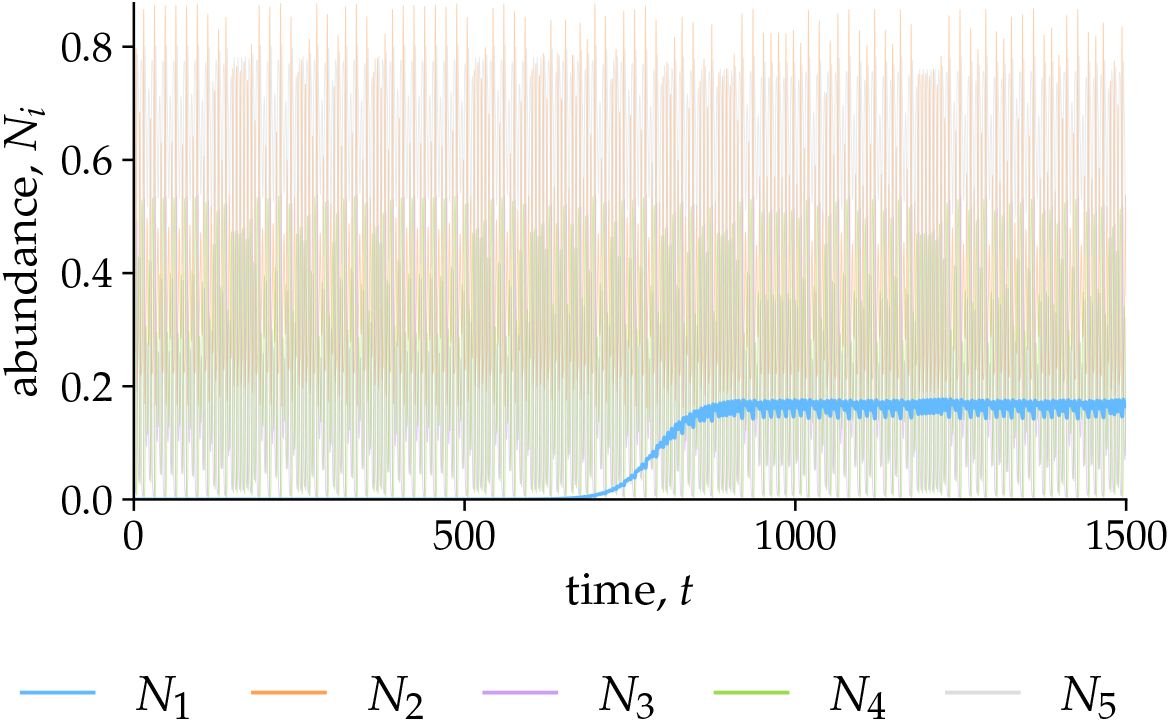
Invasion dynamics after introduction of stage-structured species 1. Starting from the final state of the resident simulation (Fig. S8.1), we introduce a small abundance of juveniles 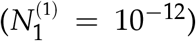 at the new time *t* = 0. The invader’s total abundance 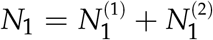 (thick blue line) increases over time, despite the irregular fluctuations in its growth rate driven by the residents’ chaotic dynamics (colored background lines).

**Figure S8.3:**
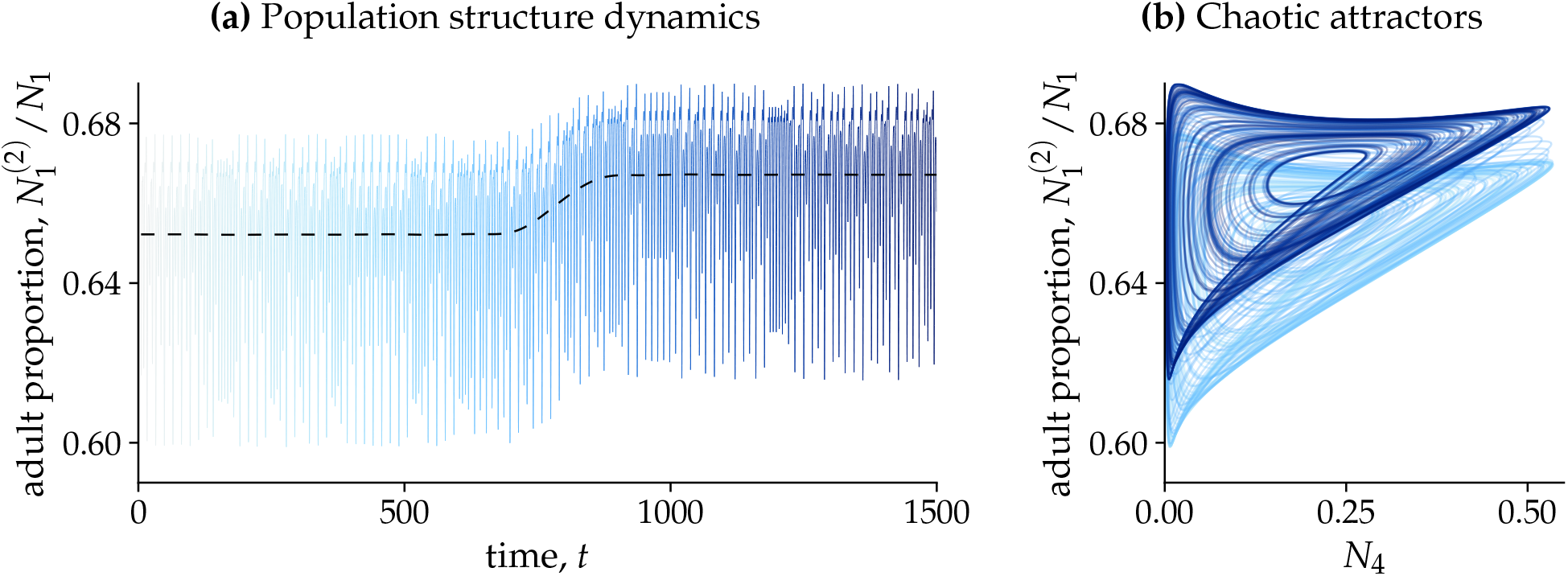
Population structure during invasion. As in Main Text Figure 2a, we consider the proportion of adults among the invader’s population, *N*^(2)^/*N*_1_. **(a)** Population structure dynamics are shown by plotting the adult proportion over time (thin blue line). To better display long-term dynamics, the first time unit is not visualized (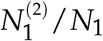 rapidly increasing from 0). For comparison to the next panel, the color varies from light blue (*t* = 0) to dark blue (*t* = 1500). Dotted line shows the smoothed average over a centered window of 50 time units. **(b)** Phase plane plot of adult proportion vs. the abundance of species 4, *N*_4_. Line color changes over time, matching (a), revealing two distinct attractors: one for resident-driven invasion dynamics (light blue) and one for the entire five-species system (dark blue).

**Figure S8.4:**
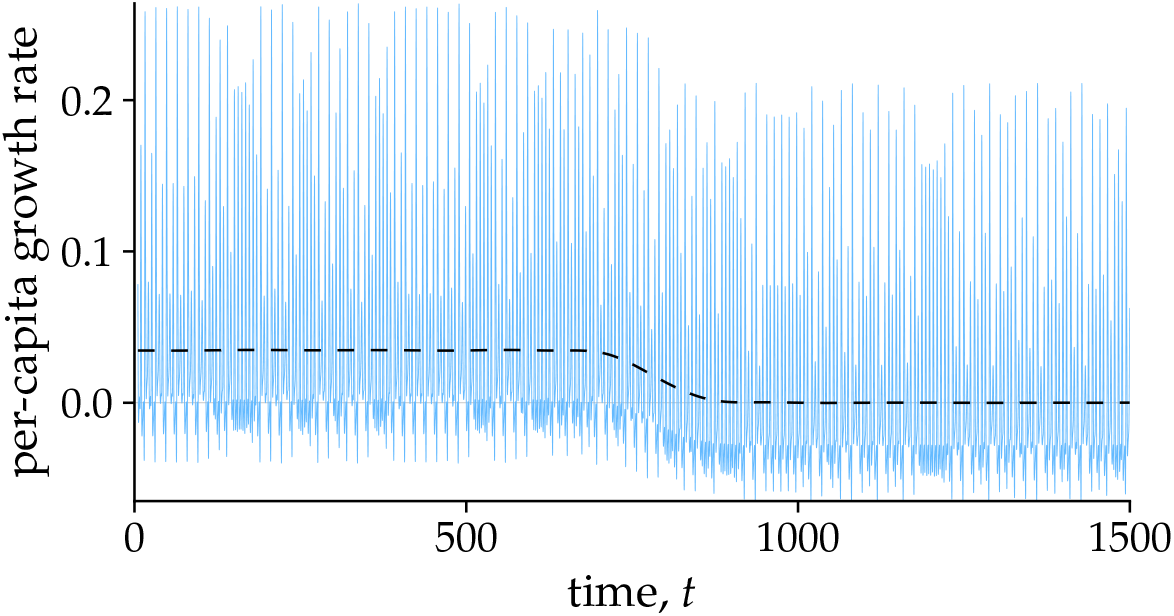
Per-capita growth rate of the stage-structured invader. (thin blue line) and its smoothed average over a centered window of 50 time units (black dashed line).

### Convergent invader dynamics

Despite these complex fluctuations, the insights of our framework still apply. While the invader remains rare, the dynamics of its population structure converge on a consistent attractor driven by the chaotic fluctuations of the residents. Indeed, a phase plane reveals that the invader’s population structure traces a chaotic attractor during invasion (Fig. S8.3b, light blue portion), before shifting to a different attractor once the invader reaches high abundance (Fig. S8.3b, dark blue). This means that if we keep the invader’s abundance low and take a long enough time period, its average properties—including its long-term growth rate—will converge to a consistent value. Indeed, evaluating the invader’s per-capita growth rate for our simulation (Fig. S8.4, thin blue line), we can confirm that the average growth rate (black dashed line) converges to a consistent value of IGR = 0.035 when the invader remains rare (i.e., until about *t* = 600).

### Calculating invasion growth rate using rescaling algorithm

As an alternative to simulating the full invasion trajectory, as in Figs. S8.2–S8.4, we can also apply the more efficient numerical algorithms in Appendix S3 to our chaotic system. As noted in the main text, the invasion growth rate of species 1 is equivalent to the Lyapunov exponent of the system around its resident attractor (species 2–5, Fig. S8.1). Accordingly, we can apply a variant of the algorithm for calculating Lyapunov exponents (Fig. S8.5a). Stated simply, we simulate the system as before, but repeatedly rescale the total abundance of the invader *N*_1_ in order to keep it rare. The growth rate GR_*i*_ between rescaling events can be calculated using the total abundance before rescaling (GR_*i*_ = log [*N*(*t*_*i*_)/*N*_0_] /Δ*t*). To calculate the invasion growth rate, we discard the initial burn-in period where the system has not yet reached its invasion attractor, then take the average of this growth rate over a large number of intervals (e.g., averaging over *t* = 5 to 1500 for the calculation in Fig. S8.5). Comparing the actual invasion dynamics to exponential growth at the rate predicted by our algorithm (IGR_1_ = 0.035) confirms that we have correctly found the invader’s long-term growth rate when it is rare.

### Calculating invasion growth rate using Floquet theory

Floquet theory rigorously holds only for periodic systems. However, by approximating our chaotic invasion system as a periodic system with a very long period (*T →* ∞), we can apply the Floquet theory calculation as a numerical algorithm for calculating the invasion growth rate in our chaotic system (Farrell & Ioannou, 1999; Sharma & Sinha, 2018). Note that when the invader is rare, the invader’s abundances 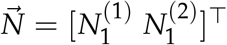 grow according to the time-varying linear system

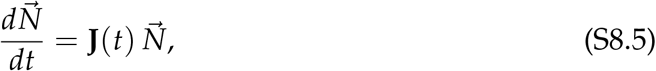

where **J** is the time-dependent Jacobian for species 1’s two stages,

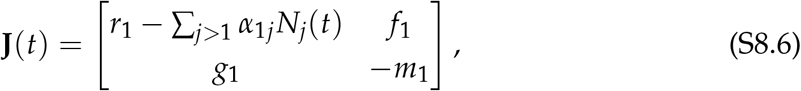

and the resident abundances *N*_*j*_(*t*) fluctuate chaotically over time. Applying the general method for calculating the monodromy matrix **M** (main text Box 3, eq. B3.5), we simulate the linear system in eq. S8.5 starting from the standard basis vectors 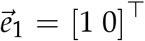 and 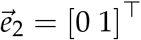 for *T* time units, where *T* is the large period used to approximate the chaotic system. For *T* = 1500 (matching the previous simulation), this gives

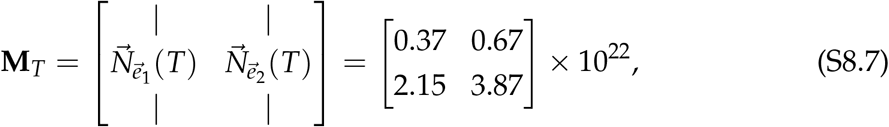

with the leading eigenvalue (i.e., Floquet multiplier) *ρ* = 4.24 *×* 10^22^. This corresponds to a Floquet exponent of (log *ρ*) /*T* = 0.035, matching the previous calculations. Comparing the results of this Floquet method to those from the previous algorithm for different simulation times (Fig. S8.6) highlights that the results converge to the same true invasion growth rate. Because the abundances do not need to be rescaled, this method also runs somewhat faster (∼ 2.5*×*) than the previous algorithm.

**Figure S8.5:**
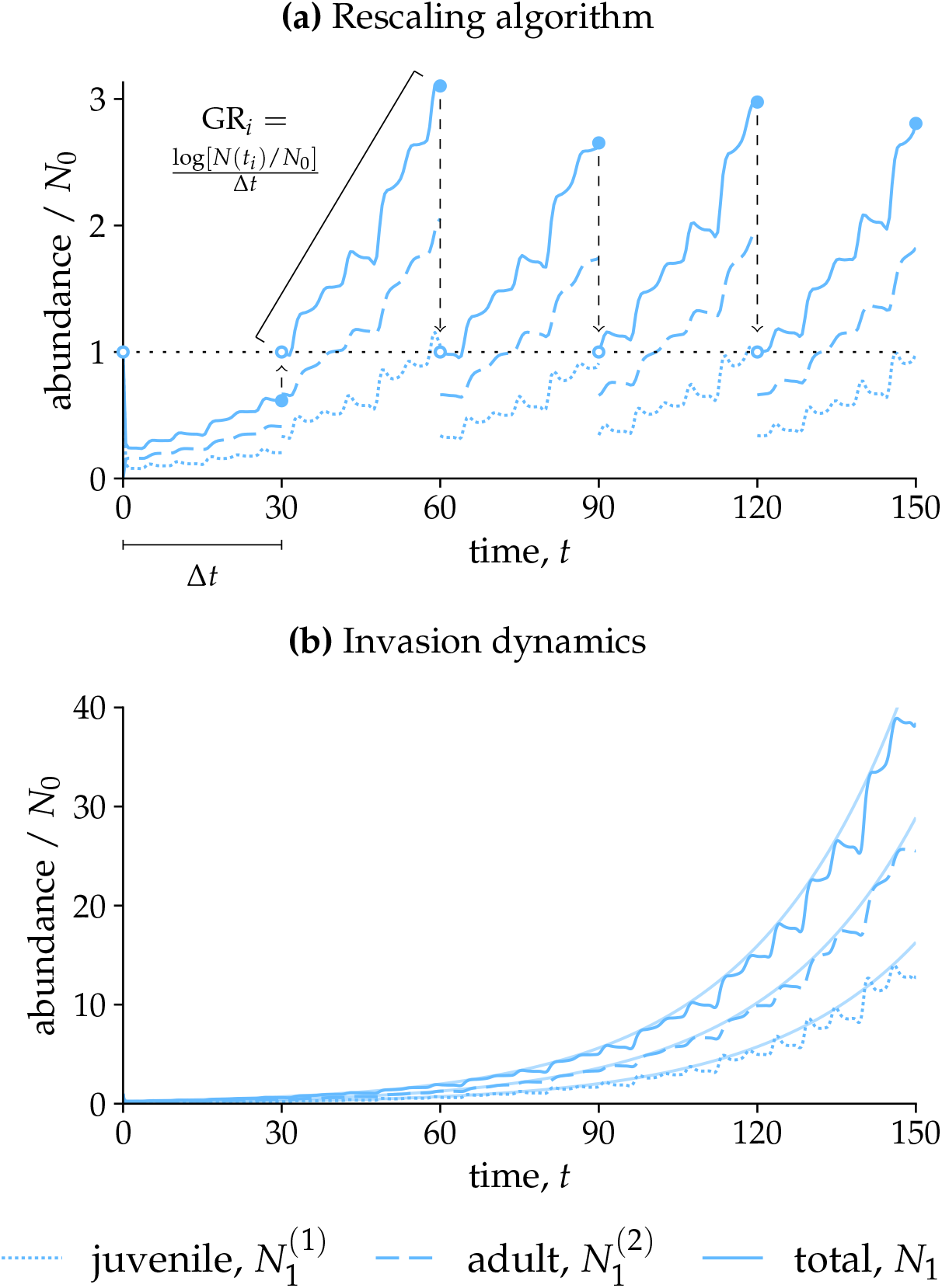
Finding the invasion growth rate in the chaotic model. **(a)** Applying the rescaling algorithm to simulate the invasion growth rate. The invader is introduced at a small target abundance *N*_0_ = 10^*−*12^ (dotted black horizontal line) and simulated for intervals of Δ*t* (here, for visualization purposes, 30 time units). After each interval *i*, the populations are rescaled proportionally, such that the total abundance (solid blue line) is scaled from *N*(*t*_*i*_) back to *N*_0_ (dashed arrows). The exponential growth rate GR_*i*_ is calculated for each interval, and the final IGR is obtained by averaging the interval growth rates over a large number of intervals after some burn-in period (Appendix S3). **(b)** Comparison of model dynamics (solid, dashed, and dotted blue lines) to exponential growth (faint blue curves) at the invasion growth rate IGR_1_ = 0.035 obtained using the rescaling algorithm. For greater accuracy, the invasion growth rate calculation uses a smaller interval (Δ*t* = 0.1) and a much larger time horizon (averaging from *t* = 5 to 1500) than depicted in panel (a).

**Figure S8.6:**
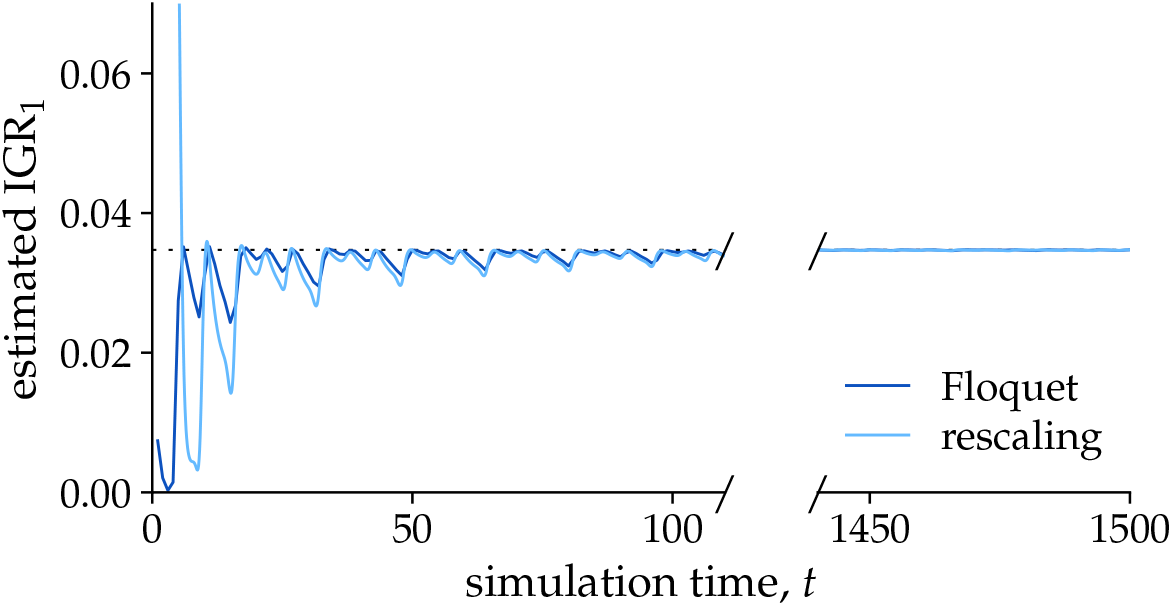
Comparison of methods for calculating invasion growth rate. The *x*-axis shows the length of simulation used to calculate invasion growth rate. The *y*-axis shows the estimated invasion growth rate: for the rescaling algorithm (light blue), this is the average growth rate between times 5 and *t*; for the Floquet method, this is (log *ρ*) /*t* for a monodromy matrix calculated for period *t*. Both methods converge to the same value of IGR_1_ (dotted line).

## S9 Floquet theory

Floquet theory considers solutions to the periodic system of differential equations

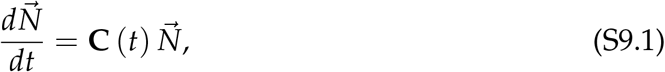

where **C** is a periodically varying matrix such that **C** (*t* + *T*) = **C** (*t*) for period *T*. Here, we provide a summary of key results in Floquet theory. Rather than a full set of formal proofs, we provide intuitive justifications for key derivations and refer readers to standard texts such as Chicone (1999) for a more rigorous treatment.

### S9.1 Principal fundamental matrix solution

For any system of the form in eq. S9.1, there exists a fundamental matrix solution **Φ** (*t*) such that **Φ** (0) = *I*, known as the *principal fundamental matrix solution*. To see this, let **Ψ** (*t*) be any fundamental matrix solution. Then,

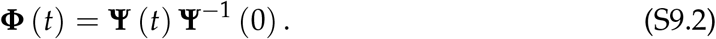

Indeed, this is a linear combination of columns of **Ψ** and is therefore also a solution. Moreover, when *t* = 0, the matrices cancel to give the identity matrix *I*, ensuring that **Φ** (0) = *I* as required.

Note that the principal fundamental matrix solution has the convenient property that any solution to the system can be written

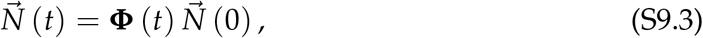

where 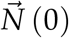 is the initial condition. This also gives a convenient way to numerically calculate **Φ**. Let 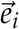 be the *i*-th standard basis vector; i.e., a vector with one at position *I* and zeros elsewhere. Then the product **Φ** (*t*) with 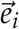 is simply its *i*-th column; on the other hand, it also represents the solution at time *t* of a system starting with 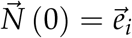. Thus, we can assemble **Φ** (*t*) from a series of columns representing different solutions to eq. S9.1 at time *t*:

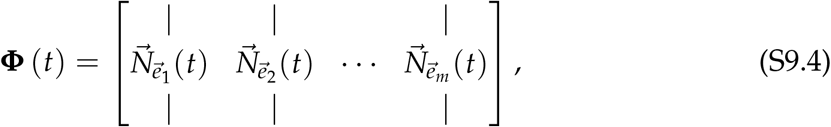

where the *i*-th column 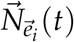 uses the *i*-th standard basis vector 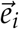 as its initial population vector.

### S9.2 The monodromy matrix M

Floquet proved that for a principal fundamental matrix solution **Φ** (*t*),

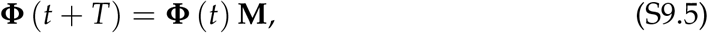

for all *t*, where **M** is the *monodromy matrix*, defined as

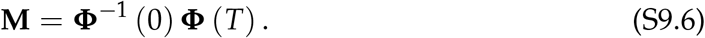

To see this, let **Ψ** (*t*) = **Φ** (*t* + *T*). This time-shifted function is also a fundamental matrix solution of the system due to the fact that **C** (*t* + *T*) = **C** (*t*), which we can verify by showing that its derivative satisfies the original system (eq. S9.1). Thus, there should exist some linear combination of columns of **Φ** that gives us **Ψ**.

We show that the monodromy matrix **M** defined above fulfills this condition. Consider the product of the original solution with **M** on the right: rewriting in terms of **Ψ**, this is **Φ** (*t*) **M** = **Φ** (*t*) **Φ**^*−*1^ (0) **Ψ** (0). Due to the fact that the system is linear, **Φ** (*t*) **M** (where each column is a linear combination of the columns of **Φ**) is also a solution of eq. S9.1. Moreover, its value matches that of **Ψ** at *t* = 0 due to cancellation of the inverse in

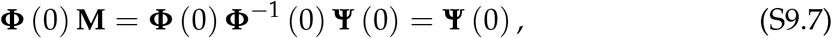

i.e., the solutions share an initial value. The solutions of a linear system are unique, meaning that **Φ** (*t*) **M** = **Ψ** (*t*) = **Φ** (*t* + *T*) for all *t*, as required.

Note that eq. S9.5 involves the product of the principal fundamental matrix solution with **M** on the right. We now derive the result in the main text, which concerns **M** on the left multiplying an individual vector solution 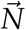. Note that the monodromy matrix is specific to a particular choice of fundamental matrix solution. Accordingly, let **Ψ** (*t*) be any fundamental matrix solution and let **M**′ represent its monodromy matrix. Then, 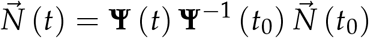 since the right-hand side equals 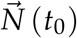 at *t* = *t*_0_. Then, we can write

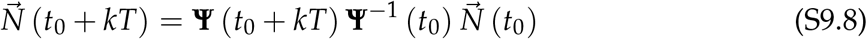

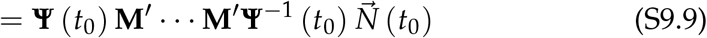

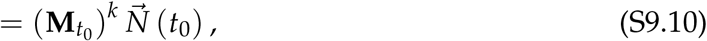

where

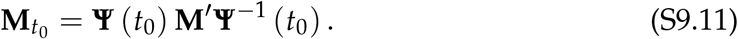

Unlike in eq. S9.5, the matrix that advances a single vector by one period depends on phase. Nonetheless, **M**_*t*0_ still corresponds to a specific monodromy matrix; in particular, it belongs to the specific fundamental matrix solution with **Ψ** (*t*_0_) = *I*. Thus, for the case *t*_0_ = 0 considered in the main text, **M** is precisely the monodromy matrix of the *principal* fundamental matrix solution, matching the suggested procedure for its column-wise calculation. More generally, though the actual value of **M** depends on the choice of fundamental matrix solution, its eigenvalues do not change, meaning the invasion and stability properties are well defined.

### S9.3 The Floquet exponent matrix B and the Floquet normal form

It is possible to show that, because **M** (the monodromy matrix for *t*_0_ = 0) is an invertible matrix, there exists some matrix **B** such that

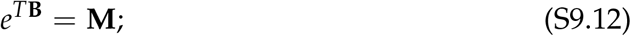

though the formal proof is beyond the scope of this appendix, interested readers may consult Chicone (1999, Chapter 2). Just as **M** captures the growth of the principal fundamental matrix solution over entire periods (i.e., in discrete time), **B** captures its overall growth in continuous time. However, we need another ingredient to capture the system’s variation within each period. To do so, we define a periodically varying matrix **P** (*t*) as

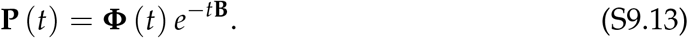

This function is also periodic with period *T* because **P** (*t* + *T*) = **Φ** (*t* + *T*) *e*^*−T***B**^*e*^*−t***B**^ = **Φ** (*t*) **MM**^*−*1^*e*^*−t***B**^ = **Φ** (*t*) *e*^*−t***B**^ = **P** (*t*). Furthermore, we can rearrange the equation to write

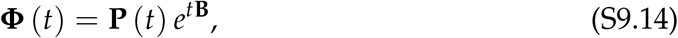

known as a *Floquet normal form* for the fundamental matrix **Φ** (*t*). Here, *e*^*t***B**^ is a matrix solution of the linear ordinary differential equation system *d***X**/*dt* = **BX** where **X** (0) = **I**. Accordingly, we can view **P** (*t*) as a periodically varying variable transformation mapping the underlying linear system described by **B** to the original periodic Floquet system described by eq. S9.1.

### S9.4 Eigenvalue analysis: the Floquet multipliers *ρ*_*i*_ **and Floquet exponents** *λ*_*i*_

Just as in non-periodic systems, eigenvalue analysis can tell us important information about the long-term fate of the system. Because **Φ** (*kT*) = **M**^*k*^ for any integer *k*, we can write the solution after a whole number *k* of periods using the monodromy matrix **M** and the initial condition 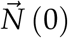 as

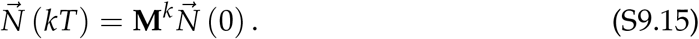

Letting the eigenvalues *ρ*_*i*_ and corresponding eigenvectors 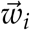 be solutions of 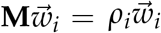, we can also write this whole-period solution as a sum of contributions from different eigenvectors:

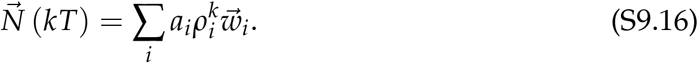

Thus, the eigenvalues and eigenvectors of **M** tell us the factor by which contributions to the solution can grow or shrink over one whole period; accordingly, the values of *ρ*_*i*_ are termed Floquet multipliers of the system.

Meanwhile, in continuous time, the eigenvalues and eigenvectors of **B** play an analogous role after we have accounted for the transformation **P** (*t*). Using eq. S9.14, we can write

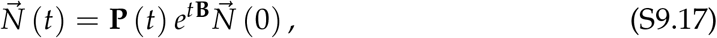

and analogously, we can write the 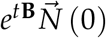 term as a sum of contributions from eigenvalues *λ*_*i*_ and eigenvectors 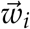 of **B**:

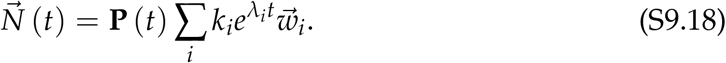

Since 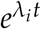 is a scalar, we can let 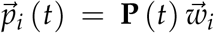 represent the periodically varying contribution associated with each eigenvector of the underlying linear system and rewrite the system as

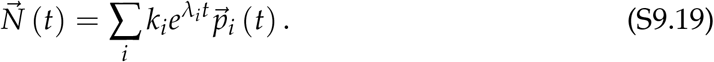

Here, the eigenvalues *λ*_*i*_ represent the continuous-time exponential growth rates of the system and are termed the Floquet exponents of the system.

Finally, note that it is possible to map between the discrete- and continuous-time representations of the system. Generally, the eigenvalues of a matrix **A** correspond to the eigenvalues of its exponential *e*^**A**^: if *λ*_*i*_ is an eigenvalue of **A**, 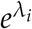 is an eigenvalue of *e*^**A**^ (Chicone, 1999, Chapter 2); furthermore, we can prove that the same eigenvector 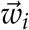 is associated with both values. The *i*th eigenvalue *λ*_*i*_ of **B** corresponds to an eigenvalue *Tλ*_*i*_ of *T***B** and, since **M** = *e*^*T***B**^, the *i*th eigenvalue *ρ*_*i*_ of **M** is equal to 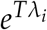. This allows us to relate the two sets of eigenvalues:

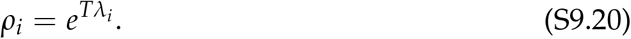

### S9.5 Long-term population dynamics

To apply this theory to population dynamics, we consider the special case when the original periodic system is positive: all solutions starting at positive abundances remain positive for all time. As discussed in Appendix S1, this is a natural requirement for a structured population model to be biologically realistic. Additionally, we will assume that after some integer number of periods *k*, a system starting from individuals only in one state *i* will eventually contain a nonzero quantity of state *j*.

Because multiplying by the monodromy matrix **M** advances the system by one period, **M** must also satisfy the conditions to maintain discrete-time positivity. As discussed for discrete-time matrices in Appendix S1, Box S1.B1, this means that all entries of **M** must be nonnegative. Meanwhile, our assumption that all states can be reached from all other states after some integer number of periods means that **M** must also be irreducible. Because **M** is nonnegative and irreducible, the Perron–Frobenius theorem (Appendix S1, Box S1.B2) guarantees that it has a unique, simple leading eigenvalue *ρ*_1_ and associated positive eigenvector 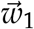 which is uniquely defined up to a constant.

We can also map these properties onto the continuous-time description of the system given by **B**. Because *e*^*T***B**^ = **M**, every eigenvector of **B** is also an eigenvector of **M** and their eigenvalues *ρ*_*i*_ and *λ*_*i*_ are related by 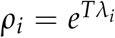. It is also possible to prove that **B** has a simple real leading eigenvalue

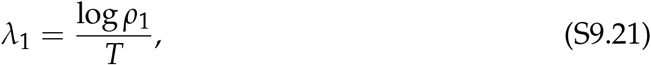

associated with the same unique positive eigenvector 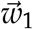 from above. This gives the key result of the main text, that population dynamics can be decomposed as

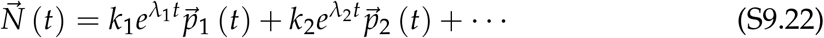

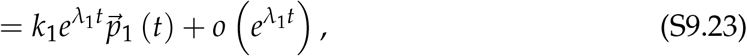

where *λ*_2_ has a strictly smaller real part than *λ*_1_. Here, the little-o notation 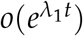 indicates that the transient terms grow more slowly than 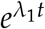.

## S10 Discrete-time perennial plant model

In this Appendix, we study competition between stage-structured perennial plant populations in discrete time. We use a model structure commonly applied to studies of plants experiencing discrete growth seasons. In particular, we consider a simplified version of the perennial model of Song *et al*. (2021) and Uricchio *et al*. (2019), without density-dependent seedling-to-adult recruitment. Closely related models have been applied in other studies modeling annual–perennial plant competition (e.g., Larios *et al*., 2017; Mordecai *et al*., 2015; Werner *et al*., 2025), and a model with the same structure was also analyzed in a general theoretical study by Fujiwara *et al*. (2011).

Each species *i* has three stages: seeds *N*_*i*_, juveniles *J*_*i*_, and adults *A*_*i*_. Each year, a proportion *γ*_*i*_ of seeds germinates and, of the remaining portion (the seed bank), a fraction *l*_*i*_ loses viability. A proportion *b*_*i*_ of germinants successfully becomes juveniles in the current year. A proportion *h*_*i*_ of juveniles successfully becomes adults in the next year, and the remainder is lost at the end of the year. Finally, a fraction *m*_*i*_ of adults dies each year, and all adults in a given year produce seeds (in the next year) according to some density-dependent fecundity *f*_*i*_, for instance the Beverton–Holt functional response (used in, e.g., Fujiwara *et al*., 2011; Song *et al*., 2021; Uricchio *et al*., 2019):

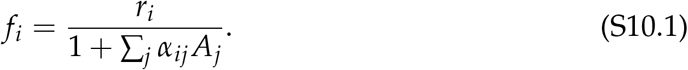

As a full analysis of the model is beyond the scope of this simplified case study, we write this functional response generally as *f*_*i*_ (*A*_1_, …, *A*_*n*_), where *n* is the number of species in the system.

Note that we census the seed population *N*_*i*_ at the beginning of a year, before any germination events, and juveniles *J*_*i*_ and adults *A*_*i*_ at the end of a year. With this in mind, the dynamics of seeds are given by

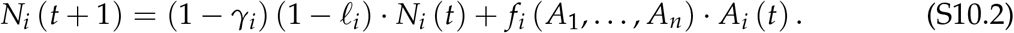

Juvenile abundances are proportional to the seed population in the same year,

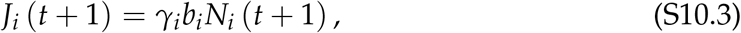

and adult abundances are given by

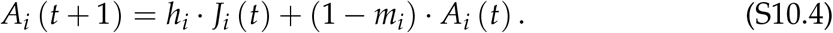

Note we do not need to track juveniles as a separate state variable, since their abundance can be written in terms of the seed population. Thus the entire system can be written as a recurrence relation giving seed and adult abundances in year *t* + 1 as a function of those in the current year *t*,

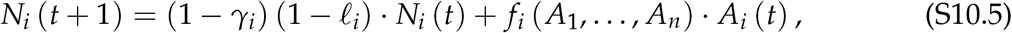

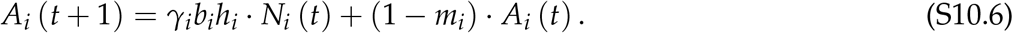

Define composite parameters: let *p*_*i*_ = 1 *−* (1 *− γ*_*i*_) (1 *− l*_*i*_) = *γ*_*i*_ + (1 *− γ*_*i*_) *l*_*i*_ represent the fraction of seeds lost each year (either due to successful germination or loss from the seed bank), and let *g*_*i*_ = *γ*_*i*_*b*_*i*_*h*_*i*_ represent the overall recruitment rate from seeds to adults in the next year (combining germination, seedling survival, and the juvenile–adult transition). Then

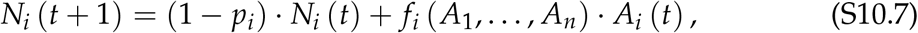

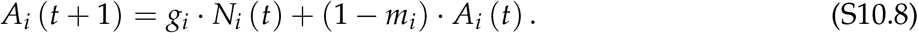

Now we derive the long-term growth rate of a species. Note *f*_*i*_ is the only demographic transition rate which is density-dependent; we will omit the function arguments for brevity. Defining 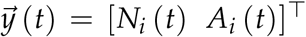 as a vector giving the state of the focal species *i*’s population at time *t*, we can write the dynamics in matrix form as

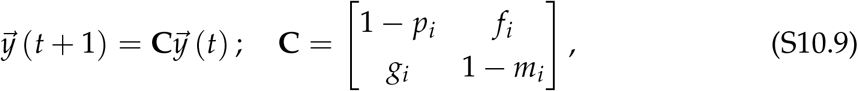

where **C** depends on adult densities through its upper-right *f*_*i*_ entry. Note that the matrix **C** describes discrete-time transitions (Appendix S1, Section S1.2); though this matrix was denoted **A** in Appendix S1, we use **C** here to avoid confusion with the adult abundance variable *A*_*i*_. Assuming the value of *f*_*i*_ is fixed at its value during invasion, the long-term growth factor per time step of species *i* is given by the leading eigenvalue of **C**, and we can derive the condition for invasibility in several ways.

### S10.1 Method 1.2: Strategic linear combination

Define the new variable

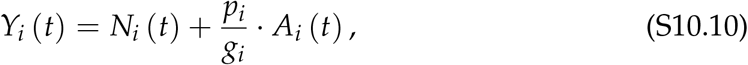

and assume that under long-term population growth, *A*_*i*_ (*t*) is some proportion of *Y*_*i*_ (*t*), i.e.,

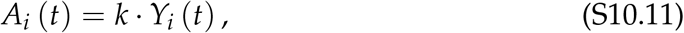

where *k* is positive. Then we can rewrite the recurrence relation in terms of only *Y*_*i*_ and

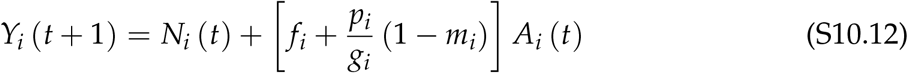

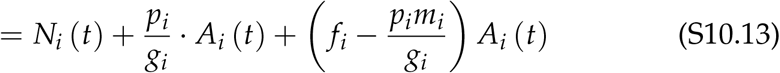

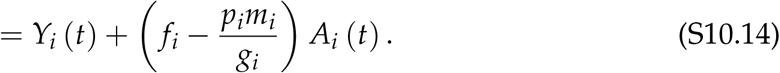

Furthermore, using eq. S10.11, we can write

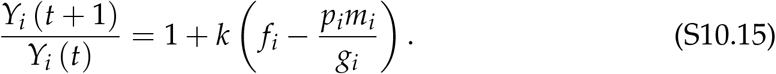

Since we are in a discrete-time system, positive population growth requires that this ratio is greater than unity. Since *k* > 0, we require that *f*_*i*_ *− p*_*i*_*m*_*i*_/*g*_*i*_ > 0, giving the simple persistence condition

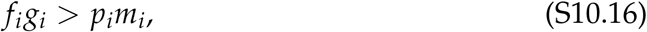

i.e., the positive effects of fecundity *f*_*i*_ and recruitment *g*_*i*_ on population growth must be great enough to account for the negative effects of seed loss *p*_*i*_ and adult mortality *m*_*i*_.

### S10.2 Method 2.1: Direct solution for eigenvalues

To find this specific value, we compute the characteristic polynomial of **C**:

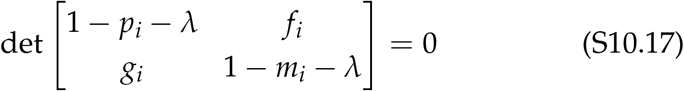

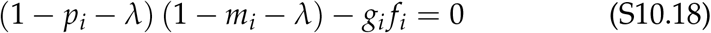

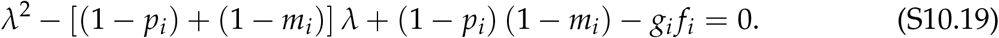

Then the eigenvalue with the greatest real part is given by

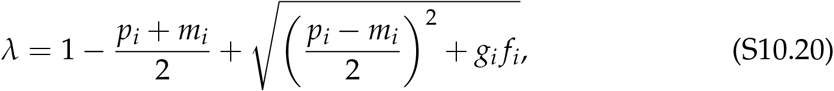

where the value under the square root is positive (due to *g*_*i*_ and *f*_*i*_ necessarily being positive). This matches the expression given in Fujiwara *et al*. (2011, see eq. 4 there).

Note that, as in the continuous-time case, *λ* determines the long-term growth rate of a species. However, its interpretation is different from that of the continuous-time invasion growth rate IGR: persistence requires *λ* > 1, not *λ* > 0 as in continuous time. One option is to define an analogous IGR in terms of *λ* such that IGR > 0 is the persistence condition: e.g., IGR = log *λ* (Spaak & De Laender, 2020) or IGR = *λ −* 1. Alternatively, we can directly solve for the conditions under which *λ* > 1. For this to occur, we require that

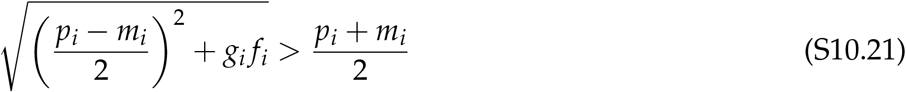

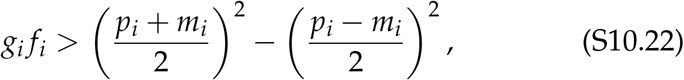

which we can simplify using the difference of squares to *g*_*i*_ *f*_*i*_ > *p*_*i*_*m*_*i*_, identical to the simple condition (eq. S10.16) derived above.

### S10.3 Method 2.2: Stability conditions

In the previous calculation, we looked for the conditions under which the maximum eigenvalue of **C** was greater than unity:

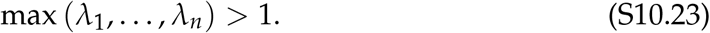

In the case where *n* = 2, this was feasible by directly solving for *λ* and rearranging the quadratic root in the solution. However, when *n* > 2, solving the characteristic polynomial may be difficult or impossible; moreover, eq. S10.23 does not match the conditions which the Routh–Hurwitz criteria diagnose (i.e., whether the leading eigenvalue is positive or negative). As described in Appendix S1, section S1.2, we can shift the matrix by the identity to obtain a problem where the Routh–Hurwitz criteria can be applied. In brief, subtracting the identity matrix,

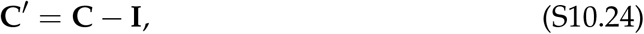

shifts each eigenvalue to 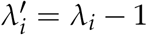. Thus, the discrete-time growth threshold *λ* > 1 becomes *λ*′ > 0, allowing us to apply the Routh–Hurwitz criteria to **C**′.

#### Two-dimensional example

Directly using the system above (eq. S10.9), we can derive **C**′ by subtracting one from the diagonal elements, giving

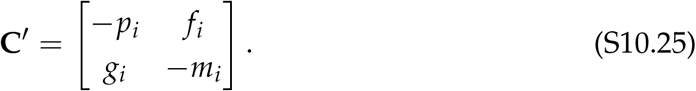

The Routh–Hurwitz conditions for a 2 *×* 2 matrix state that all eigenvalues have negative real parts if and only if det **C**′ > 0 and tr **C**′ < 0. For nonnegative population growth (one or more eigenvalues with nonnegative real parts), at least one of these conditions must not hold. Note that tr **C**′ = *−p*_*i*_ *− m*_*i*_ is always negative due to seed loss *p*_*i*_ and mortality *m*_*i*_ being non-zero proportions. Then nonnegative population growth requires that the determinant condition not hold; i.e., det **C**′ *≤* 0; moreover, we can infer that positive population growth requires det **C**′ < 0. Thus, the persistence condition is

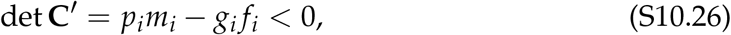

matching eq. S10.16.

#### Three-dimensional example

As an illustration of the advantage of using the Routh– Hurwitz conditions, say we had not simplified the system by removing the variable for juvenile population. In this case, the juvenile population (censused after germination) is determined by multiplying the seed abundance from the same year (censused before germination) by germination and juvenile recruitment *γ*_*i*_*b*_*i*_ (eq. S10.3). This gives the full three-stage system

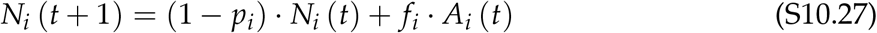

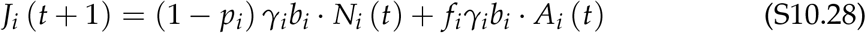

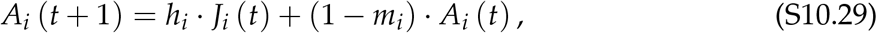

where *f*_*i*_ represents the density-dependent fecundity function *f*_*i*_ (*A*_1_, …, *A*_*n*_). Furthermore, we modify this system by assuming that juveniles can persist between years, such that a proportion 1 *− q*_*i*_ is retained each year into the next:

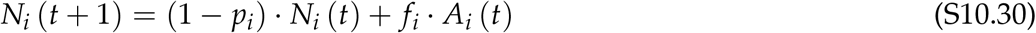

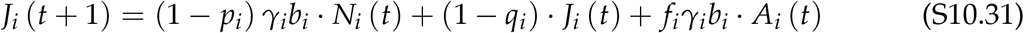

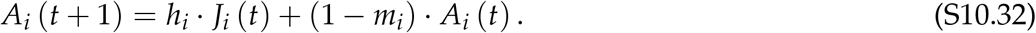

Letting 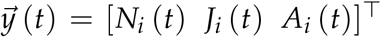 with an extra juvenile stage, the system is now 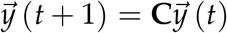 with

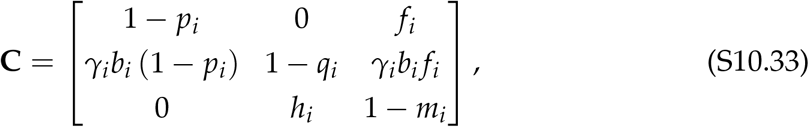

and the shifted matrix

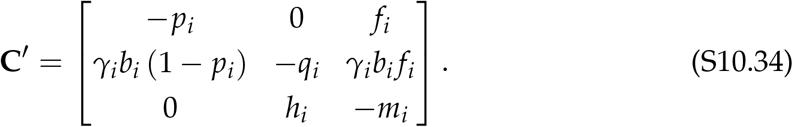

The characteristic polynomial is now

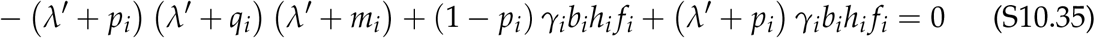

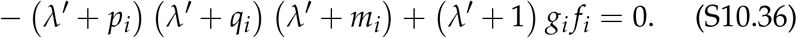

Expanding and taking the negative of both sides to obtain a positive leading coefficient of one,

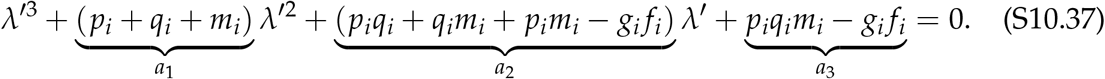

The Routh–Hurwitz conditions state that all eigenvalues have negative real parts if and only if *a*_1_ > 0, *a*_3_ > 0, and *a*_1_*a*_2_ > *a*_3_. Note that *a*_1_ > 0 holds automatically because all three parameters (loss from seed, juvenile, and adult stages) are positive. Then, *a*_3_ > 0 is necessary and sufficient for all eigenvalues to have negative real parts. To see this, rearrange the last condition to *a*_1_*a*_2_ *− a*_3_ > 0 and substitute:

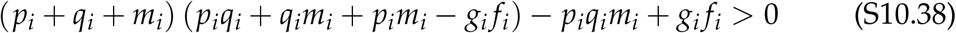

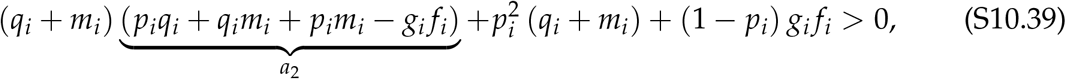

where the second term is necessarily positive and the last term is nonnegative (because *p*_*i*_ *≤* 1). Note that *a*_3_ < *a*_2_ because

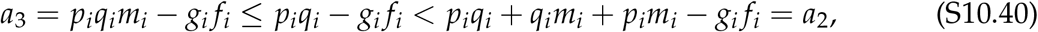

and thus, as long as *a*_3_ *≥* 0, the entire left-hand side is positive. Furthermore, note that in the boundary case *a*_3_ = 0, the leading eigenvalue is exactly zero.

Thus, persistence *λ*′ > 0 occurs if and only if *a*_3_ < 0, giving the persistence condition

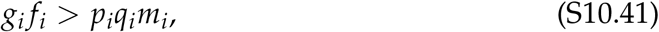

which has an interpretation paralleling the previous conclusion: recruitment *g*_*i*_ and fecundity *f*_*i*_ must outweigh losses in the three stages. When *q*_*i*_ = 1 (complete loss of juveniles), we recover the simpler two-stage model and its persistence condition (eq. S10.16). This simple criterion also generalizes to other specialized reproductive strategies, e.g., semelparous (juvenile persistence *q*_*i*_ < 1 and complete adult mortality *m*_*i*_ = 1) or biennial life history (no juvenile persistence *q*_*i*_ = 1 and complete adult mortality *m*_*i*_ = 1).

## References

Barabás, G., D’Andrea, R. & Stump, S.M. (2018) Chesson’s coexistence theory. Ecological Monographs, 88, 277–303. 10.1002/ecm.1302.

Barone, S.R., Narcowich, M.A. & Narcowich, F.J. (1977) Floquet theory and applications. Physical Review A, 15, 1109–1125. 10.1103/PhysRevA.15.1109.

Benaïm, M. & Schreiber, S.J. (2009) Persistence of structured populations in random environments. Theoretical Population Biology, 76, 19–34. 10.1016/j.tpb.2009.03.007.

Benettin, G., Galgani, L., Giorgilli, A. & Strelcyn, J.-M. (1980) Lyapunov Characteristic Exponents for smooth dynamical systems and for hamiltonian systems; A method for computing all of them. Part 2: Numerical application. Meccanica, 15, 21–30. 10.1007/BF02128237.

Brennan, J. & Schreiber, S.J. (2026) Using Modern Coexistence Theory to Understand Community Disassembly. Ecology Letters, 29, e70424. 10.1111/ele.70424.

Bukkuri, A. & Brown, J.S. (2023) Integrating eco-evolutionary dynamics into matrix population models for structured populations: Discrete and continuous frameworks. Methods in Ecology and Evolution, 14, 1475–1488. 10.1111/2041-210X.14111.

Caswell, H. (2001) Matrix Population Models: Construction, Analysis, and Interpretation. Sinauer Associates, Sunderland, Mass, 2nd ed.

Chesson, P. (2000) Mechanisms of maintenance of species diversity. Annual review of Ecology and Systematics, 31, 343–366.

Chesson, P. & Huntly, N. (1997) The Roles of Harsh and Fluctuating Conditions in the Dynamics of Ecological Communities. The American Naturalist, 150, 519–553. 10.1086/286080.

De Roos, A.M. & Persson, L. (2013) Population and Community Ecology of Ontogenetic Development. Princeton University Press, 1st ed.

De Roos, A.M., Schellekens, T., Van Kooten, T., Van De Wolfshaar, K., Claessen, D. & Persson, L. (2008) Simplifying a physiologically structured population model to a stage-structured biomass model. Theoretical Population Biology, 73, 47–62. 10.1016/j.tpb.2007.09.004.

DeMalach, N., Shnerb, N. & Fukami, T. (2021) Alternative States in Plant Communities Driven by a Life-History Trade-Off and Demographic Stochasticity. The American Naturalist, 198, E27–E36. 10.1086/714418.

Diekmann, O., Heesterbeek, J.A.P. & Metz, J.A.J. (1990) On the definition and the computation of the basic reproduction ratio r 0 in models for infectious diseases in heterogeneous populations. Journal of mathematical biology, 28, 365–382.

Diekmann, O., Heesterbeek, J.A.P. & Roberts, M.G. (2009) The construction of next-generation matrices for compartmental epidemic models. Journal of the royal society interface, 7, 873.

Ellner, S.P., Snyder, R.E. & Adler, P.B. (2016) How to quantify the temporal storage effect using simulations instead of math. Ecology Letters, 19, 1333–1342. 10.1111/ele.12672.

Ellner, S.P., Snyder, R.E., Adler, P.B. & Hooker, G. (2019) An expanded modern coexistence theory for empirical applications. Ecology Letters, 22, 3–18. 10.1111/ele.13159.

Ellner, S.P., Snyder, R.E., Adler, P.B., Hooker, G. & Schreiber, S.J. (2020) Technical Comment on Pande et al. (2020): Why invasion analysis is important for understanding coexistence. Ecology Letters, 23, 1721–1724. 10.1111/ele.13580.

Floquet, G. (1883) Sur les équations différentielles linéaires à coefficients périodiques. Annales scientifiques de l’École Normale Supérieure, 12, 47–88. 10.24033/asens.220.

Grainger, T.N., Levine, J.M. & Gilbert, B. (2019) The invasion criterion: a common currency for ecological research. Trends in ecology & evolution, 34, 925–935.

Gurney, W. & Nisbet, R. (1985) Fluctuation periodicity, generation separation, and the expression of larval competition. Theoretical Population Biology, 28, 150–180. 10.1016/0040-5809(85)90026-7.

Hofbauer, J. & Schreiber, S.J. (2010) Robust permanence for interacting structured populations. Journal of Differential Equations, 248, 1955–1971. 10.1016/j.jde.2009.11.010.

Iles, D.T., Salguero-Gómez, R., Adler, P.B. & Koons, D.N. (2016) Linking transient dynamics and life history to biological invasion success. Journal of Ecology, 104, 399–408. 10.1111/1365-2745.12516.

Iritani, R. & Day, T. (2026) Evolutionary invasion analysis for structured populations: a synthesis. 10.64898/2026.03.23.713828.

Ke, P.-J. & Levine, J.M. (2021) The temporal dimension of plant-soil microbe interactions: mechanisms promoting feedback between generations. The American Naturalist, 198, E80–E94. 10.1086/715577.

Klausmeier, C.A. (2008) Floquet theory: a useful tool for understanding nonequilibrium dynamics. Theoretical Ecology, 1, 153–161. 10.1007/s12080-008-0016-2.

Kussell, E. & Leibler, S. (2005) Phenotypic diversity, population growth, and information in fluctuating environments. Science, 309, 2075–2078.

Lefkovitch, L.P. (1965) The Study of Population Growth in Organisms Grouped by Stages. Biometrics, 21, 1. 10.2307/2528348.

Leslie, P.H. (1945) On the use of matrices in certain population mathematics. Biometrika, 33, 183–212. 10.1093/biomet/33.3.183.

Letten, A.D., Dhami, M.K.Ke, P.-J. & Fukami, T. (2018) Species coexistence through simultaneous fluctuation-dependent mechanisms. Proceedings of the National Academy of Sciences, 115, 6745–6750. 10.1073/pnas.1801846115.

Letten, A.D., Yamamichi, M., Richardson, J.A. & Ke, P.-J. (2024) Microbial Dormancy Supports Multi-Species Coexistence Under Resource Fluctuations. Ecology Letters, 27, e14507. 10.1111/ele.14507.

Lyapunov, A.M. (1907) Problème général de la stabilité du mouvement. Annales de la Faculté des sciences de l’Université de Toulouse pour les sciences mathématiques et les sciences physiques, 9, 203–474.

MacDougall, A.S., Gilbert, B. & Levine, J.M. (2009) Plant invasions and the niche. Journal of Ecology, 97, 609–615. 10.1111/j.1365-2745.2009.01514.x.

Nakazawa, T. (2015) Ontogenetic niche shifts matter in community ecology: a review and future perspectives. Population Ecology, 57, 347–354. 10.1007/s10144-014-0448-z.

Nakazawa, T. & Huang, C.-G. (2016) Two-species metacommunity dynamics mediated by habitat preference. Oikos, 125, 1334–1341. 10.1111/oik.03101.

Otto, S.P. & Day, T. (2011) A Biologist’s Guide to Mathematical Modeling in Ecology and Evolution. Princeton University Press.

Pande, J., Fung, T., Chisholm, R. & Shnerb, N.M. (2020) Mean growth rate when rare is not a reliable metric for persistence of species. Ecology Letters, 23, 274–282. 10.1111/ele.13430.

Polis, G.A., Myers, C.A. & Holt, R.D. (1989) The Ecology and Evolution of Intraguild Predation: Potential Competitors That Eat Each Other. Annual Review of Ecology and Systematics, 20, 297–330. 10.1146/annurev.es.20.110189.001501.

Price, P.W., Westoby, M., Rice, B., Atsatt, P.R., Fritz, R.S., Thompson, J.N. & Mobley, K. (1986) PARASITE MEDIATION IN ECOLOGICAL INTERACTIONS. Annual Review of Ecology and Systematics, 17, 487–505. 10.1146/annurev.es.17.110186.002415.

Roth, G. & Schreiber, S.J. (2014) Persistence in fluctuating environments for interacting structured populations. Journal of Mathematical Biology, 69, 1267–1317. 10.1007/s00285-013-0739-6.

Schreiber, S.J., Yamamichi, M. & Strauss, S.Y. (2019) When rarity has costs: coexistence under positive frequency-dependence and environmental stochasticity. Ecology, 100, e02664.

Shoemaker, L.G., Barner, A.K., Bittleston, L.S. & Teufel, A.I. (2020) Quantifying the relative importance of variation in predation and the environment for species coexistence. Ecology Letters, 23, 939–950. 10.1111/ele.13482.

Smith, G.R. & Wan, J. (2019) Resource-ratio theory predicts mycorrhizal control of litter decomposition. New Phytologist, 223, 1595–1606. 10.1111/nph.15884.

Song, C. & Spaak, J.W. (2024) Trophic tug-of-war: coexistence mechanisms within and across trophic levels. Ecology Letters, 27, e14409.

Song, C., Uricchio, L.H., Mordecai, E.A. & Saavedra, S. (2021) Understanding the emergence of contingent and deterministic exclusion in multispecies communities. Ecology Letters, 24, 2155–2168. 10.1111/ele.13846.

Spaak, J.W., Ke, P.-J., Letten, A.D. & De Laender, F. (2023) Different measures of niche and fitness differences tell different tales. Oikos, 2023, e09573.

Spaak, J.W. & Schreiber, S.J. (2023) Building modern coexistence theory from the ground up: the role of community assembly. Ecology Letters, 26, 1840–1861.

Spaak, J.W. & De Laender, F. (2020) Intuitive and broadly applicable definitions of niche and fitness differences. Ecology Letters, 23, 1117–1128. 10.1111/ele.13511.

Sprott, J.C. (2010) Elegant Chaos: Algebraically Simple Chaotic Flows. World Scientific, Singapore.

Usinowicz, J. & Levine, J.M. (2018) Species persistence under climate change: a geo-graphical scale coexistence problem. Ecology Letters, 21, 1589–1603. 10.1111/ele.13108.

Walker, E.J. & Gilbert, B. (2025) Allee effects and coexistence. The American Naturalist, 205, 559–571.

Wan, J., Ke, P.-J., Hordijk, I., Bialic-Murphy, L. & Crowther, T.W. (2025) Functional coexistence theory: Identifying mechanisms linking biodiversity and ecosystem function. Ecological Monographs, 95, e70033. 10.1002/ecm.70033.

Werner, C.M., Hallett, L.M. & Shoemaker, L.G. (2025) Fluctuation-Dependent Coexistence of Stage-Structured Species. The American Naturalist, 205, 327–341. 10.1086/733382.

Xu, L., Sun, X. & Zakem, E.J. (2026) Dependency–competition tradeoffs structure microbial niches and nitrogen cycling. ISME Communications, 6, ycag134. 10.1093/ismeco/ycag134.

Yamamichi, M., Gibbs, T.L. & Levine, J.M. (2022) Integrating eco-evolutionary dynamics and modern coexistence theory. Ecology Letters, 25, 2091–2106. 10.1111/ele.14078.

## References

Chicone, C. (1999) Ordinary Differential Equations with Applications. Number 34 in Texts in Applied Mathematics. Springer New York, New York, NY.

Farrell, B.F. & Ioannou, P.J. (1999) Perturbation Growth and Structure in Time-Dependent Flows. Journal of the Atmospheric Sciences, 56, 3622–3639. 10.1175/1520-0469(1999)056<3622:PGASIT>2.0.CO;2.

Frobenius, F.G. (1912) Über Matrizen aus nicht negativen Elementen. Sitzungsberichte der Königlich Preussischen Akademie der Wissenschaften zu Berlin, 1912, 456–477.

Fujiwara, M., Pfeiffer, G., Boggess, M., Day, S. & Walton, J. (2011) Coexistence of competing stage-structured populations. Scientific Reports, 1, 107. 10.1038/srep00107.

Hurwitz, A. (1895) Ueber die Bedingungen, unter welchen eine Gleichung nur Wurzeln mit negativen reellen Theilen besitzt. Mathematische Annalen, 46, 273–284. 10.1007/BF01446812.

Larios, L., Hallett, L.M. & Suding, K.N. (2017) Where and how to restore in a changing world: a demographic-based assessment of resilience. Journal of Applied Ecology, 54, 1040–1050. 10.1111/1365-2664.12946.

Liénard, A. & Chipart, M.H. (1914) Sur le signe de la partie réelle des racines d’une équation algébrique. Journal de Mathématiques Pures et Appliquées, 10, 291–346.

Metzler, L.A. (1945) Stability of Multiple Markets: The Hicks Conditions. Econometrica, 13, 277–292. 10.2307/1906922.

Mordecai, E.A., Molinari, N.A., Stahlheber, K.A., Gross, K. & D’Antonio, C. (2015) Controls over native perennial grass exclusion and persistence in California grasslands invaded by annuals. Ecology, 96, 2643–2652. 10.1890/14-2023.1.

Perron, O. (1907) Zur Theorie der Matrices. Mathematische Annalen, 64, 248–263. 10.1007/BF01449896.

Routh, E.J. (1877) A Treatise on the Stability of a Given State of Motion, Particularly Steady Motion. Macmillan and Co., London.

Sharma, A. & Sinha, S.C. (2018) An Approximate Analysis of Quasi-Periodic Systems Via Floquét Theory. Journal of Computational and Nonlinear Dynamics, 13, 021008. 10.1115/1.4037797.

Sprott, J.C. (2010) Elegant Chaos: Algebraically Simple Chaotic Flows. World Scientific, Singapore.

Uricchio, L.H., Daws, S.C., Spear, E.R. & Mordecai, E.A. (2019) Priority effects and nonhierarchical competition shape species composition in a complex grassland community. The American Naturalist, 193, 213–226. 10.1086/701434.

Ważewski, T. (1950) Systèmes des équations et des inégalités différentielles ordinaires aux deuxièmes membres monotones et leurs applications. Annales de la Société Polonaise de Mathématique, 23, 112–166.

